# NEURD offers automated proofreading and feature extraction for connectomics

**DOI:** 10.1101/2023.03.14.532674

**Authors:** Brendan Celii, Stelios Papadopoulos, Zhuokun Ding, Paul G. Fahey, Eric Wang, Christos Papadopoulos, Alexander B. Kunin, Saumil Patel, J. Alexander Bae, Agnes L. Bodor, Derrick Brittain, JoAnn Buchanan, Daniel J. Bumbarger, Manuel A. Castro, Erick Cobos, Sven Dorkenwald, Leila Elabbady, Akhilesh Halageri, Zhen Jia, Chris Jordan, Dan Kapner, Nico Kemnitz, Sam Kinn, Kisuk Lee, Kai Li, Ran Lu, Thomas Macrina, Gayathri Mahalingam, Eric Mitchell, Shanka Subhra Mondal, Shang Mu, Barak Nehoran, Sergiy Popovych, Casey M. Schneider-Mizell, William Silversmith, Marc Takeno, Russel Torres, Nicholas L. Turner, William Wong, Jingpeng Wu, Szi-chieh Yu, Wenjing Yin, Daniel Xenes, Lindsey M. Kitchell, Patricia K. Rivlin, Victoria A. Rose, Caitlyn A. Bishop, Brock Wester, Emmanouil Froudarakis, Edgar Y. Walker, Fabian Sinz, H. Sebastian Seung, Forrest Collman, Nuno Maçarico da Costa, R. Clay Reid, Xaq Pitkow, Andreas S. Tolias, Jacob Reimer

**Affiliations:** Center for Neuroscience and Artificial Intelligence, Baylor College of Medicine, Houston, USA; Department of Neuroscience, Baylor College of Medicine, Houston, USA; Allen Institute for Brain Science, Seattle, USA; Princeton Neuroscience Institute, Princeton University, Princeton, USA; Electrical and Computer Engineering Department, Princeton University, Princeton, USA; Computer Science Department, Princeton University, Princeton, USA; Brain & Cognitive Sciences Department, Massachusetts Institute of Technology, Cambridge, USA; Department of Mathematics, Creighton University, Omaha, USA; Department of Electrical and Computer Engineering, Rice University, Houston, USA; Department of Physiology and Biophysics, University of Washington, Seattle, USA; UW Computational Neuroscience Center, University of Washington, Seattle, USA; Institute of Molecular Biology and Biotechnology, Foundation for Research and Technology Hellas, Heraklion, Greece; Institute for Bioinformatics and Medical Informatics, University Tübingen, Tübingen, Germany; Institute for Computer Science, University Göttingen, Göttingen, Germany; Research Exploratory Development Department, Johns Hopkins University Applied Physics Laboratory, Baltimore, USA

**Keywords:** EM connectomics, Neural Morphology, Automated Proofreading, Neural Annotation

## Abstract

We are now in the era of millimeter-scale electron microscopy (EM) volumes collected at nanometer resolution (Shapson-Coe et al., 2021; Consortium et al., 2021). Dense reconstruction of cellular compartments in these EM volumes has been enabled by recent advances in Machine Learning (ML) (Lee et al., 2017; Wu et al., 2021; Lu et al., 2021; Macrina et al., 2021). Automated segmentation methods produce exceptionally accurate reconstructions of cells, but post-hoc proofreading is still required to generate large connectomes free of merge and split errors. The elaborate 3-D meshes of neurons in these volumes contain detailed morphological information at multiple scales, from the diameter, shape, and branching patterns of axons and dendrites, down to the fine-scale structure of dendritic spines. However, extracting these features can require substantial effort to piece together existing tools into custom workflows. Building on existing open-source software for mesh manipulation, here we present “NEURD”, a software package that decomposes meshed neurons into compact and extensively-annotated graph representations. With these feature-rich graphs, we automate a variety of tasks such as state of the art automated proofreading of merge errors, cell classification, spine detection, axon-dendritic proximities, and other annotations. These features enable many downstream analyses of neural morphology and connectivity, making these massive and complex datasets more accessible to neuroscience researchers focused on a variety of scientific questions.

## Introduction

To understand the morphological features of individual neurons and the principles governing their connectivity, the use of large-scale electron microscopy and reconstruction of entire neural circuits is becoming increasingly routine. For example, the MICrONS Consortium published a millimeter-scale open-source dataset of mouse visual cortex (Consortium et al., 2021) (approximately 80,000 neurons and 500 million synapses; “MICrONS dataset”), and a team at Harvard published a similar reconstructed volume of human temporal lobe (Shapson-Coe et al., 2021) (approximately 15,000 neurons, 130 million synapses; “H01 dataset”). These reconstructions offer opportunities for analysis of neural morphology and synaptic connectivity at a scale that was previously inaccessible. However, effectively using these massive and complex datasets for scientific discovery requires a new ecosystem of software tools.

Here, we describe NEURD — short for “NEURal Decomposition” — a Python software package that extracts useful information from the 3-D mesh and synapse locations for each neuron, and implements workflows for automated proofreading, morphological analysis, and connectomic studies. NEURD decomposes the 3-D meshes of neurons from EM reconstructions into a richly-annotated graph representation with many pre-computed features. These graphs characterize the neuron at the level of non-branching segments in the axonal and dendritic arbor and can support powerful queries spanning spatial scales from the geometry of the neuropil to the morphology of boutons and spines.

We begin by demonstrating the utility of this framework in an automated proofreading pipeline that is highly effective at correcting merge errors using heuristic rules. We focus on merge errors because of their catastrophic effects on connectivity analyses. We next show how the pre-computed features extracted by NEURD can enable us to recapitulate and extend a variety of previous observations about neural morphology and geometry, taking advantage of the diverse feature set computed on thousands of reconstructed neurons spanning all cortical layers in these volumes. Finally, we examine the potential of the NEURD workflow to yield novel scientific insights about neural circuit connectivity, including higher-order motifs where we observe at least three nodes (neurons) and edges (synapses) in a specific graph arrangement. NEURD includes a fast workflow to identify axonal-dendritic proximities (regions where the axon of one neuron passes within a threshold distance of a postsynaptic dendrite).

Like other open-source software packages that have supported the widespread adoption of other complex data modalities such as calcium imaging (CaImAn, Suite2P; Giovannucci et al. 2019; Pachitariu et al. 2017), neuropixel recordings (KiloSort, MountainSort; Pachitariu et al. 2023; Chung et al. 2017), label-free behavioral tracking (DeepLabCut, MoSeq, SLEAP; Mathis et al. 2018; Pereira et al. 2022; Markowitz et al. 2018), and spatial transcriptomics (Giotto, Squidpy; Dries et al. 2021; Palla et al. 2022), the goal of NEURD is to make “big neuroscience data” (in this case, large-scale EM reconstructions) accessible to a larger community. As more large-scale EM reconstructions become available, tools like NEURD will become increasingly essential for exploring principles of neural organization across multiple species.

The following tables are provided in the appendix for further reference: Table 1 - cell type subclass abbreviations and excitatory/inhibitory classification glossary, Table 2 - dataset sizes and relevant Ns for all figures and statistics, and Table 3 - a comprehensive guide to the pre-computed features provided by NEURD.

## Results

### Summary of large-scale dense EM reconstructions

Data collection for the MICrONS and H01 dataset has been described in previous publications (Consortium et al., 2021; Shapson-Coe et al., 2021). The tissue preparation, slicing procedure, and imaging resolution (4-8nm x 4-8nm x 30-40nm) was roughly similar in both cases. However, the imaging and reconstruction workflows for the two datasets were very different. The MICrONS volume was collected with transmission electron microscopy (TEM) (Yin et al., 2020), while the H01 volume was collected with scanning electron microscopy (SEM) (Hayworth et al., 2014), and different reconstruction pipelines were used.(Macrina et al., 2021; Consortium et al., 2021; Shapson-Coe et al., 2021). However, all volumetric reconstructions produce similar 3-D meshes as a common data product downstream of the segmentation process. The capabilities of NEURD are focused at the level of these mesh representations, which are much more lightweight than the original EM data, but still capture rich information about the microscale anatomy of neurons that can be useful for a variety of downstream analyses, including comparative analyses of neural circuitry across species, volumes, and reconstructions.

### Preprocessing of Neuronal Meshes

EM reconstructions yield neural meshes with varying levels of completeness, and with different kinds of merge errors (Fig. 1b-e). Merge errors include multiple whole neurons connected together (Fig. 1c), and disconnected pieces of neurite (“orphan neurites”) merged onto different neural compartments (Fig. 1d). Merge errors may also include glia or pieces of blood vessels merged onto neurons (Fig. 1f). We take advantage of existing tools for mesh processing (Fabri and Rineau, 2023; Cacciola et al., 2023; Gao et al., 2023; Dorkenwald, 2022) and apply them in an initial workflow that is agnostic to the identity of the mesh object (Fig. 2, see Methods). Systematic inspection by manual proofreaders confirmed the high accuracy of the soma, axon, dendrite, glia, compartment and spine annotations generated during the mesh processing workflow (Fig. 6, Fig. 7).

**Fig. 1.**
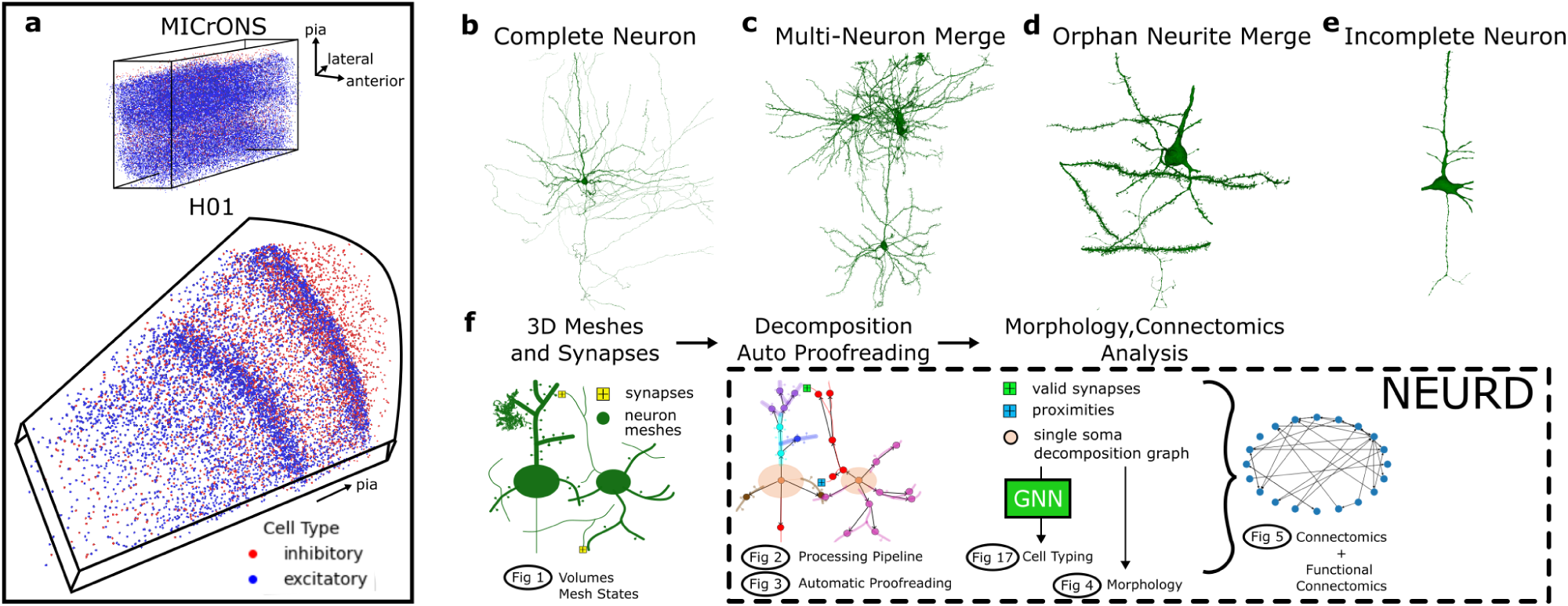
Working with neuronal meshes from large-volume EM segmentations. **a** The MICrONS Minnie65 volume is an approximately 1300 × 820 × 520 µm^3^ rectangular volume from mouse visual cortex, while H01 is a wedge-shaped volume from human temporal cortex with a longest dimension of 3mm, a width of 2mm, and a thickness of 150*µ*m. Panels **b-e** illustrate the range of accuracy across neural reconstructions in both the MICrONS and H01 volumes. **b** Example of a nearly complete (manually-proofread) single neuron, and **c** a mesh containing two merged neurons from the MICrONS volume. **d** Example of an orphan merge error with a piece of dendrite incorrectly merged onto a neuron mesh, and **e** an incompletely-reconstructed neuron from the H01 volume. Panel **f** provides an overview of the NEURD workflow with numbers representing the main figures in the paper.

**Fig. 2.**
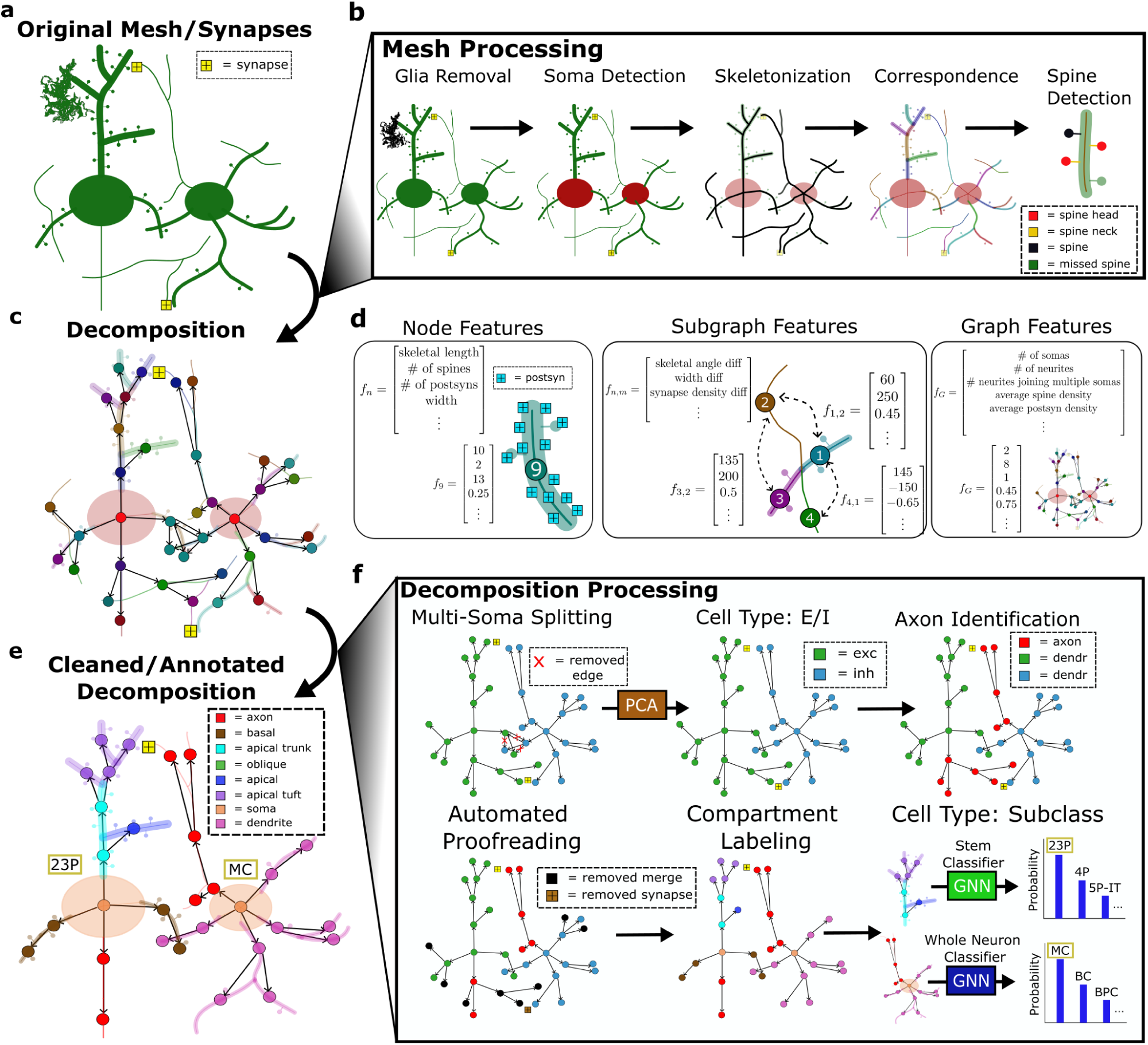
Decomposition, feature extraction, and graph annotation. **a** The input data (mesh and synapses) required for the NEURD workflow. **b** The reconstructed meshes are pre-processed through a number of steps including decimation, glia and nuclei removal, soma detection, and skeletonization. Mesh features are projected back onto the skeleton and spines are detected. **c** Decomposition graph object composed of two neurons merged together. The decomposition compresses the skeleton, mesh and synapse annotations of a non branching segment into a single node in a graph, with directed edges to the downstream segments connected at a branch point. The soma is the singular root node of this tree. **d** NEURD automates computation of features at multiple levels. Node (non-branching segment)-level features include basic mesh characteristics (e.g. diameter of the neural process or number of synapses per skeletal length). Subgraph features capture relationships between adjacent nodes like branching angle or width differences. Graph features capture characteristics of the entire neuron and are computed by weighted average or sum of node features, or by counting subgraph motifs. **e** The final product is a cleaned and annotated decomposition object with a single soma that can be fed into a variety of downstream analyses. **f** NEURD supports a variety of operations and manipulations on the decomposition objects. Multi-soma splitting is performed with heuristic rules. The entire decomposition graph is classified as excitatory or inhibitory and one subgraph is identified as the axon. Automated proofreading is performed to remove probable merge errors (see Fig. 3). A set of heuristic rules are implemented to label neural compartments, followed by a finer scale cell type classification using GNNs (see Fig. 17).

### Graph Decomposition

We decompose skeletons of axonal and dendritic processes into a directed tree graph (NetworkX object in Python Hagberg et al. 2008, and we provide a step-by-step online tutorials on how to export these as SWC files). In these graphs the root node is the soma and the other nodes are individual non-branching segments. Edges project downstream away from the soma, and subgraphs downstream of the soma are a stem. Multiple soma nodes are split apart if more than one soma is detected, and any cycles in the graph are broken during the decomposition process (see Methods, Fig.2f). Previous work has emphasized the utility of this kind of graph representation of each neuron, which facilitates flexible queries and analyses of features and annotations at different scales (Pastor et al., 2021; Schneider-Mizell et al., 2016).

NEURD computes a large number of features at the branch (node), stem (subgraph), or whole neuron (graph) level (Fig. 2c,d). These multi-scale features make it straightforward to translate neuroscience domain knowledge into neuron or compartment-level operations and queries. The most important context for this translation is automatic proofreading (Fig. 3), and NEURD also includes workflows for common tasks such as cell-type classification (Fig. 17), morphological analysis (Fig. 4), and connectivity analysis (Fig. 5).

**Fig. 3.**
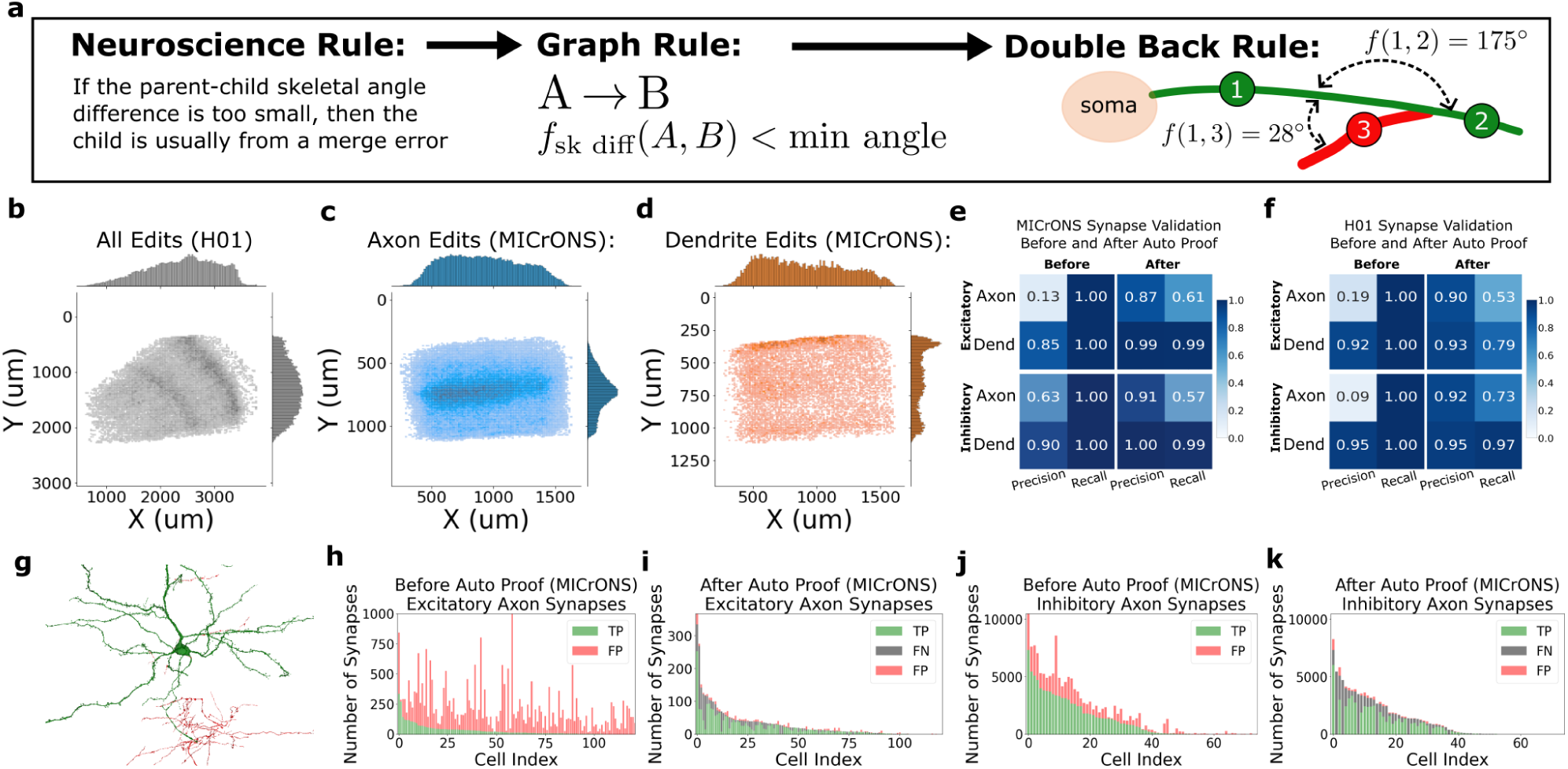
NEURD graph decomposition enables automated proofreading. **a** Implementing domain knowledge as subgraph rules to automatically identify and remove merge errors (see Fig. 8) **b** Laminar distribution of merge errors the H01 dataset. The inhomogeneity of errors across different layers may be due to differences in neuropil density. Pial surface to the right and slightly up (see Fig. 12 for more details) **c** In the MICrONS dataset, an increased frequency of axon edits is observed in layer 5 of cortex. Pial surface is up. **d** Dendritic errors in the MICrONS dataset are increased near the top layers of the volume, where fine excitatory apical tufts lead to more frequent merges (see Fig. 11 for more details). **e,f** Validation of MICrONS and H01 neurons quantified by synapse precision and recall compared to manual proofreading (“ground truth”). “Before” describes the accuracy of the raw segmentation prior to any proofreading. Note the high precision of dendrites in both volumes, even prior to automated proofreading. The substantial increases in precision “After” automated proofreading (especially for axons) indicates that the cleaned neurons have good fidelity approaching the accuracy of manual merge error removal. The reduction in “After” recall indicates that we are losing some valid synapses in the automatic proofreading process (mostly on the axons) but still retaining the majority of correct synapses (see also Fig. 14 restricted to single somas). **g**. Example excitatory neuron from the MICrONS dataset in the 75th percentile of merge error skeletal length; identified merge errors in red. **h-k** Number of true positive (TP; green) and false positive (FP; red) axonal synapses from individual excitatory or inhibitory neurons in the validation set before and after automated proofreading, illustrating the large number of false positive (red) synapses in the raw segmentation that are removed by automated proofreading (see Fig. 13 and Fig. 14 for more details on the MICrONS dataset and Fig. 15 for similar validation on the H01 dataset).

**Fig. 4.**
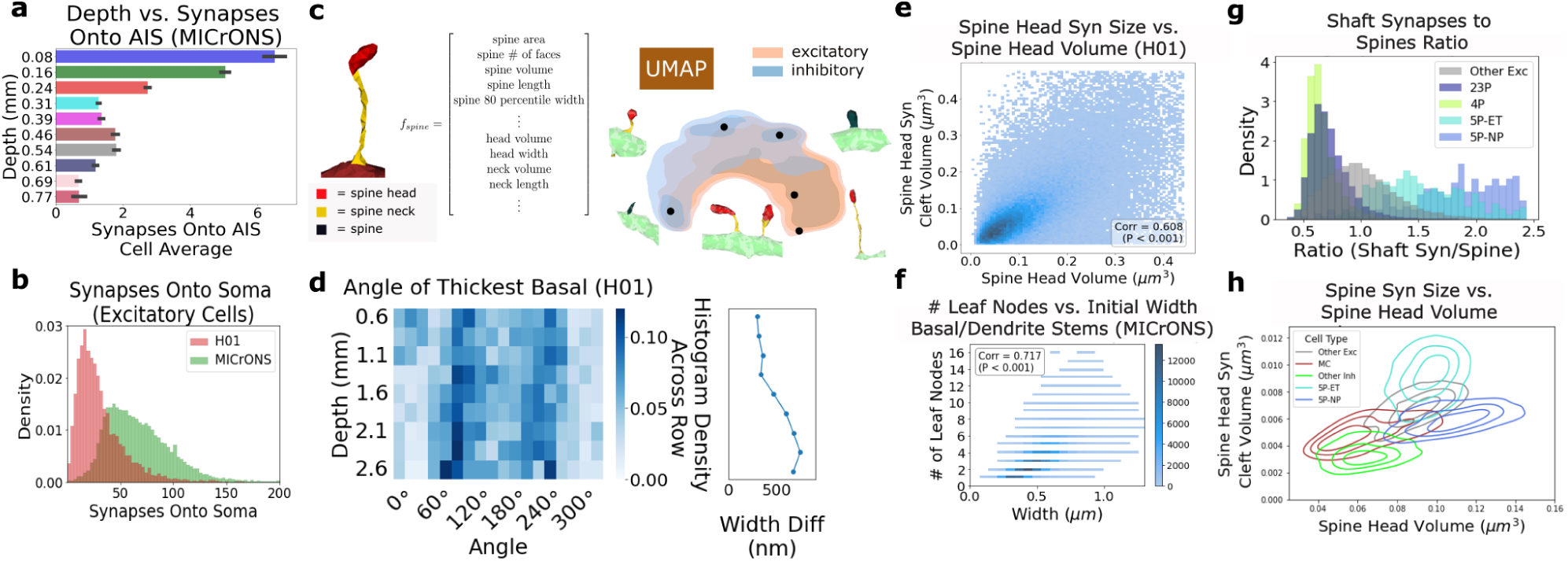
Morphological analysis enabled by NEURD feature extraction. **a** Average number of synapses onto the axon initial segment (AIS) of cells at different laminar depths (mean +/- std) for MICrONS **b** Distribution of the number of soma synapses per cell. As expected, neurons in the MICrONS volume have more identified synapses onto their soma, despite the smaller surface area compared to human somas (see Fig. 21 for more AIS and soma synapse results). **c** An example spine segmentation with the features extracted for each spine submesh followed by a kernel density estimation of the UMAP embedding of these features for spines sampled from the MICrONS dataset (see Fig. 24 for more details). **d** Histograms showing the distribution of the mean skeletal angle of the thickest basal stem as a function of volume depth (see Fig. 22 for more details). **e** Spine head synapse size and spine head volume joint distribution for the H01 dataset (see Fig. 25 for more details). **f** Histogram for all the non-apical dendritic stems of every neuron in the MICrONS volume comparing the initial width of the stem to the number of leaf nodes, showing a positive correlation (see Fig. 23 for more details). **g** The ratio of non-spine synapses to spine counts varies across cell types. **h** Distribution of spine characteristics for different cell types, comparing each spine head’s volume and the size of the largest synapse on that spine head, where outlines indicate the quartile boundaries for each distribution (see Fig. 20 for more cell type distributions).

**Fig. 5.**
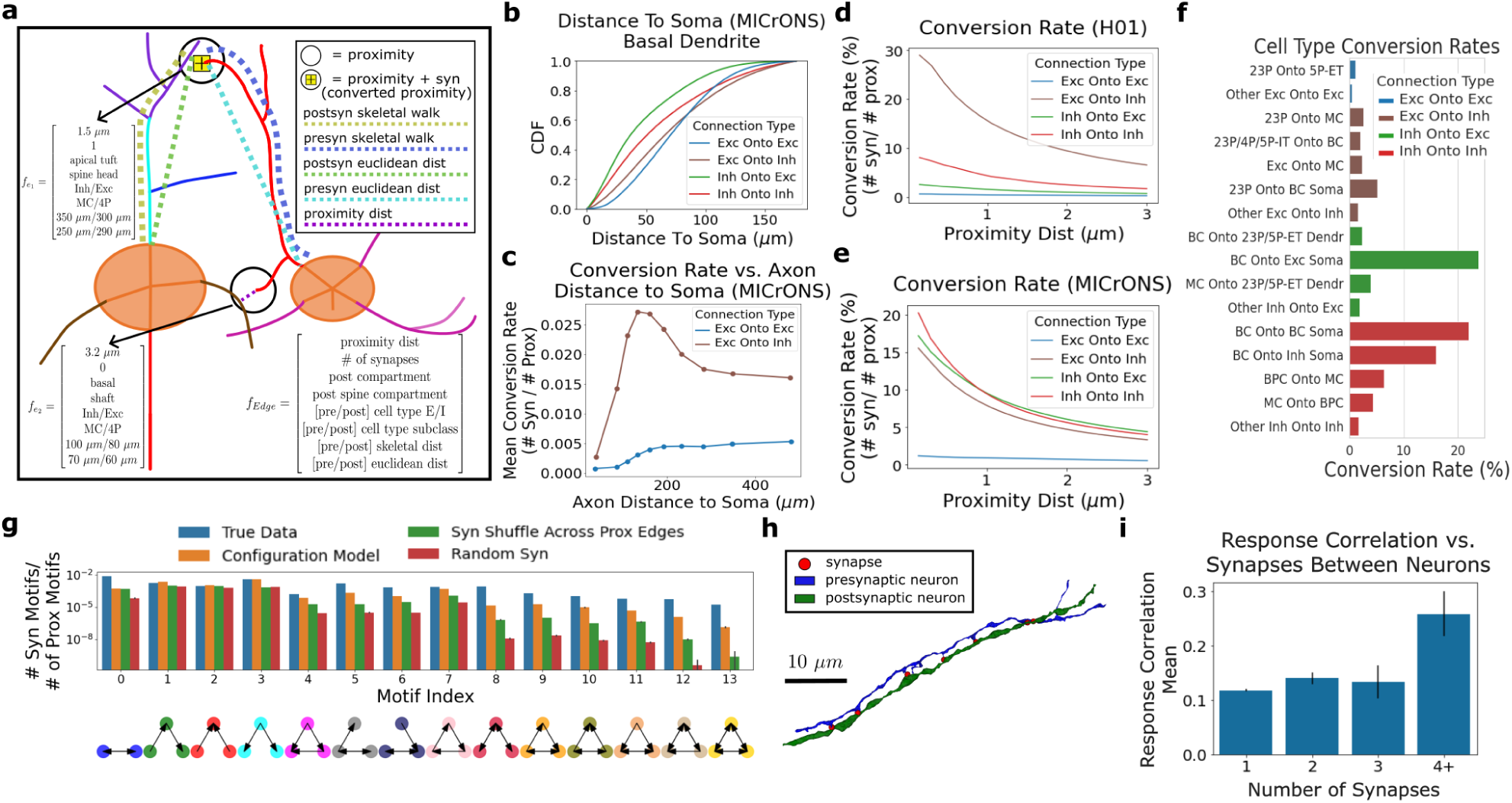
Connectivity analysis enabled by NEURD. **a** Schematic illustrating two proximities between a pair of neurons (axon passing within 5 *µ*m radius of dendrite). Only one proximity has a synapse, thus the “conversion rate” is 50%. Each proximity creates an edge in a proximity graph between the two neurons, and each proximity is annotated with a variety of pre-computed features such as distance to the soma, etc. **b** Cumulative density function (CDF) of the postsynaptic skeletal walks for different connection types (apical and soma synapses excluded), demonstrating excitatory inputs occur further along the dendrite from the soma (see Fig. 28 for more details). **c** Mean conversion rate as a function of distance along the axon (see Fig. 28 for more details). **d,e** Conversion rates (synapses / proximities) for different excitatory and inhibitory combinations. The x-axis represents the maximum distance that is considered a “proximity”. **f** Conversion rates for different cell type subclasses and compartments in the MICrONS dataset are largely consistent with previous studies (cell type labels generated from GNN classifier, see Table 1 for glossary). See Fig. 29 for more conversion rates. **g** The frequency of reciprocal connections or edge-dense three-node motifs was enriched compared to null distributions where synaptic degree distribution is held the same but edges are shuffled (orange), where the synaptic edges are shuffled only between neurons with an existing proximity edge (green), or where synapses are randomly shuffled between neurons regardless of proximity (red); 250 random graph samples for each null distribution comparison (see Fig. 30 for more details and inhibitory/excitatory-only graphs). **h** Example multi-synaptic connection (n=7 synapses) from an excitatory to inhibitory neuron in the H01 dataset. **i** Distribution of response correlation between pairs of functionally matched excitatory neurons in the MICrONS dataset. Response correlation is significantly larger for pairs of neurons with 4 or more synapses connecting them (n=11 pairs) compared to those with 1, 2, or 3 synapses; n=5350, 280, 34 pairs respectively (see Fig. 32 for more details).

### Automated Proofreading

Node, graph, and subgraph features can be queried to identify patterns of features that are commonly found at merge errors in the reconstruction (for example sharp discontinuities in width between adjacent dendritic nodes, or axonal branches that “double-back” sharply towards the soma). Once the error location is identified, all nodes downstream of the error are stripped from the mesh and returned to the “sea of unconnected fragments” in the volume, excluding them from any subsequent analysis. The NEURD proofreading workflow is easily extensible, and the user can define which graph filters to apply in which order. For any error correction, the rule and relevant parameters that determined the edit are stored for subsequent evaluation and use, potentially providing a rich set of training data for machine learning approaches.

To illustrate this process, we provide a small set of heuristic proofreading rules implemented as graph filters (Fig. 3a, Fig. 8; see Methods) that yielded good performance on merge error correction in both volumes, but especially in the MICrONS dataset. Manual validation of these rules was performed in the context of standard proofreading and multi-soma splitting using the NeuVue Proofreading Platform (Xenes et al., 2022) by the proofreading team at Johns Hopkins University Applied Physics Laboratory (APL). We provided APL with suggested error locations in the MICrONS volume, and experienced proofreaders evaluated each proposed split for accuracy (Fig. 9a-c). This validation set included multi-soma splitting, axon-on-axon, and axon-on-dendrite merge errors, and enabled us to measure both the accuracy of these proofreading rules and the speed benefits of a semi-supervised approach compared to a fully-manual effort. We were also able to optimize these rules based on proofreader feedback, and we identified specific rules and parameter thresholds could be applied with high confidence to correct merge errors without human intervention. Validation of this “high confidence” subset of axon-on-axon merges and axon-on-dendrite merges yielded a 99% and 95% agreement between the algorithm’s split operations and those performed by a human proofreader (Fig. 9e). We applied nearly 150,000 of these high-confidence automatic edits in bulk to the publicly-available MICrONS segmentation volume. Using NEURD suggestions in a semi-supervised manner to guide the challenging process of splitting multi-soma segments increased the speed of this process more than three-fold compared to other methods (Fig. 9d, see Methods).

In the MICrONS dataset, we identified hundreds of thousands of merge errors corresponding to dozens of meters of incorrectly-merged axons and dendrites within the 1mm^3^ volume (Fig. 10). Corrections in the H01 dataset were an order of magnitude smaller due to fewer cells and the less complete initial reconstructions in that volume, but were still substantial (Fig. 10). Merge errors were more prevalent in some regions due to sectioning artifacts (Fig. 11g-l, Fig. 12h-n), or due to intrinsic differences in the morphology of neurons across layers (Fig. 3b-d, Fig. 11a-f, Fig. 12a-g,) For example high- and low-degree axon edits in the MICrONS dataset were frequently made in upper layer 5, potentially due to higher quantities of inhibitory neuropil, while dendritic double-back and width jump errors were more frequently located near the top layers of the volume, due to merges between fine distal apical tufts of excitatory cells (Fig. 11, Fig. 12).

We compared the outcome of automatic proofreading (all edits, not just the high confidence subset) to manual proofreading “ground truth” on a test set of cells in the MICrONS (n=122 excitatory and n=75 inhibitory) and H01 (n=49 excitatory and n=18 inhibitory) volumes (see Fig. 35 for manually-proofread cell locations).

The precision of the synaptic data, i.e. the number of actually-true synapses that were labeled as true divided by the total number of synapses labeled as true, was substantially higher after proofreading (e.g., 0.87 after compared to 0.13 before for MICrONS excitatory axons, a greater than sixfold increase). For the same axons, this increase in precision was achieved with only a 40% reduction in recall (number of correctly-identified synapses divided by the number of true synapses). Low precision can be catastrophic for downstream analyses, while low recall can support many analyses based on the assumption that the observed synapses are a representative sample of the full distribution (precision and recall are summarized in Fig. 3e,f, Fig. 13, Fig. 14, and Fig. 15 ). The precision and recall of our automated method is captured visually in the plots in Fig. 3h-k. Comparing “before” and “after” proofreading performed with NEURD, red false positives are almost entirely removed, at the cost of a thin margin of false negatives (gray) cutting into the green true positives. For example, for the axons of the the 122 excitatory neurons in the MICrONS dataset, NEURD correctly removed 24,430 false synapses and incorrectly removed 1,420 true synapses, leaving 2,216 true synapses and 324 false synapses. Full numbers are available in Appendix Table 2.

Because our automated proofreading procedure only removes data, recall is measured based on the true synapses in the automatic segmentation (after merge errors were removed manually), and does not include synapses from any manual extensions. Recall was especially high for dendrites ( 99% for MICrONS single-soma neurons, Fig. 14), reflecting the high performance of the initial segmentations. Overall recall was lower for axons (approximately 60% for both excitatory and inhibitory cells in the MICrONS dataset), indicating that NEURD incorrectly removed a larger number of axonal segments compared to dendrites. Performance on the H01 dataset was also reduced compared to MICrONS because the less-extensive reconstruction was associated with fewer merge errors overall. Extensive, sometimes centimeter-scale arbors remained after removing merge errors (Fig. 3g, Fig. 16). In summary, both from the perspective of synapses (Fig. 3h-k, Fig. 13i-p, Fig. 15i-p) and skeletons (Fig. 13e-h, Fig. 15e-h), our automated proofreading approach can be applied at scale to remove merge errors with accuracy approaching manually-cleaned cells (Fig. 16).

### Cell-type classification

Densely-reconstructed EM volumes hold great promise for understanding the connectivity between different neural subtypes (Schneider-Mizell et al., 2023, 2021; Dorkenwald et al., 2022a; Weis et al., 2022; Dorkenwald et al., 2022b; Peters and Feldman, 1976; Martin and Whitteridge, 1984). Because EM provides limited access to genetic markers, cell types must be identified by morphological features or connectivity (if sufficient proofreading is performed). The relationship with molecularly-defined cell classes can sometimes be inferred from extensive previous work relating morphological features to transcriptomic classes (Scala et al., 2021; Peng et al., 2021; Gouwens et al., 2020), including from our consortium (Gamlin et al., 2023). Previous studies have demonstrated that rich information enabling cell-type classification is available even in local nuclear and peri-somatic features (Elabbady et al., 2022; Al-Thelaya et al., 2021), small segments of neural processes (Dorkenwald et al., 2022b), different resolution views of single nuclei segments (Zinchenko et al., 2023), and the shape of postsynaptic regions (Seshamani et al., 2020). NEURD provides an additional rich and interpretable feature set at the level of non-branching segments that can be used for accurate cell-type classification via a number of different approaches, as we describe below.

As expected (Azouz et al., 1997), we found that a logistic regression model trained on just two spine and synapse features separates excitatory and inhibitory cells with high accuracy, using the same parameters for classification across both the MICrONS and H01 dataset (Fig. 17a,b; MICrONS n=3,985 excitatory and n=897 inhibitory; H01 n=5,800 excitatory and n=1,755 inhibitory). To test whether NEURD graph objects could be used to distinguish even finer cell types, we turned to Graph Convolutional Networks (GCN) (Fig. 17c-f). We trained a simple GCN on the dendritic subgraph of a variety of hand-labeled cell types in the MICrONS volume (n=873 total cells), which represent a relatively complete set of cell type classes for this volume and are more thoroughly described in (Schneider-Mizell et al., 2023; Elabbady et al., 2022). We focused on the dendrites in this volume because of their high recall from the initial segmentation and the high precision after automated proofreading (Fig. 3f). Most of the embedding space was covered by the labeled dataset (Fig. 17c), and cells outside the labeled dataset had soma centroids at expected laminar depths (Fig. 17d); even though no coordinate features were used in the GCN classifier.

We evaluated cell type classification performance on a held-out test set using a GCN with access to the entire dendritic graph (n=178 test neurons; mean class accuracy = 0.82; class accuracy: 23P (0.94), 4P (0.69), 5P-IT (0.60), 5P-NP (1.00), 5P-ET (0.86), 6P-CT (0.58), 6P-IT (0.62), BC (0.85), BPC (0.89), MC (1.00), NGC (1.00); Fig. 17e, also see actual counts for training, validation, and test in Fig. 19a-c). We also evaluated the classification performance using only disconnected dendritic stems (n=1023 test stems; mean class accuracy = 0.66; class accuracy: 23P (0.73), 4P (0.78), 5P-IT (0.33), 5P-NP (0.78), 5P-ET (0.78), 6P-CT (0.56), 6P-IT (0.49), BC (0.80), BPC (0.70), MC (0.79), NGC (0.50); Fig. 17f, counts for training, validation, and test in Fig. 19d-f). The information present in disconnected individual dendritic stems (branching segments connected to the soma) is thus sufficient to perform fine cell-type classification nearly as well as graphs representing entire neurons, consistent with previous literature classifying cells based on more local features (Elabbady et al., 2022; Al-Thelaya et al., 2021; Dorkenwald et al., 2022b). See Table 1 for the cell-type abbreviation glossary. Because the classifier is a deep learning model, the output from the final softmax layer can be used as a confidence measurement, making it possible to restrict downstream analyses to high confidence cell-type labels.

### Morphological Analysis

The features extracted by NEURD - including features of different compartments (Fig. 24a), the geometry of axonal and dendritic compartments (Fig. 22a), and spine features (Fig. 24b) - provide a rich substrate for morphological analysis (Fig. 20).

In particular, extensive work has linked spine morphology to synaptic strength and stability, making them important targets for understanding plasticity and connectivity in neural circuits. A variety of methods have been developed to automate spine detection in 2-D or 3-D image data using fully-automatic (Xiao et al., 2018; Driscoll et al., 2019; Janoos et al., 2009; Shi et al., 2014; Basu et al., 2018) or semi-automatic (Benavides-Piccione et al., 2013) approaches. NEURD offers an accurate spine detection workflow that achieves high accuracy with a fully-automated mesh segmentation approach. Precision and recall for spines with a skeletal length larger than 700 nm was 90% or higher (Fig. 6). In addition, NEURD segments the spine head from the neck (when possible), and computes statistics about the individual head and neck submeshes, creating a feature-rich dataset for testing hypotheses about spine morphology that can then be conditioned on postsynaptic compartment type or the cell type of the pre- or postsynaptic cell. As expected, the spine head volume and synaptic density volume were the only strongly correlated spine features (Fig. 25; Harris and Stevens 1989; Arellano et al. 2007). The spatial distribution of UMAP embeddings (two-dimensional projection) for feature vectors of spines sampled from the MICrONS and H01 dataset showed a similar structure, with spines that share similar features embedded in similar locations and a somewhat consistent embedding pattern for inhibitory and excitatory spines in the two volumes. This similarity suggests that H01 and MICrONS spines sample from a similar landscape of diverse spine shapes. (Fig. 24c,d), consistent with previous work examining the distribution of non-parametric representations of post-synaptic shapes across diverse neural subtypes (Seshamani et al., 2020).

We attempted to replicate and extend several other findings observed in previous studies of the MICrONS and H01 datasets regarding the sub-cellular targeting of synaptic inputs. First, we computed the number of synapses onto the axon initial segment (AIS) of neurons at different depths. Replicating a previous report, in the MICrONS volume, superficial L2/3 pyramidal cells received the largest number of AIS synapses, with up to 2-3 times the innervation of the lower cortical layers (Fig. 4a; Schneider-Mizell et al. 2021; Wang et al. 2019; Inan et al. 2013). However, in the H01 dataset, this laminar inhomogeneity in AIS synapses was much less prominent, with more similar numbers of AIS inputs observed across all depths (Fig. 21h). Additionally, like AIS synapses, we found a striking difference in the distribution of somatic synapses across depth between the MICrONS and H01 dataset. (Fig. 21e,f). Lastly, the overall frequency of somatic synapses were also distinct across the two volumes, consistent with previous literature describing fewer somatic synapses in the human compared to mouse (Fig. 4b; Wildenberg et al. 2021); however, we found the opposite trend for the AIS, with fewer AIS synapses in the mouse volume compared to the human volume (Fig. 21i).

In H01, deep layer pyramidal cells were previously observed to have a strong bias in the radial angle of their thickest basal dendrite (Shapson-Coe et al., 2021). We examined the MICrONS volume and did not observe a strong bias in thickest basal, even in deep layers (Fig. 22b). Then, looking at H01, we were able to replicate the pattern of thickest basal dendrite direction preferences in deeper layers (Fig. 22c). However, we also found that this pattern appeared to continue into more superficial layers, extending the previous finding. In deep layers the difference between the thickest and second-thickest dendrite was nearly twice what it was in more superficial layers, making this effect more salient.

The diversity of pre-computed features offered by NEURD enabled us to identify several interesting morphological features that differ across cell types, including many that have been reported previously in other studies. For example, the spindly, non-branching basal dendrites of NP cells (Schneider-Mizell et al., 2023; Weis et al., 2022) are clearly distinct from extensively-branching basal dendrites of L2/3 pyramidal cells (Fig. 20h), and neurogliaform cells are the most highly-branched neurons with the largest number of leaf nodes (Fig. 23a). Across all neurons, dendritic stems with larger numbers of leaf nodes had a larger initial dendritic diameter at their connection to the soma (Fig. 23b-c), potentially reflecting developmental or metabolic constraints.

Synapses onto the dendritic shaft and synapses onto dendritic spines roughly correspond to inhibitory versus excitatory inputs (Ribak et al., 1981; DeFelipe and Fariñas, 1992; Kwon et al., 2019). In a histogram of shaft to spine synapses, NP cells were again located at the higher end of the distribution, while L4 and L2/3 pyramidal cells had the lowest shaft-synapse to spine-synapse ratio (Fig. 4g), suggesting they receive a relatively larger fraction of excitatory (compared to inhibitory) input. Because NEURD also automatically segments both soma meshes and spine heads and necks, this enables comparison across cell types of features like soma volume and somatic synapses (Fig. 20b,f), spine neck length (Fig. 20i), spine density (Fig. 20a), and the relationship between spine synapse size and spine head volume as in Fig. 4h and Fig. 25.

### Connectivity and Proximities

Next, we examined the connectivity graph in the MICrONS and H01 datasets after automatic proofreading. As expected, proofreading substantially reduced the mean in-degree (number of incoming synapses onto a neuron) and out-degree (number of projecting synapses from a neuron) across both volumes due to the removal of merge errors, resulting in a sparsely-sampled but high-fidelity graph (Fig. 3e,f). A variety of connectivity statistics including number of nodes and edges, mean in and out degrees, and mean shortest path between pairs of neurons along excitatory and inhibitory nodes are provided in Fig. 27.

To facilitate the analysis of synapse specificity in sparse connectomes, we implemented a fast workflow for identifying axonal-dendrite proximities, regions where the axon of one neuron passes within a threshold distance (here 5 *µm*) of the dendrite of another neuron (See Methods, Fig. 5a). Previous studies have computed proximities from skeletons of simulated models (Udvary et al., 2022), or manually traced data (Mishchenko et al., 2010; Kasthuri et al., 2015), with a similar logic. Proximities are necessary but not sufficient for the formation of a synapse (Peters and Feldman, 1976; Brown and Hestrin, 2009; Mishchenko et al., 2010; Costa and Martin, 2011), and so the “proximity graph” can serve as a valuable null distribution for comparing potential connectivity with synaptic connectivity between neurons: Instead of looking at synapse counts between cells which are dependent on the geometry and completeness of the neuropil, proximities make it possible to calculate “conversion rates” - the fraction of proximities which resulted in actual synaptic connections. NEURD also provides functions to compute “presyn skeletal walk” - the distance from a synapse to the soma of the presynaptic neuron along the axon, and “postsyn skeletal walk” - the distance from synapse to soma along the postsynaptic dendrite. Combined with cell typing, compartment labeling and spine annotation, these features enable powerful analyses of neural connectivity conditioned on the cellular identity and subcellular location of synapses on both pre- and post-synaptic partners (Figures 28, 31).

Conversion rates between neural subtypes in the MICrONS dataset replicated previous results from connectivity measured via slice multi-patching and EM reconstructions, especially the prolific connectivity of basket cells onto both excitatory and inhibitory somas (Fig. 5f; Jiang et al. 2015; Schneider-Mizell et al. 2023; Lee et al. 2013; Freund and Katona 2007) and inhibitory-inhibitory relationships including BC inhibiting other BC, MC avoiding inhibiting other MC, and BPC preferentially inhibiting MC (Fig.29; Pfeffer et al. 2013; Jiang et al. 2015; Lee et al. 2013; Schneider-Mizell et al. 2023, see Table 1 for the cell-type abbreviation glossary).

The subcellular targeting of different inputs is apparent in plots of the “postsyn skeletal walk” distance to the soma for synapses arriving at the basal dendrite. As has been previously described (Ribak et al., 1981; Hwang et al., 2021; Megıas et al., 2001), inhibitory-onto-excitatory synapses tend to be found closer to the somatic compartment than excitatory-onto-excitatory synapses (Fig. 5b, Fig. 28g). At an even smaller scale, with the spine head, spine neck, or shaft classification propagated to synapses, we can study how excitatory and inhibitory inputs to spines display different scaling relationships between synapse size and spine head volume (Fig. 26). We also show, as expected, that excitatory and inhibitory cells differ in the number and relative sizes of synapses on their target spine heads (Fig. 26; Parnavelas et al. 1977; Megıas et al. 2001).

Conversion rates for excitatory-to-excitatory proximities were low in both H01 and MICrONS, consistent with previous findings of sparse pyramidal cell connectivity in the cortex (Fig. 5d,e; Campagnola et al. 2022; Jiang et al. 2015; Kasthuri et al. 2015; Mishchenko et al. 2010). However, conversion rates were substantially higher for excitatory-to-inhibitory proximities (Fig. 5d,e), especially in H01, and especially for proximity distances less than 2 microns (unlike excitatory synapses onto excitatory cells, where spines presumably reduce the dependence on distance). Combining the presynaptic (axonal) skeletal walk features and proximity analyses revealed an interesting similarity in excitatory-onto-inhibitory connectivity between the MICrONS and H01 datasets, with conversion rates peaking in the more proximal axon a few hundred microns from the soma (Fig. 5c, Fig. 28e; Bock et al. 2011; Bopp et al. 2014; Schmidt et al. 2017; Dorkenwald et al. 2022b), a pattern that could reflect a lateral inhibition motif. Conversion rates were also higher above (more superficial to) the presynaptic soma than below (deeper than) the presynaptic soma for excitatory-onto-inhibitory connections in both volumes (Fig. 28b,c).

Large-volume EM connectomics offer tremendous potential opportunities to examine higher-order motifs on a large scale. We found that more densely-connected triangle motifs were enriched in the MICrONS volume compared to several controls (Fig. 5g, and Fig. 30 for excitatory and inhibitory subgraphs). The over-abundance of densely-connected triangle motifs that we observed is consistent with previous findings suggesting that this higher-order organization is enriched in the cortex (Song et al., 2005; Perin et al., 2011; Milo et al., 2002; Udvary et al., 2022; Schneider-Mizell et al., 2021). A similar pattern was observed in the H01 dataset, consistent with previous modeling of connections and proximities there (Udvary et al., 2022). However, in the H01 volume several of the three-node motifs with larger numbers of connected edges were missing due to the less complete reconstruction (Fig. 30).

### Functional Connectomics

A key advantage of the MICrONS dataset is the functional characterization of matched neurons in vivo prior to EM data collection. The relationship between function and synaptic connectivity is covered in detail in a separate paper (Ding et al., 2023), which includes analysis using NEURD’s automatically-proofread data. Here, we wanted to provide an illustration of how automated proofreading can enable a specific functional connectomics analysis that would be otherwise very challenging. We identified pairs of excitatory neurons connected by one, two, three, or four or more synapses. Querying for these rare high-degree connections between pyramidal cells was only possible after automated proofreading, since approximately 97% of the original 10,000+ pairs with four or more connections were identified as merge errors during automatic proofreading. Connections were further restricted to synapses onto postsynaptic spines in order to guard against possible missed merge errors where an inhibitory axon segment might still be merged to an excitatory neuron. Examples of these multi-synaptic connections have been highlighted in the H01 data set (Fig. 5h), and rare examples can also be found in the MICrONS data set (Fig. 32b; see also Chicurel and Harris 1992). We were able to identify n=11 of these pairs in exclusively-automatically proofread neurons (no manual proofreading) where both neurons also had been characterized functionally (Fig. 32c). The average response correlation was calculated for each group of pairs (see Methods and Ding et al. 2023; Wang and Tolias 2023). We found that neurons with four or more synapses had significantly higher response correlations (0.259 +- 0.042 (n=11)) to visual stimuli than neurons with < 4 synapses ( syn = 1: 0.118 +- 0.002 (n=5350), syn = 2: 0.140 +- 0.011 (n=280), syn = 3: 0.133 +- 0.030 (n=34)) connecting them (Fig. 5i), consistent with a Hebbian “fire together/wire together” rule governing high-degree connectivity in the cortex, and this same pattern was also observed for n=12 pairs of manually-proofread neurons. For autoproofread neurons, two-sample Kolmogorov-Smirnov test and t-test for comparing each multi-synaptic group’s null likelihood of being drawn from the same distribution as the one-synapse group: 2 synapses KS test statistic = 0.068 and p = 0.17, t-test p<0.03, 3 synapses KS test statistic = 0.139 and p=0.49, t-test p = 0.60, 4 synapses KS test statistic = 0.508 and p < 10^−2^, t-test p<10^−2^ (and still significant after the Bonferroni correction for multiple comparisons with a significance threshold of p<0.02).

## Discussion

NEURD is an end-to-end automated pipeline capable of cleaning and annotating 3-D meshes from large electron microscopy volumes and pre-computing a rich set of morphological and connectomic features that are ready for many kinds of downstream analyses. Building on existing mesh software packages, NEURD adds a suite of neuroscience-specific mesh functions for soma identification, spine detection, and spine segmentation that are applicable across multiple data sets, as well as workflows for skeletonization and mesh correspondence that complement existing tools. We demonstrate the utility of this integrated framework for morphological (Figures 4, 20, 21, 22, 23, 24, 25), and connectomic analyses (Figures 5, 26, 27, 28, 29, 30 , 32), most of which involved queries against pre-computed features, as well as demonstrating how these features can be combined to ask new questions (Figures 4h, 21i, 21j, 28d,28e, 32d, 26g, 26h). The set of features generated by NEURD is easily extensible, and our hope is that NEURD will make these daunting datasets more accessible for a larger group of researchers.

Several previous studies have proposed post-hoc methods for automated proofreading including merge and split error detection and correction. Some of these methods make use of Convolutional Neural Networks (CNNs) ( either supervised Zung et al. 2017; Gonda et al. 2021 or unsupervised Rolnick et al. 2017), reinforcement learning methods (Nguyen et al., 2022), or other machine learning approaches (Schubert et al., 2019; Schmidt et al., 2022; Berman et al., 2022). Others make use of heuristic rules applied to neural skeletons (Meirovitch et al., 2016; Sicat et al., 2013), and at least one approach uses both skeleton heuristics and CNNs (Matejek et al., 2019). Compared to automated methods applied directly to the EM segmentation that may be heavily memory-intensive, NEURD benefits from the lightweight features computed from mesh representations, allowing analysis to scale to larger volumes more easily. Methods based on the EM segmentation have the advantage that intracellular features can be used to aid proofreading, and these methods could potentially still be employed upstream of NEURD. A key strength of the coarse-scale graph with pre-computed annotations is the flexible querying across multiple scales. For instance, distinguishing whether a thin, aspiny projection from a dendrite is the true axon or a merge error might require both local features and also additional information about the neuron type, the distance from the soma, and the spine density of the parent branch.

Our present implementation does not address some types of errors in automated segmentation. For example, it cannot presently handle merge errors that result in a co-linear segment of skeleton that we interpret as a single non-branching segment. Second, because NEURD’s present implementation is focused on removing false merges, it is unable to fix incomplete neural processes (Fig.13c). Motivated by the low recall of many neurons in these volumes, we plan to try using NEURD to perform automated extensions in the future by “over-merging” candidate segments at a truncation point, and then correcting the resulting “merge error” to choose the best candidate for extension.

We performed extensive validation of our automated proofreading approach to determine the precision and recall of our error correction, but as a general disclaimer it is important to note that some of the results we have presented here might change if the same neurons were manually proofread and fully extended. For any particular scientific question, researchers using these volumes will need to weigh the relative importance of precision, recall, and the number of neurons that it is feasible to proofread using manual or automated methods. Finally, our results currently only present findings from two mammalian volumes, but there is nothing in principle preventing the application of NEURD to large EM volumes from other species in the future.

Combining automated proofreading to generate a clean but incomplete graph, with proximities to serve as a null distribution is a powerful approach that can begin to reveal principles of pairwise and higher-order connectivity motifs in incomplete reconstructions. We observe a general overexpression of densely-connected triangle motifs in comparison to proximity controls and some standard null models, as previously reported in (Udvary et al., 2022; Song et al., 2005; Perin et al., 2011; Milo et al., 2002). However, the ability to expand this work to include cell type colorings of these motifs and add proximity based controls will enable investigation of more complicated motif questions, unleashing the power of these datasets for addressing questions about higher-order circuit connectivity. As additional reconstructed volumes are released spanning species and brain areas, our ability to extract scientific insights will depend critically on integrated and scalable frameworks that make neurons and networks accessible for analysis, with a rich feature space suitable for machine learning approaches to understanding these complex datasets.

## Methods

### Data Management

For simplified data management and querying of input neuron reconstruction meshes, NEURD intermediate decomposition graphs, and all derived statistics and data products, we utilized the DataJoint Python package (Yatsenko et al., 2015, 2018)

### Mesh Preprocessing

NEURD operates on 3-D meshes which are represented in a standard form as lists of vertices and faces in 3-D coordinates. A connected mesh component is a set of faces and vertices in which all faces have at least one adjacent edge to another face. Segmentation algorithms may not output a single connected component as a mesh, but instead may generate several disconnected submeshes, each of which is a subset of faces that is a connected component. NEURD is generally robust to discontinuous meshes, meshes of different resolutions, and several kinds of meshing errors.

The resolution of meshes delivered as part of the MICrONS and H01 datasets was sufficiently high that we performed an initial decimation of the mesh (reduced to 25% for MICrONS and 18% for H01) to speed up subsequent computations while retaining all the detail necessary for morphological characterization even of fine axons and spine necks. This decimation was performed using the MeshLab Quadric Edge Collapse Decimation function (Cignoni et al., 2008). Following decimation, we separated this decimated mesh into connected components. We next applied a Poisson Surface Reconstruction (Cignoni et al., 2008) to each connected component. This can be thought of as “shrink-wrapping” the mesh - it smooths discontinuities on the surface of the mesh and ensures that each connected component is “water-tight” (i.e. no gaps or missing faces). This pre-processing facilitates the subsequent decomposition steps.

### Glia, Nuclei Removal

Glia and nuclei submeshes are identified and filtered away using ambient occlusion functions (Cignoni et al., 2008) to identify regions with a high density of inside faces. Inside faces are mesh faces that are almost fully surrounded by other mesh faces. For example, glia that are merged onto neurons appear as cavities filled with a high density of mesh faces, and are distinct from the hollow reconstructions of most excitatory and inhibitory neurons. Similarly, the mesh representation of the soma surrounds the nucleus mesh and thus nuclei are almost entirely made up of inside faces. Therefore to identify glia and nuclei in the reconstructed meshes, we look for large connected components with high percentages of inside faces as candidates. To determine whether mesh faces are internal or external, we simulate an external “light source” that emits from all angles and we compute the exposure each face receives. This metric is thresholded to classify faces as either inside or outside faces, and submeshes made up mostly of inside faces are candidates for removal. We then apply additional thresholds on the candidate submeshes volume and number of faces to classify them as a glia mesh, nuclei mesh or neither. Finally, for glia we include all floating meshes within the bounding box of the submesh or within a search radius (3000 nm) of any faces of the submesh. This post-hoc mesh agglomeration serves to clean up the areas around glia segmentation, which can be very unconnected and non-standard.

### Soma detection

Soma detection is run on any segment containing at least one detected nucleus (note that nucleus detection was performed previously as part of the segmentation and annotation workflow Consortium et al. 2021; Shapson-Coe et al. 2021). To detect the soma, we first perform a temporary heavy decimation of the mesh to remove small features and facilitate detection of the large somatic compartment. We then segment this low-resolution mesh into contiguous submeshes using the CGAL mesh segmentation algorithm (Yaz and Loriot, 2023). This function not only provides the specific faces in each submesh but also an estimate of the width of the submesh as an SDF value (Shape Diameter Function, a measure of diameter at every face of the mesh surface Yaz and Loriot 2023). We then filter all the resulting submeshes for soma candidates by restricting to those within a set size (number of faces), SDF range, bounding box length and volume threshold. We restrict to candidates that are sufficiently spherical to represent the general shape of a soma, but liberal enough to account for somas that are partially reconstructed (for example at the edge of the volume). Once we identify candidate somas in the low-resolution mesh, we return to the original mesh representation (prior to starting soma detection) and apply a final size and width threshold. Given the initial restriction to segments with at least one detected nucleus, if we are not able to detect at least one soma after this process, we relax the thresholds slightly and iterate until a soma is detected or a threshold limit for number of attempts is reached.

### Decomposition

With the glia and nuclei submeshes identified, we filter those away from the original mesh, which may cause splitting into additional connected components. We identify connected components containing at least one soma submesh (note that some segmentations may contain multiple somas prior to soma splitting). Mesh fragments that are not connected to somas may be floating meshes inside the soma (which are filtered away using the same ambient occlusion methods described for glia above), or detached mesh pieces of neural processes that can be stitched to the neuron representation later. For each of the soma-containing meshes we filter away the soma submeshes and identify connected components of these meshes as stems. Any stem submesh must contain at least one set of connected adjacent edges and common vertices shared with a soma submesh (“border vertices”). We construct a connectivity graph where edges only exist between stem and soma nodes if there exists border vertices between the stem and the soma. Through the graph constructed in this manner, stems that provide paths between multiple somas can be easily identified for subsequent splitting (see below).

Each of the stem submeshes is then processed into a skeleton - a 3-D “line-segment” representation that is a set of vertices and edges. We use the Meshparty package for this initial round of skeletonization because of its efficient implementation, and because it provides both a width estimate and a correspondence between the faces of the original mesh and each vertex in the skeleton (Dorkenwald, 2022). This skeleton is then further divided into branches (non-branching subskeletons). The corresponding meshes of branches with an average width greater than a threshold are re-skeletonized with a higher-fidelity method that yields skeletons which pass through the hollow centers of the mesh to provide a better estimation of the location and curvature of the surrounding mesh. This is particularly important for some neural processes, for example wide apical trunks where a skeleton that is not centered within the mesh could be displaced nearly a *µm* from the actual center of the trunk. This higher-fidelity method is performed using the CGAL Triangulated Surface Mesh Skeletonization algorithm (Gao et al., 2023). For mesh correspondence and width determinations of all skeletons, the NEURD algorithm employs a custom mesh correspondence workflow based on the following steps: First, each non-branching segment of the skeleton is divided into smaller pieces, and for each piece a cylindrical search radius is used to identify the mesh correspondence. The closest distance between the skeleton and corresponding mesh faces is computed at multiple points along the skeletal segment, and these are averaged to get a mean radius. Finally, all mesh correspondences of sub-branches are combined into the mesh correspondence of the branch. Concatenating the widths along the sub-branches forms a width array along the branch, with the entire branch width determined from the average of the array. This method results in one face of the original branch possibly corresponding to more than one branch’s mesh correspondence, so the algorithm employs a final graph propagation step from unique mesh correspondence faces to allow branches to claim the previously conflicting faces.

The procedure described above yields a collection of disconnected non-branching skeletal segments as well as their associated mesh correspondence. The skeleton of finer-diameter processes is the initial MeshParty Dorkenwald (2022) skeleton which tracks along the mesh surface, while larger-diameter processes have skeletons that track through the center of their volume. These pieces are then all stitched together into a single connectivity graph where each non-branching segment is a node, and the edges between them represent their connectivity. Any conflicts in the mesh correspondence of adjacent nodes at stitching points is again resolved, yielding a smooth and connected mesh representation of the entire stem where each mesh face is associated with a single node (non-branching segment). This entire process is repeated for every stem submesh connected to the soma. Finally, all floating meshes outside the soma are decomposed in the same manner as the stems, and then appended to the existing skeleton if they have any faces within a threshold distance of another node (for example, in the MICrONS dataset the maximum stitch distance was set at 8 *µm*).

The soma(s) and decomposed stems of a neuron are then represented as a NetworkX graph object (Hagberg et al., 2008). In the ideal scenario there is a single soma root node with multiple stem subgraphs, and each stem subgraph is a directed tree structure representing the connectivity between non-branching segments of the skeleton from the most proximal branch connecting to the soma to the most distal leaves of the axonal or dendritic process. In less ideal cases (which are common), cycles may exist in the skeleton due to self-touches of the axonal or dendritic process (close proximities of neurites that are incorrectly meshed together), and multiple somas may be included in a single segmented object. Handling of these cases is described in “Multi-Soma and Multi-Touch Splitting” below. The soma node contains the soma submesh and SDF values generated during the soma extraction, and each branch node in each stem stores the mesh correspondence, skeleton and width array for that node. Using these raw features, many more features of these branches can be extracted (for example spines), and additional annotations can be added (such as synapses).

### Spine Detection

The non-branching segments produced by the mesh decomposition of each node present an ideal scenario for spine detection. Briefly, we started by using an existing mesh segmentation algorithms (Fabri and Rineau, 2023) which calculates a local estimate of the volume for each face of the mesh (SDF), applies a Gaussian Mixture Model (GMM) to the distribution of SDF values to enable a soft clustering of faces to k clusters, and finally minimizes an energy function defined by the alpha-expansion graph-cut algorithm to finish with a hard cluster assignment over the mesh. This last step takes a smoothness parameter controlling the likelihood that adjacent faces with concave edges will be more or less likely to be clustered together. We found that setting the number of clusters to 5 and smoothness value to 0.08 was optimal for both the MICrONS and H01 dataset to produce an over-segmentation of the branch mesh to serve as input for the next spine detection step. Then, we convert the branch segmentation into a graph representation (branch segmentation graph) where the nodes are submeshes of the segmentation and edges exist between submeshes with adjacent faces. The dendritic shaft subgraph is determined by establishing the longest contiguous shaft line path in the graph (from most likely node candidates defined by size, width and diameter thresholds), and then spine candidates are identified as subgraphs (not in the shaft path) based on size, volume, and distance from the mesh surface. The final product of this stage is a collection of individual spine submeshes with calculated spine statistics (volume, length, area). Based on these statistics, in some cases we perform an additional processing step that divides the submesh of larger spines into a spine head and spine neck. At the completion of this workflow, each mesh face in the node receives a spine label of head, neck, shaft or just “spine” (if no head and neck segmentation could be performed). Finally, the width of each branch is recomputed after removing spines that may have previously confounded that measurement. A step by step tutorial of how to optimize the parameters (and a complete explanation of each parameter available for tuning) for the spine detection and head/neck segmentation are included in the tutorials of the Github repository online.

### Synapse Addition

Synapses from the reconstruction pipeline are mapped to the closest mesh face of the closest branch. Any annotations of the associated face (for example) spine head, spine neck or shaft can then be propagated to the synapse. In addition, the closest skeletal point on the associated branch object is computed to define an anchor point for the synapse on the neuron’s skeleton. This anchor point enables computation of metrics such as skeletal walk distance to the closest spine, closest neurite branch point (upstream or dowstream) or skeletal walk distance to the soma.

### Multi-Soma and Multi-Touch Splitting

After the initial decomposition, cycles may exist in the stem graphs due to self-touches in the decomposed mesh (regions where neurites pass very close to each other, resulting in inappropriate connectivity of faces in the mesh representation). Furthermore, stems may include edges with multiple somas if two or more somas are merged together in the same mesh object. This is a challenging problem that requires a general solution since stems can be both multi-touch and multi-soma of any degree. For example, some apical stems of neurons in the MICrONS dataset were initially connected to 9 or more somas due to close mesh proximities with apical tufts of other cells. The aim of this stage in the processing pipeline is to split the stem objects optimally, while attributing the correct portion of the stem mesh and skeleton to the correct neurite.

The workflow for splitting both multi-touches and multi-somas proceeds as follows: For every stem identified as having a multi-touch or multi-soma connection, the process first starts by identifying cyclic or soma-to-soma paths. The best edge to cut on the path is then determined using a series of heuristic rules that are applied in the order listed below:

1. Break any edge on the multi-touch or multi-soma path where the angle between the skeleton vectors of two adjacent branches on the path is larger than some threshold (reflecting the fact that neurite processes generally do not abruptly double-back on themselves).
2. Break any edge on the multi-touch or multi-soma path where more than two downstream branches exist at a branch point and the best match for skeletal branch angle and width is not on the multi-touch or multi-soma path.
3. Break any edge on the multi-touch or multi-soma path where there is a difference in width between two nodes along the path greater than a threshold amount.

After an edge is removed based on any of these rules, the process restarts and the graph representation is checked again to see if cyclic or multi-soma paths still exist. The process is repeated until no such paths exist. If no candidate edge is identified by using these rules then, depending on the user settings, the the stem may be completely discarded from the neuron object or cut at the very first or last branch.

This splitting algorithm is not guaranteed to optimally split all multi-soma or multi-touch paths, but residual errors from a sub-optimal split may be cleaned by further proofreading steps. As with any automated proofreading, the rule and relevant parameters that determined the edit are stored for subsequent evaluation and use.

### Excitatory/Inhibitory Classification

Once each neuron object has a single soma, the NEURD workflow moves on to an initial round of coarse cell classification, determining whether each neuron is excitatory or inhibitory. Performing the classification at this point in the workflow enables the use of subsequent proofreading or annotation algorithms that are excitatory- or inhibitory-specific. For example, axon identification (see below) is much easier if the coarse E/I type of the cell is known beforehand. The cell class is determined via logistic regression on two features: postsynaptic shaft density (number of synapses onto dendritic shafts per *µm* of skeletal distance on the postsynaptic dendrite) and spine density (number of spines per *µm* of skeletal distance on the postsynaptic neuron). These two features enabled linearly-separable elliptical clusters for excitatory and inhibitory cells. To enable this classification prior to proofreading, we applied two restrictions to the unproofread graph that reduce potential confounds due to merge errors and ambiguity between axon and dendrite. First, we restrict to larger dendrites using a simple width threshold to exclude potential orphan axon merges, and second, we restrict to the proximal dendrite within a limited skeletal walk distance from the soma center. The latter reduces confounds due to dendritic merge errors which are more common at the distal branches (Fig. 3d). When compared to human E/I labels, the classifier results are shown in Fig. 17a,b. These results are also robust against an approximate 10:1 and 1.8:1 excitatory to inhibitory class imbalance in the labeled MICrONS and H01 datasets respectively. Additionally, because a logistic regression is the classifier, there is a confidence score associated with the classification (which could be thresholded downstream to ensure higher fidelity labels).

### Non-Neuronal Filtering

While all the neurons in the H01 dataset were hand-checked as neurons and manually annotated for cell types, the MICrONS dataset initially was not. Consequently, segments with nuclei in the MICrONS dataset could include blood vessels, glia cells, or agglomerations of orphan axons without a neuron mesh due to an incorrect nucleus merge, and we did observe this frequently in the version of MICrONS processed in this study (version 374). However this issue is now largely resolved in the most current data release with tables that indicate which segments are predicted as non-neuronal, using a method independent from NEURD, and with more accurate nuclei merging. To filter away non-neuronal segments without sacrificing a significant amount of valid neurons, we found a suitable filter using cell type classification (predicted by our logistic regression model), number of soma synapses and the mean dendritic branch length (this may very well be a MICrONS-specific filter). Specifically, the filter excluded the following: all segments with less than 3 soma synapses, excitatory cells with less than 17 soma synapses and a mean dendritic branch length less than 35 *µm*, and inhibitory cells with less than 17 soma synapses and a mean dendritic branch length less than 28 *µm*. This filter then removed approximately 14,000 segments from all of our downstream analysis.

### Axon Identification

The goal of this stage in the pipeline is to identify at most one connected component subgraph that represents the axon of the cell. In the absence of merge and split errors, identifying the axon would be a simple process of identifying the subgraph with presynaptic connections, but un-proofread datasets pose a number of challenges to this approach.

1. Due to partial reconstruction of cells, the true axon may not exist or only the axon initial segment (AIS) may exist. In this case there would be no true presynaptic connections from the cell.
2. Postsynaptic synapses on dendritic segments may be incorrectly labeled as presynaptic connections if the synapse classifier is incorrect.
3. Orphan axons may be incorrectly merged onto the cell’s dendrite or soma. These frequent merge errors add incorrect presynaptic connections onto dendrites that make identifying the true axon subgraph more difficult. We observed in these volumes that if the algorithm simply chose the connected component subgraph with the highest presynaptic density, this would almost always be an orphan-axon-onto-dendrite merge error.

Our approach to axon identification is thus motivated by the following neuroscience “rules” which we implement as heuristic selection criteria. Note that in this and following sections “up” or “higher” refers to the pial direction, while “down” or “lower” refers to the white matter.

1. Axons can either project directly from the soma or from a proximal dendritic branch.
2. The axon is the only compartment (possibly excluding the soma) that forms presynaptic connections.
3. The width of axon segments are typically thinner than most dendritic branches
4. Axon segments do not have spines (although boutons may have similar features)
5. Axons receive postsynaptic inputs at the AIS, but these typically have low postsynaptic density compared to dendrites. (Note that we found the latter was not necessarily true in the H01 dataset and adjusted accordingly.)
6. For excitatory cells, the axon typically projects directly from the soma or from dendritic stems that originate from the deeper half of the soma.
7. For excitatory cells, the AIS starts at most 14 *µ*m skeletal distance from the soma and the mean skeleton vector of the AIS typical projects downwards. The AIS does not split into multiple branches close to the soma.
8. For inhibitory cells the AIS can start much farther away (up to 80 *µ*m skeletal distance from soma) and can come off the soma or dendritic branch. The inhibitory AIS has very low postsynaptic and presynaptic density.

NEURD identifies candidate axonal submeshes based on a combination of these heuristics applied in a cell-type-specific manner. For example, if the neuron being analyzed is excitatory, the search space of potential axonal stems is restricted to only those with a projection angle from the soma greater than 70 degrees relative to the top of the volume. Candidate AIS branches must fall within a maximum and minimum width range, they must have a synapse density below a threshold value, and they must be within a threshold skeletal distance from the soma that dictated by the cell type. If multiple potential candidates exist, the best potential axonal subgraph is selected based on longest skeletal length and closest proximity to the soma. Subgraphs that meet the heuristic criteria for axons but are not chosen as the actual axon of the cell in question retain a label of “axon-like” which facilitates subsequent proofreading.

An additional round of skeletonization is performed once the axon is detected. This re-skeletonization better captures fine details of the axonal trajectory and enables auto-proofreading methods to catch more subtle axon-to-axon merge errors.

### Automatic Proofreading

The goal of the automatic proofreading stage is to identify merge errors and remove all downstream branches. NEURD implements a series of heuristic proofreading rules to identify merge errors based on graph filters - configurations of nodes and attributes that typically indicate merge errors. The graph filters are either directed one-hop or zero-hop configurations where one-hop configurations consider a node and its immediate adjacent nodes and zero-hop configurations consider only the node’s features itself. These graph filters have parameters that can be tuned for axons or dendrites, excitatory or inhibitory cells, or different data sources (H01 vs MICrONS). For example, a graph filter for resolving graph configurations of dendritic branches with 3 or more downstream nodes is useful for the H01 dataset which has a higher probability of dendritic merge errors than MICrONS. For each filter, the algorithm finds all branches that match the current error motif, and then the mesh, graph nodes and synapses associated with all those error branches are removed before proceeding to the next filter. Therefore, in the current successive order there is no overlap in error locations for different filters in the same run. Metadata for each correction is stored for subsequent review or for training non-heuristic models.

The following graph filters exist for proofreading axon submeshes. Those only used for excitatory cell types are indicated (visualizations shown in 8).

1. <u>High Degree Branching</u>: The filter identifies any node (below a potential width threshold to exclude myelinated sections) with more than two downstream nodes. The filter assumes this configuration results from a single or multiple crossing axon(s) merged at an intersection point, adding 2 or more additional downstream nodes. The filter aims to identify the one true downstream node. Possible upstream to downstream node pairings are filtered away if the width, synapse density or skeletal angle differ by threshold amounts. If multiple downstream nodes are viable, the algorithm attempts to find a downstream candidate with the best match of skeletal angle or width. If no clear winner exists, the algorithm can either mark all downstream nodes as errors if the user wishes to be conservative, or can pick the best skeletal-match candidate. There are more rare scenarios where a myelinated axon has 2 collateral projections protruding very close to one another and these would be incorrectly filtered away, but a large majority of these occurances are simply merge errors.
2. <u>Low Degree Branching Filter</u>: The filter processes any subgraph with one upstream node and exactly two downstream nodes and is only attempted on non-myelinated axon sections (as determined by a width threshold). The method attempts to find one of the following subgraph features within this directed 3 node subgraph, and if a match occurs either the algorithm marks all of the downstream nodes as errors or attempts to determine the correct downstream pairing.

a. **Axon Webbing**: An error is detected by an overly-thin mesh at the branching point of an upstream to downstream node. The filter attempts to differentiate between natural branching with cell membrane that forms a “webbing like” appearance as opposed to merge errors where no such thickening occurs.
b. **T-Intersection**: An error is detected by the presence of downstream branches that are thicker than an upstream branch and the downstream branches are aligned and resemble a continuous non-branching axon segment.
c. **Double-Back** (excitatory only): An error is detected when a downstream node “doubles-back” towards the upstream node at an unnatural skeletal angle.
d. **Parallel Children (or Fork Divergence)**: An error is detected when the two downstream skeletons are nearly parallel without a natural gap between them.
e. **Synapse At Branching**: An error is detected if a synapse occurs very close to the branch point between upstream and downstream nodes; this usually indicates a merge of an orphan axon to a bouton segment.
3. <u>Width Jump</u>: Processes a subgraph with any number of downstream nodes (only applies to non-myelinated sections). Any downstream node with an absolute width difference between segments above a certain threshold is marked as an error.

The following graph filters exist for proofreading dendrite submeshes.

1. <u>Axon on Dendrite</u>: Nodes that were previously labeled “axon-like” during the process of axon identification (see above) that do not end up in the axon submesh are marked as errors.
2. <u>High Degree Branching</u> (H01 only, excitatory only, apical trunks excluded): An error is detected using the same algorithm as described above for the axonal High Degree Branching Filter except it is applied to dendritic nodes below a thresholded width.
3. <u>Width Jump</u>: An error is detected using the same graph filter as described for axons but with larger width difference thresholds
4. <u>Double Back</u>: An error is detected using the same algorithm and parameters as for axons above.

### APL Validation of Multi-Soma Splitting and High Confidence Orphan Merge Edits

Our collaborators at APL (Johns Hopkins University Applied Physics Laboratory) helped extensively validate multiple aspects of the NEURD automated proofreading workflow. In particular, they provided information about the following:

1. Validation of specific edits in the context of multi-soma splitting.
2. Data about the time that proofreaders took to evaluate these split suggestions compared to other methods.
3. Validation of specific edits focused on axon-onto-dendrite or axon-onto-axon merges

This information made it possible to determine whether our suggestions for multi-soma split locations speed up the process (they do, more than three-fold), and if we could identify a set of heuristics and parameters for axon-on-dendrite and axon-on-axon merge corrections that could be executed with high confidence on the entire volume without human intervention (we identified two classes of edits with performance >95%, and more than 150,000 of them have been applied to the MICrONS volume to date; see Fig. 9).

A key method for both validating and applying automatic edits was the functionality in Neuroglancer (Perlman, 2019) which allows the placement of point annotations to define a split in the PyChunkedGraph segmentation (Dorkenwald et al., 2022a,c). To facilitate the proofreading process, APL created a web-based interface called NeuVue (Xenes et al., 2022) that allows for the efficient queuing, review and execution of split suggestions in Neuroglancer. We built the logic required to translate mesh errors identified by NEURD into split point annotations that can be executed by the NeuVue pipeline. This capability allowed proofreaders at APL to not only evaluate error locations identified by NEURD, but also a proposed set of points that could be subsequently executed in the PyChunkedGraph to correct the error. For a more detailed description of the NeuVue review pipeline see (Xenes et al., 2022).

For multi-soma split edits, we generated point annotations for suggested splits that would contribute to separating neurons with between 2 and 6 possible somas in a single segment. APL had both experts and trained student proofreaders review these edits. The classifications for each of the edits was one of the following:

1. “yes”: same split point annotations the proofreader would have chosen.
2. “yesConditional”: split point is correct, but point annotations required very minor adjustment.
3. “errorNearby”: split point is not correct but is very close by, and split point annotations require adjustment
4. “no”: the correct split location was not at or near the suggested location

The expert proofreaders reviewed 5134 unique suggestions with no overlap between proofreaders, while the student proof-readers reviewed 2355 suggestions with some redundancy so the same suggested edit was seen by multiple student proofreaders and a majority vote determined the classification of the edit. The results of reviewing the first approximately 4000 of those edits are shown in Fig.9a and the accuracy was determined to be 76.12% when the “yes”,”yesConditional” and “errorNearby” categories were considered true positive classes. Additionally, because each split suggestion had an associated heuristic rule and set of parameters that was used to generate the suggestion, we were able to show that some rules were much higher fidelity than others, and that the parameters could be tuned to achieve a higher classification accuracy (Fig.9b,c).

To compare against the performance achieved with NEURD suggestions, multi-soma splits were also performed by expert proofreaders using a tool that highlighted the path along the neural processes connecting two somas. This comparison enabled us to measure if the NEURD suggestions could potentially speed up the soma-splitting process. Because a single segment with a multi-soma merge could contain more than 2 somas and because different merges may require a different amount of work and number of cuts to be applied, we measured the overall time spent reviewing all edits and estimated the additional time that would have been required to make the slight adjustments required in the case of “yesConditional” (+30s) or “errorNearby” (+60s), prior to executing the edit. Note that “no” classifications added time to the review process without any possibility of contributing to an actual edit. Based on these metrics, we divided total time by total number of edits to determine the mean time per edit. We then compared this metric to the mean time per edit when proofreaders used a standard pathfinding tool that displayed the skeletal path connecting multiple somas, and they had to search along this path to identify errors manually. We observed a more than three-fold speed up when using NEURD suggestions (Fig.9d).

Finally, outside the context of multi-soma splitting, APL proofreaders evaluated two kinds of merge error corrections that strip orphan axons from both excitatory and inhibitory neurons: axon-on-dendrite, and high degree axon-on-axon. The feedback on each error from proofreaders using the NeuVue pipeline allowed us to determine a subset of parameters that was correlated with high accuracy. Thresholds for the axon-on-dendrite included minimum parent width, distance from the soma, and skeletal length of the error segment. Thresholds for axon-on-axon included a minimum skeletal length, and a branching pattern that resembled a two line segment crossing, where the segments are closer to perpendicular in order to make the correct connectivity more obvious. For the review of orphan merge errors, an additional label was included in the true positive class: “yesPartial”, which indicated that part but not all of the merge was removed by the split point annotations. The feedback from this effort provided our collaboration with enough evidence to then apply nearly 150,000 of these high-confidence automatic edits back into the current dynamic segmentation of the MICrONS dataset (Fig.9e).

### Automatic Compartment Labeling

After automatic proofreading removes as many merge errors as possible, compartment labeling is performed for excitatory cells, classifying graph submeshes as apical trunk, apical tuft, basal, and oblique. NEURD first attempts to identify the apical trunk based on the geometry relative to the soma and total skeletal length. Branches downstream of the end of the apical trunk are classified as the apical tuft, and branches off of the trunk with a skeletal angle close to 90 degrees are labeled as oblique. If the criteria for a defined trunk is not met, then NEURD applies a generic “apical” label. Other dendrites are classified as basal. Additionally, for the MICrONS dataset if the soma center is close enough to the pia as defined by a depth threshold, there can be multiple generic “apical” stems protruding from the top of the soma if they each meet the required width and geometry thresholds.

### Connectome-level features computed by NEURD

At the level of the connectome graph, nodes represent individual singlenucleus neurons and edges represent synaptic connections. In addition to the rich sub-cellular features that NEURD computes for the decomposition graph of each cell, NEURD provides a variety of features at the connectome graph level:

1. Node Attributes: a wide range of global properties measured for the individual cells (compartment skeletal lengths, synapses, bounding box, spine densities, synapse densities, average width, cell type, etc).
2. Edges: connections between neurons with a valid presynaptic connection and postsynaptic connection where neither were filtered away in the auto-proofreading stage
3. Edge Attributes: properties for each of the presynaptic and postsynaptic neurons (compartment, skeletal/euclidean distance to neuron’s soma, size, spine label) and properties of the entire synaptic connection between neurons (euclidean/skeletal distance from soma of presynaptic neuron to soma of postsynaptic neuron, etc).

### GNN Cell Typing

Using PyTorch geometric software (Fey and Lenssen, 2019) we implemented a Graph Neural Network architecture to build a supervised cell type classifier (including subclasses of excitatory and inhibitory cells) from dendritic graph structure in the NEURD decompositions. We trained this classifier using manual cell types from the Allen Institute for Brain Science (Schneider-Mizell et al., 2023). To create an input graph for the classifier we first removed the soma node and filtered away the axonal subgraph and any dendritic stems with less than 25 *µ*m of total skeletal length. Each node was annotated with the following feature set:

1. Skeleton features, where theta and phi refer to polar coordinates of the skeleton vector in 3-D (skeleton_length, skeleton_vector_upstream_theta, skeleton_vector_upstream_phi, skeleton_vector_downstream_theta, skeleton_vector_downstream_phi)
2. Width features (width, width_upstream, width_downstream synapse)
3. Spine features (n_spines, spine_volume_sum, n_synapses_post, n_synapses_pre, n_synapses_head_postsyn, n_synapses_neck_postsyn, n_synapses_shaft_postsyn, n_synapses_no_head_postsyn, synapse_volume_shaft_postsyn_sum, synapse_volume_head_postsyn_sum, synapse_volume_no_head_postsyn_sum, synapse_volume_neck_postsyn_sum, synapse_volume_postsyn_sum)

For the whole neuron classifier, the soma volume and number of soma synapses for the neuron are added to each node’s feature vector and also the starting stem angle (2-D angle between the vector from the soma center to the stem’s root skeleton point and the vector in the direction of the pia) is added to each node in every stem. For the stem-based classifier, these three soma features are not included, and classification is performed on each stem individually.

The GNN architecture used as a 2 layer Graph Convolutional Network (128 hidden units for each layer, ReLU activation function) followed with one linear layer. The aggregation and update steps were implemented using self loops and symmetric normalization as shown here:

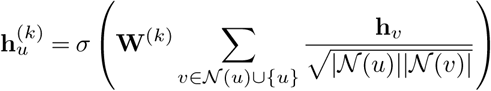

where **h***_u_*^(^*^k^*^)^ is the embedding for node u at layer k, *N* (*u*) are the neighbors for node u, **W**^(^*^k^*^)^ is the learned weight matrix at layer k and *σ* is the chosen non-linearity. For graph pooling (to get one learned vector for each graph), the weighted average of all nodes after the final hidden layer was taken (weighted by the skeletal length of the node). A 60%, 20%, 20% split for training, validation and test sets was used for labeled datasets of n = 873 whole neurons and n = 4,114 stems (Fig.19)

### Proximities

Identifying axon-dendrite proximities makes it possible to determine how often a pair of neurons capitalizes on an opportunity to form a synaptic connection. Proximities are regions where the axon of one neuron passes within a few microns of the dendrite of another neuron. They can be annotated with the same features (dendritic compartment, neural subtype) as synapses, regardless of whether a connection was formed (Fig. 5a). Proximities are identified for all neuron pairs in the volume. To reduce the number of pairwise computations, NEURD first checks whether the bounding box of the presynaptic axon skeleton and postsynaptic dendrite skeleton have any overlap. In order to reduce computation time in the the MICrONS dataset, presynaptic neurons are further restricted to those with at least five axonal synapses, and in the MICrONS volume postsynaptic neurons are restricted to neurons with at least 1 mm of dendritic length (this latter restriction excludes approximately 1% of all MICrONS neurons).

The proximity calculation is performed by converting the axonal skeleton of the presynaptic neuron and the postsynaptic skeleton to an array of coordinates without edges (at one *µm* skeletal walk resolution). A local width and compartment label is associated with every point, and the soma of the postsynaptic neuron is converted to a uniform sampling of the surface mesh face centers for its skeletal representation. The cleaned synaptic connections between the pre and post neuron are retrieved (if there were any) and then the main proximity loop begins.

1. The closest distance between a presynaptic coordinate and postsynaptic coordinate is computed (where the distances can be adjusted for postsynaptic width by subtracting the local width from the euclidean distance). This minimum distance is the current proximity distance. If the current proximity distance exceeds the thresholded maximum proximity distance (set at 5 *µ*m for both the MICrONS and H01 dataset), then the loop is exited and no more proximities are computed, but otherwise the workflow proceeds.
2. The following metrics are computed or collected for each proximity: The presynaptic and postsynaptic coordinate of the minimum distance pair, the distance between these coordinates (the proximity distance), the compartment labels and width at the postsynaptic coordinate, the presynaptic and postsynaptic skeletal walk distance, the number of spines and synaptic connections within a three *µm* radius of the presynaptic and postsynaptic coordinates.
3. After these features are collected, the skeleton points within a set radius (10 *µm*) of the presynaptic proximity coordinate are filtered away from the array of axon presynaptic coordinates.
4. All proximity information is saved and the loop continues until the current proximity distance exceeds the threshold.

### Functional Connectomics

We considered pairs of synaptically-connected functionally-matched cells available in the MICrONS dataset, restricting to pairs where both neurons met a minimal set of functional quality criteria (test score greater than 0.2 and an oracle score (correlation of a neurons response to the leave-one-out mean response across a repeated image stimuli) greater than 0.3, see Ding et al. 2023; Wang and Tolias 2023). Synaptic connections were discarded if they were not onto post-synaptic spines (to help guard against possible inhibitory merge errors resulting in increased connectivity between neurons). We then divided the pairs into groups based on whether they had 1, 2, 3 or 4+ synapses between them. The final number of functionally matched pyramidal pairs available from automatic proofreading alone were as follows: 1 synapse (5350), 2 synapses (280), 3 synapses (34) and 4+ synapses (11). We then investigated how the mean functional response correlation varies as a function of the four different multi-synaptic groups. The response correlation was calculated as detailed in (Ding et al., 2023; Wang and Tolias, 2023) through the *in silico* response correlation of their model.

### Analysis Subsets and Distributions

All morphological, connectomic and functional connectomic analyses excluded the manually proofread neurons used for validating NEURD proofreading, a small fraction of cells that errored out during the NEURD preprocessing pipeline due to a variety of factors (corrupted mesh, manifold and watertight properties unable to be automatically fixed, etc.), and non-neuronal cells that were filtered away as described in the “Non-Neuronal Filtering” section. Additionally, we excluded some neurons with very incomplete reconstructions from some analyses. For example, only a subset of automatically proofread neurons with an axon length longer than 50 *µm* were used in the investigation of questions concerning synapses onto the axon initial segment, in order to not skew results due to a reconstruction bias. Analyses of connectivity, etc between different cell subtypes only used neurons with high confidence labels from the GNN classifier (softmax output >0.7). Complete Ns for all analyses and different stages of the pipeline are included in the Appendix Table 2.

### Supplementary Discussion

Highly-annotated NEURD graphs provide a compact representation of many features that are useful for all kinds of morphological analysis. For example, a simple query reveals that the percentage of pyramidal cells with axons protruding from dendrites (rather than the soma) is higher in the mouse volume (17.8%) than the human volume (8%), which closely replicates the findings of an entire previous study focused on this question (Wahle et al., 2022). Several of the morphological properties of cell types shown in Fig. 4g-h and Fig. 20 replicate observations from previous studies (Elabbady et al., 2022; Kawaguchi et al., 2006; Villa and Nedivi, 2016). Additionally, using the spine metrics extracted by NEURD, we were able to replicate many of the findings of (Harris and Stevens, 1989; Arellano et al., 2007) concerning synapse size and spine head volume correlation, and we also show that these scaling rules and others depend on cell type (Fig.4h, Fig. 25). Looking at how synapses onto the AIS and soma vary across species, we find the expected lower rate of soma synapses on human neurons than mouse (Wildenberg et al., 2021), and replicate the expected distribution of AIS synapses across depth in the mouse dataset (Schneider-Mizell et al., 2021). We also find that the human dataset does not show a similar change over depth in AIS synapses, and that the human dataset AIS is more densely innervated than in the mouse volume (Fig.21g-h, Fig. 21i). Finally, we demonstrate the use of a query combining geometric information and branch-level characteristics to replicate the previously-reported bias in the orientation of the thickest basal segment in the H01 dataset (Shapson-Coe et al., 2021). We extend this finding with an observation that this bias is actually consistent across all depths but is just less salient in upper layers because the relative size of the thickest and second-thickest basal dendrites changes smoothly across depth (Fig.22b,c).

Using the cell type node labels and the skeletal walk length edge features, we confirmed previous work (Hwang et al., 2021; Megıas et al., 2001) describing different distal and proximal preferences for different excitatory and inhibitory connection types (Fig. 5d-e, Fig. 28f-g). Furthermore, using the proximity controls computed on the cleaned skeletons, we also were able to observe a consistent trend across datasets showing the propensity for forming connections from excitatory to inhibitory neurons peaks around 200 *µ*m away from the soma, potentially consistent with a pattern of surround suppression (Fig.28d,e). Additionally, with the cell type node labels and spine compartment and synapse size edge labels, we confirmed a variety of expected findings about synaptic and spine head size: excitatory to excitatory connections have the largest synapses, synapse size correlates with spine heads for excitatory sources but not inhibitory sources, and spine heads with inhibitory synapses generally are multi-synaptic spines where the inhibitory synapse is typically much smaller than the largest synapse on the spine head (Fig. 26).

## AUTHOR CONTRIBUTIONS

We use the CRediT system for author roles. Conceptualization: BC, JR. Methodology: BC, SP, JR. Software: BC. Validation: BC, DX, LMK, PKR, VAR, CAB, BW, FC. Formal analysis: BC. Investigation: BC. Resources: BC, SP, ZD, PGF, EW, CP, AK, SP, JAB, ALB, DB, JB, DJB, MAC, EC, SD, LE, AH, ZJ, CJ, DK, NK, SK, KL, KL, RL, TM, GM, EM, SSM, SM, BN, SP, CMSM, WS, MT, RT, NLT, WW, JW, SCY, WY, EF, EYW, FHS, HSS, FC, NMdC, RCR, XP, AST, JR. Data Curation: BC. Writing - Original Draft: BC, JR. Writing - Review Editing: BC, SD, CMSM, FC, NMdC, AST, JR. Visualization: BC, JR. Supervision: JR. Project administration: JR. Funding acquisition: HSS, NMdC, RCR, AST, JR.

## ACKNOWLEDGEMENTS

The authors thank David Markowitz, the IARPA MICrONS Program Manager, who coordinated this work during all three phases of the MICrONS program. We thank IARPA program managers Jacob Vogelstein and David Markowitz for co-developing the MICrONS program. We thank Jennifer Wang, IARPA SETA for her assistance. The work was supported by the Intelligence Advanced Research Projects Activity (IARPA) via Department of Interior/ Interior Business Center (DoI/IBC) contract numbers D16PC00003, D16PC00004, D16PC0005, and 2017-17032700004-005. The views and conclusions contained herein are those of the authors and should not be interpreted as necessarily representing the official policies or endorsements, either expressed or implied, of the ODNI, IARPA, or the U.S. Government. The U.S. Government is authorized to reproduce and distribute reprints for Governmental purposes notwithstanding any copyright annotation thereon. ABK acknowledges support from a training fellowship from the Gulf Coast Consortia, on the NLM Training Program in Biomedical Informatics & Data Science (T15LM007093). XP acknowledges support from NSF CAREER grant IOS-1552868. XP, ABK, and AST acknowledge support from NSF NeuroNex grant 1707400. AST also acknowledges support from the National Institute of Mental Health and National Institute of Neurological Disorders And Stroke under Award Number U19MH114830 and National Eye Institute award numbers R01 EY026927 and Core Grant for Vision Research T32-EY-002520-37. AST, XP, BC and JR are supported by RF1 MH130416 to AST, XP, JR. Disclaimer: The views and conclusions contained herein are those of the authors and should not be interpreted as necessarily representing the official policies or endorsements, either expressed or implied, of IARPA, DoI/IBC, or the U.S. Government.

## Software

Experiments and analysis are carried out with custom built data pipelines. The data pipeline is developed in Python with the following tools: DataJoint used for storing and managing data. Meshparty, CGAL and MeshLab were used for mesh processing (CGAL and MeshLab required python wrappers). DotMotif and NetSci was used to help query graph motifs. Numpy, Pandas, SciPy, Scikit-learn, and PyTorch were used for model training and statistical analysis. Matplotlib, Seaborn, Ipyvolume, and Neuroglancer were used for graphical visualization. Jupyter, Docker, and Kubernetes were used for code development and deployment.

## Data availability

All MICrONS data have already been released on BossDB (https://bossdb.org/project/microns-minnie, please also see https://www.microns-explorer.org/cortical-mm3) for details.

## Code availability

Code, documentation, and example tutorials are available at https://github.com/reimerlab/NEURD.git

## Supplementary Figures

**Fig. 6.**
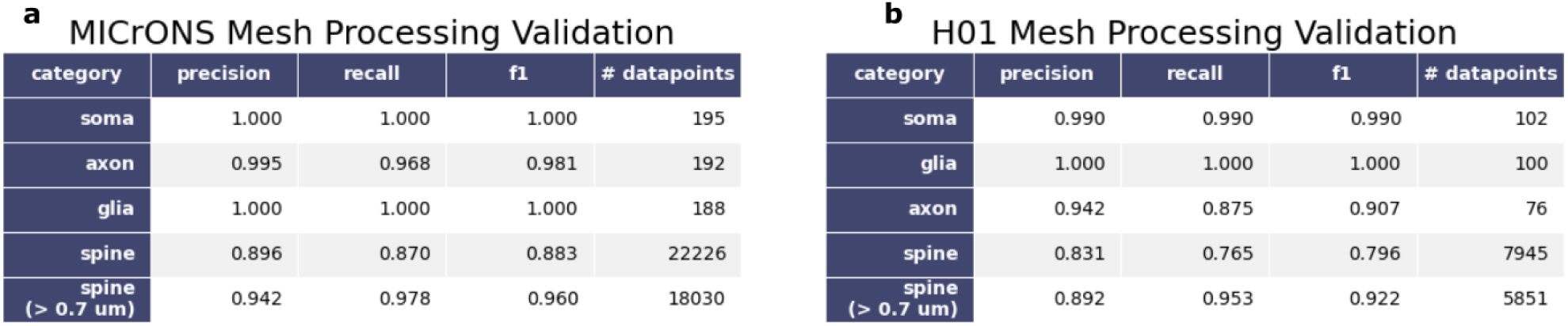
Mesh Processing Pipeline Validation. **a** (MICrONS) Validation scores of automatic submesh (compartment) identification in comparison to human labels. This dataset was produced by randomly presenting mesh segments to human proofreaders who evaluated automatic annotations as true positive, true negative, false positive, or false negative (TP,TN,FP,or FN) for each structure. Only glia merges larger than the volume of a 5 *µ*m radius sphere were considered glia merges in this processing step. The “spine (> 0.7 *µ*m)” row reports the agreement between the automatic spine detection and human spine labeling for spines with a skeletal length greater than 0.7 *µ*m. Below this threshold there was disagreement even among human proofreaders about whether small protrusions should be classified as a spine or not. Precision = true positives / (true positives + false positives), Recall = true positives / (true positives + false negatives), F1 = (precision * recall)/(precision + recall). The F1 statistic serves as a weighting between the recall and precision to ensure a degenerative solution is not achieved (Ex: marking all samples as positives in order to maximize recall at the detriment of precision). **b** Identical validation scores for H01 dataset.

**Fig. 7.**
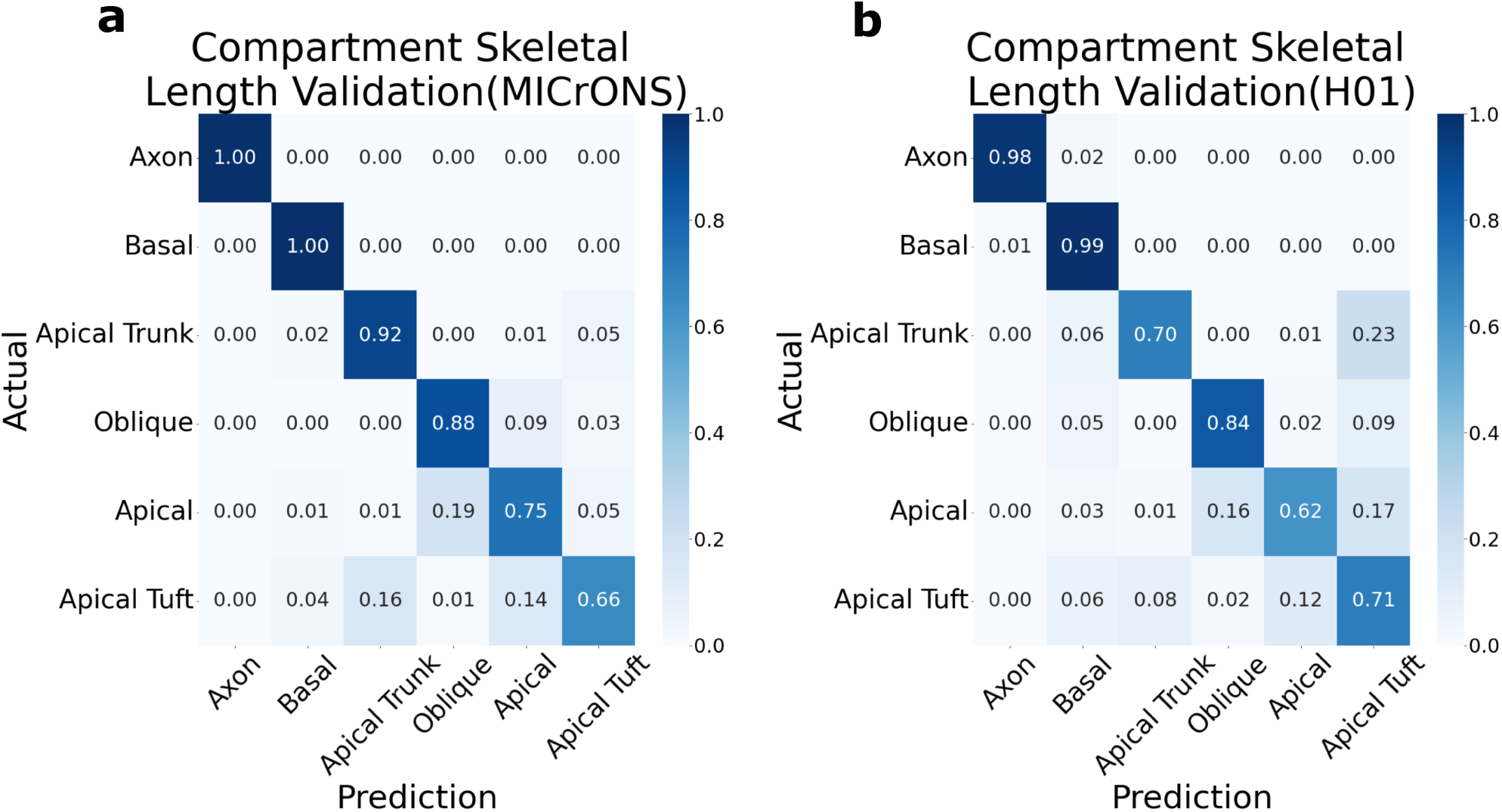
Automatic Submesh (Compartment) Labeling Validation. **a** (MICrONS) Confusion matrix comparing automatic submesh labeling to human labels (random sampling of 158 processed cells). Here the TP,FP,TN,FN metrics are computed using skeletal length agreement for each compartment. Therefore, cells with longer stems or cells with more stems of a certain compartment type more heavily influenced the scores due to the skeletal length weighting. **b** (H01) Confusion Matrix comparing automatic submesh labeling to human labeled submeshes (random sampling of 89 processed cells). Across both datasets, compartment labeling was nearly perfect for separating the axon, basal, and apical supercategory (the union of oblique, apical, apical tuft and apical trunk) compartments, but was less consistent for separating the sub-compartments of the apical supercategory.

**Fig. 8.**
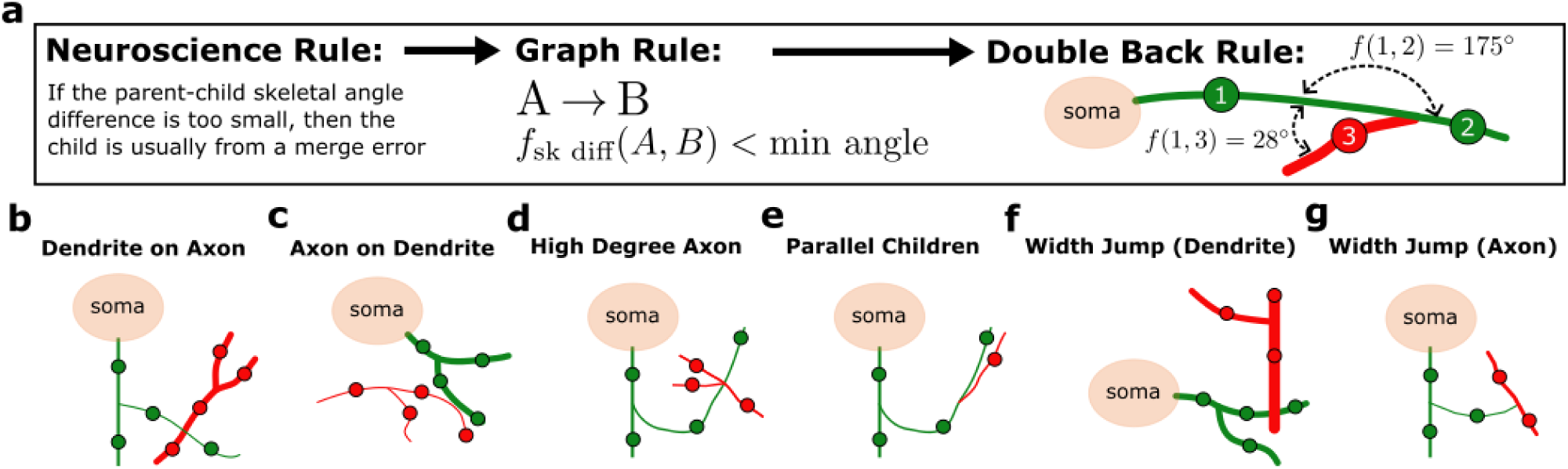
Automatic Proofreading Rule Visualizations. **a** Example implementation of domain knowledge as a subgraph rule to automatically identify and remove merge errors. Most of the same rules can be applied across excitatory and inhibitory cells in the MICrONS and H01 volume as-is, or with small changes in parameters. **b-g** Visual representation of other subgraph rules implemented for the automatic proofreading stage as described in the “Methods-Automatic Proofreading” section.

**Fig. 9.**
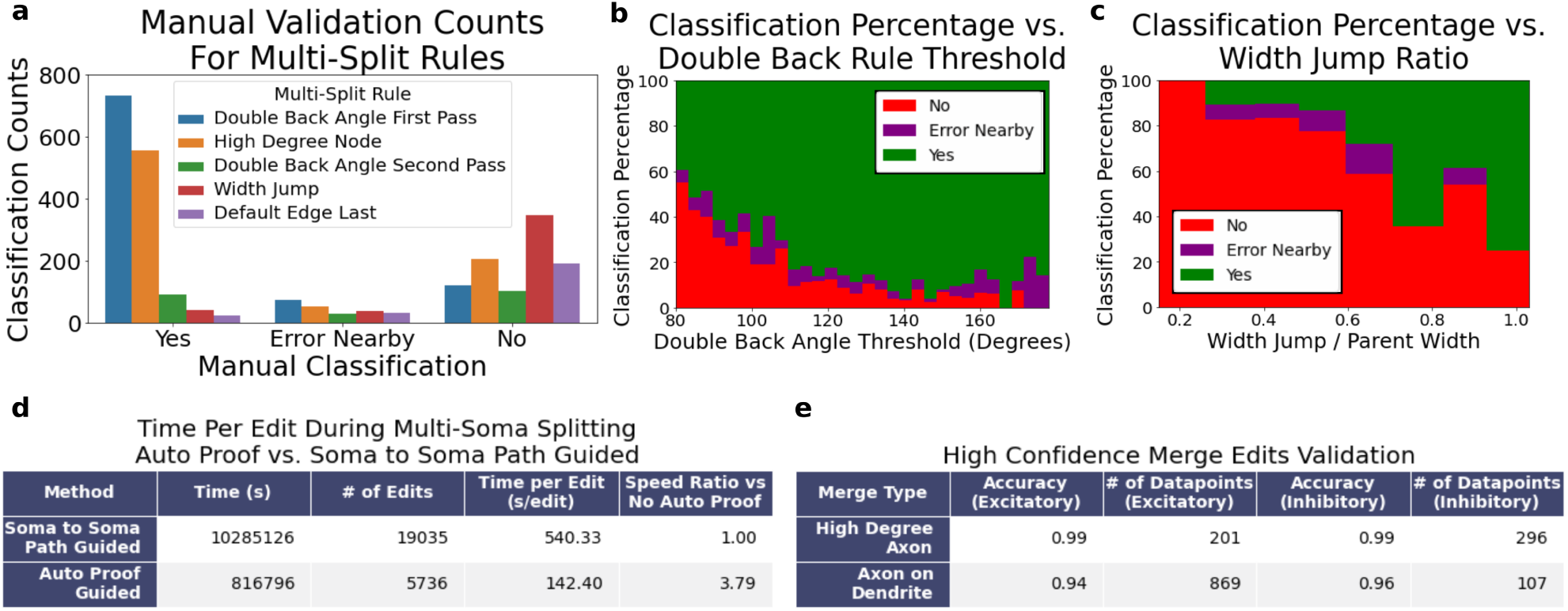
Proofreading Validation. All validation was performed by the proofreading team at Johns Hopkins University Applied Physics Laboratory (APL). In an initial round of validation, suggested error locations were evaluated in the context of splitting multi-soma cells in the MICrONS volume. As a result we were able to measure both the accuracy of these proofreading rules and the speed benefits of a semi-supervised approach compared to fully-manual proofreading. Additionally, the accuracy of two automatic proofreading rules with high-confidence parameters (axon on dendrite, high degree axon on axon merges) were evaluated. **a** Validation of split locations predicted by automatic multi-split algorithm. “Yes” (indicates that the proposed split can be executed immediately), “Error Nearby” indicates that the split location is correct within 20 *µ*m, but that the human proofreader slightly modified the suggested split points (this threshold was determined so that every suggestion in this category would prove useful to and increase the speed of human proofreaders after their attention is drawn to the relevant location), and “No” indicates that the true split location was far from the predicted location or no merge error was detected by the human proofreader). The heuristic splitting rules are applied in the order indicated by the legend. The automated proofreading accuracy varied substantially over the different heuristic rules with an overall accuracy of 76.12 % when “Yes” and “Error Nearby” are considered true positives. The best-performing rules can be selected for different datasets. **b** Even for a single rule, thresholds can be tuned to optimize performance. Manual classification of split locations predicted by the “Double Back” rule as a function of the angle measured at each predicted location illustrates that a higher accuracy could be achieved by setting a higher threshold for this algorithm. **c** Manual classification of split locations predicted by the width jump rule as a function of the width jump at each predicted location, illustrating another example where interpretable thresholds can be adjusted for higher precision. **d** Elapsed time statistics as humans performed manual tasks of splitting multi-soma neurons either using a tool that showed the path along the mesh between two somas or using the suggested split locations from our automatic multi-split algorithm. The speed at which humans could apply cuts in the correct locations more than tripled when using suggestions provided by the NEURD multi-split algorithm. Note: the validation is measured as the average amount of time for a single edit in the multi-soma splitting process; a single multi-soma split might require 20 or more edits to completely resolve the merge. **e** Accuracy of two automatic proofreading rules with high-confidence parameters (high degree axon on axon, axon on dendrite).

**Fig. 10.**
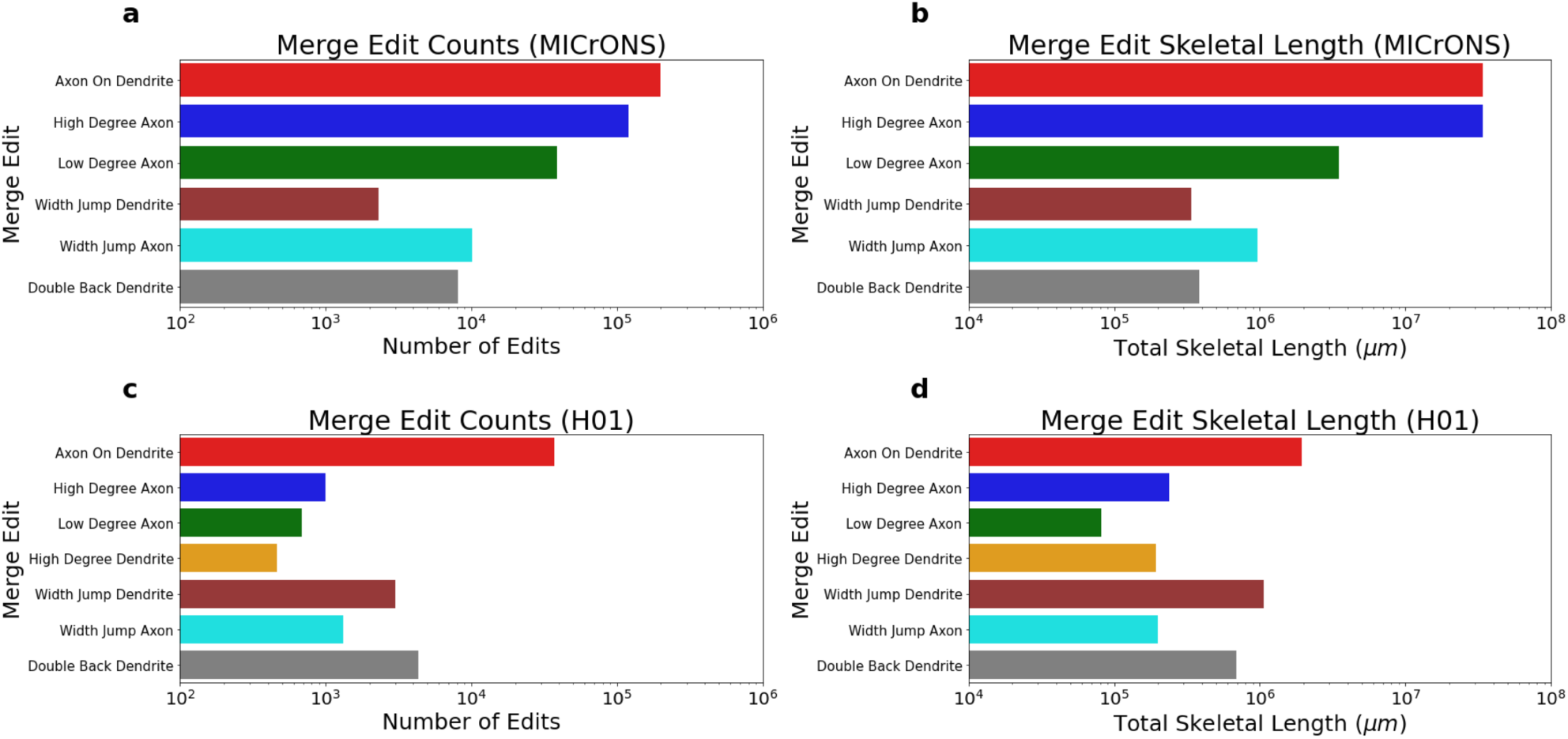
Counts and total skeletal length of merge errors corrected during automated proofreading. The different heuristic rules are presented from top to bottom in the order that they are implemented in the automated proofreading workflow. All error segments identified are then returned to the set of non-nucleus-associated fragments and are not included in the morphological or connectivity analyses. Note that errors identified by rules later in the workflow are excluded from these statistics if they are found on already-errored segments identified earlier in the workflow. **a** (MICrONS) Total number of separate locations where a specific heuristic rule corrected a merge error. **b** (MICrONS) Total skeletal length eliminated by each heuristic rule. **c** (H01) Total number of separate locations where a specific heuristic rule corrected a merge error. **d** (H01) Total skeletal length eliminated by each heuristic rule.

**Fig. 11.**
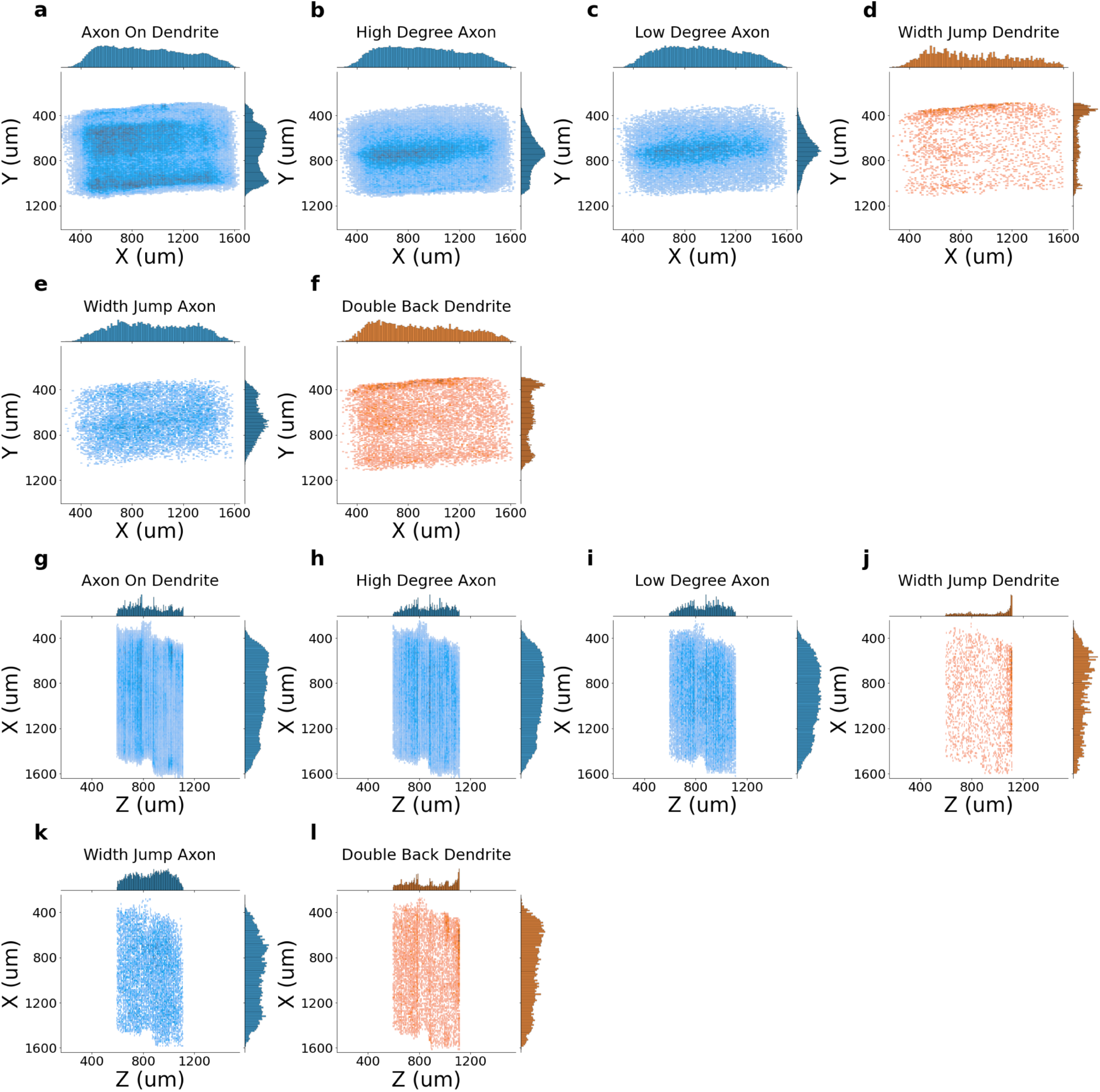
Spatial Distribution of MICrONS Merge Edit Locations. The distribution of locations show biases in the volume for certain types of merge edits. These spatial biases may be due to segmentation or slicing defects, or differences in the concentration of different kinds of neuropil throughout the volume. **a - f** X,Y merge edit locations for different heuristic rules **g - l** X,Z merge edit locations for different heuristic rules

**Fig. 12.**
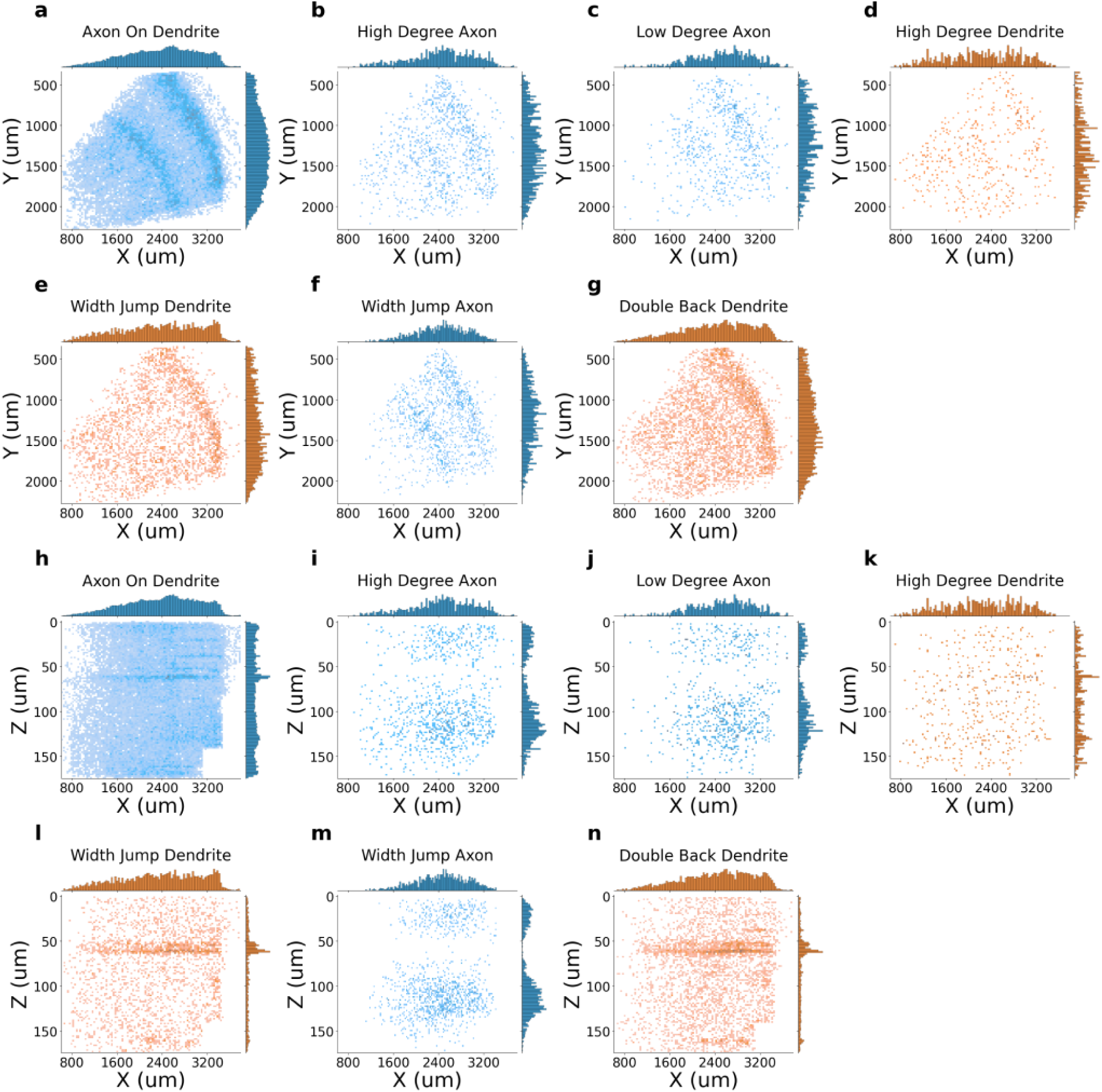
Spatial Distribution of H01 Merge Edit Locations. The distribution of locations show biases in the volume for certain types of merge edits. These spatial biases may be due to segmentation or slicing defects, or differences in the concentration of different kinds of neuropil throughout the volume. **a - g** X,Y merge edit locations for different heuristic rules **h - n** X,Z merge edit locations for different heuristic rules

**Fig. 13.**
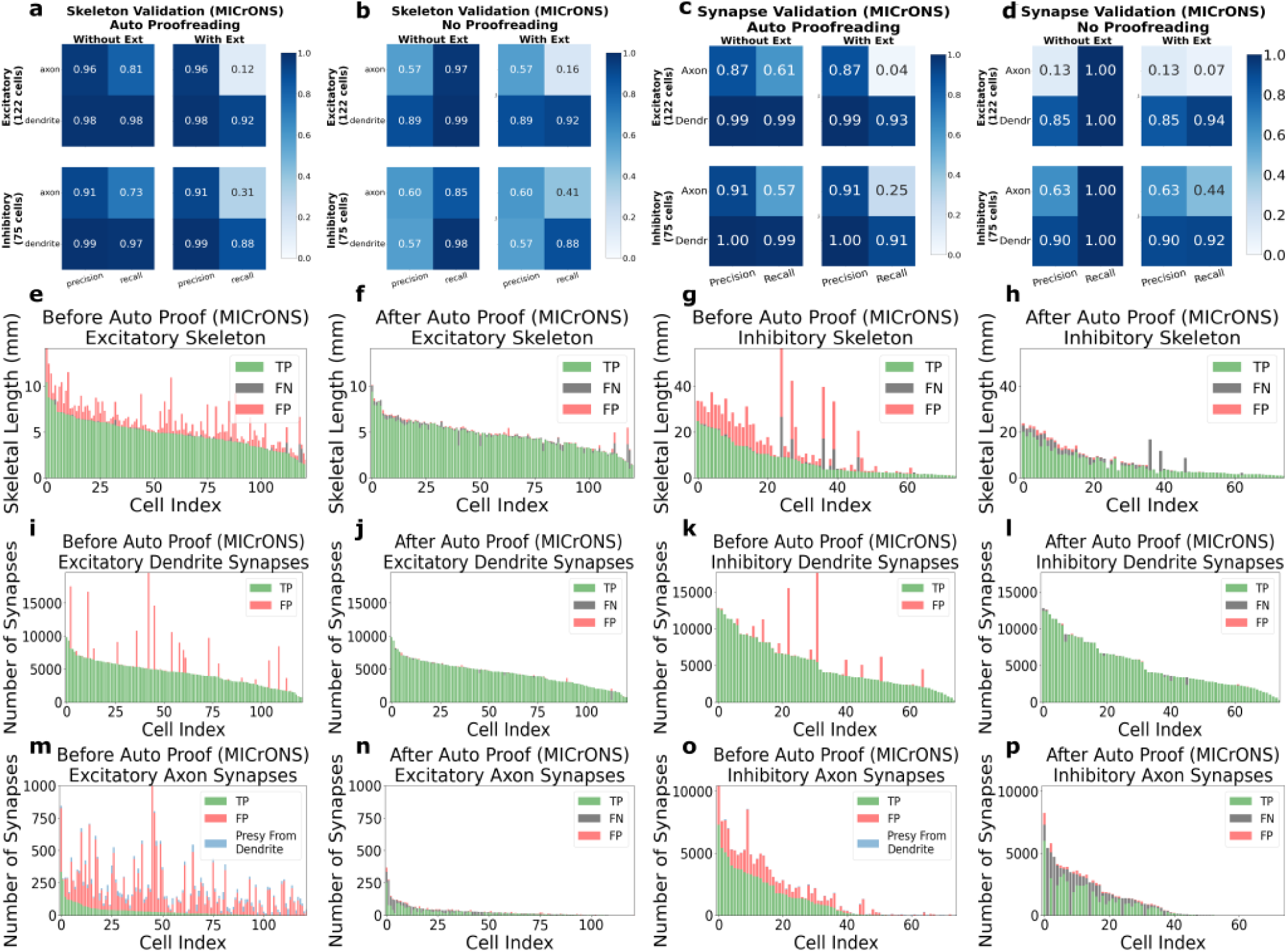
Supplemental MICrONS Auto Proofreading Validation. Validation metrics and visualizations of the automatic proof-reading step when comparing the edits made by the automatic proofreading algorithm to edits made by human proofreaders. The two different columns (“With Ext” and “Without Ext”) for the confusion matrices in panels a - d represent comparisons with two possible sources of human proofreading ground truth. The “With Ext” column refers to the skeleton or synapse state after automatic proofreading (panels a,c) or in the raw un-proofread segmentation (panels b,d) compared to the state after a human proofreader both cleaned the existing cell of merge errors and added back missing axon and dendrite segments. The “Without Ext” column performs this same comparison, but assuming that human proofreaders ONLY cleaned merge errors (without performing any extensions). The number of cells in the test set were 122 excitatory and 75 inhibitory. Histograms (panels e - p) give a visual representation of the metrics reported in the precision/recall tables (panels a - d). False Negative (FN) classifications can exist before automatic proofreading because of dropping axon/dendritic segments in the mesh and graph processing pipeline prior to the automatic proofreading step. Note: neurons with multi-soma merges are included in these visualizations and metrics. **a** The precision/recall metrics comparing the skeleton length of cells after automatic proofreading for the Axon/Dendrite compartments and for different exc/inh cell types when compared to human proofreading with extension and without. **b** The precision/recall metrics comparing the skeleton length of cells with no automatic proofreading. **c** The precision/recall metrics comparing the synapse counts of cells with automatic proofreading. **d** The precision/recall metrics comparing the synapse counts of cells with no proofreading. **e - h** TP/FN/FP classification of each test cell’s skeletons before and after automatic proofreading for both excitatory and inhibitory cells, demonstrating a large percentage of the FP skeleton segments are removed after the process. **i - l** TP/FN/FP classification of each test cell’s dendrite synapses (postsyns) before and after automatic proofreading for both excitatory and inhibitory cells, demonstrating a large percentage of the FP postsyns are removed after the application of dendrite proofreading heuristics. **m - p** TP/FN/FP classification of each test cell’s axon synapses (presyn) before and after automatic proofreading for both excitatory and inhibitory cells, demonstrating a large percentage of the FP presyns are removed after the application of axon proofreading heuristics. Those axon presyns located not on the main axon but on dendritic segments are filtered away and designated as “Presyn From Dendrite”, which does not include the heuristic rule of “Axon on Dendrite” but instead just filters away any presyns located on dendritic segments that were not filtered away using the heuristic rules.

**Fig. 14.**
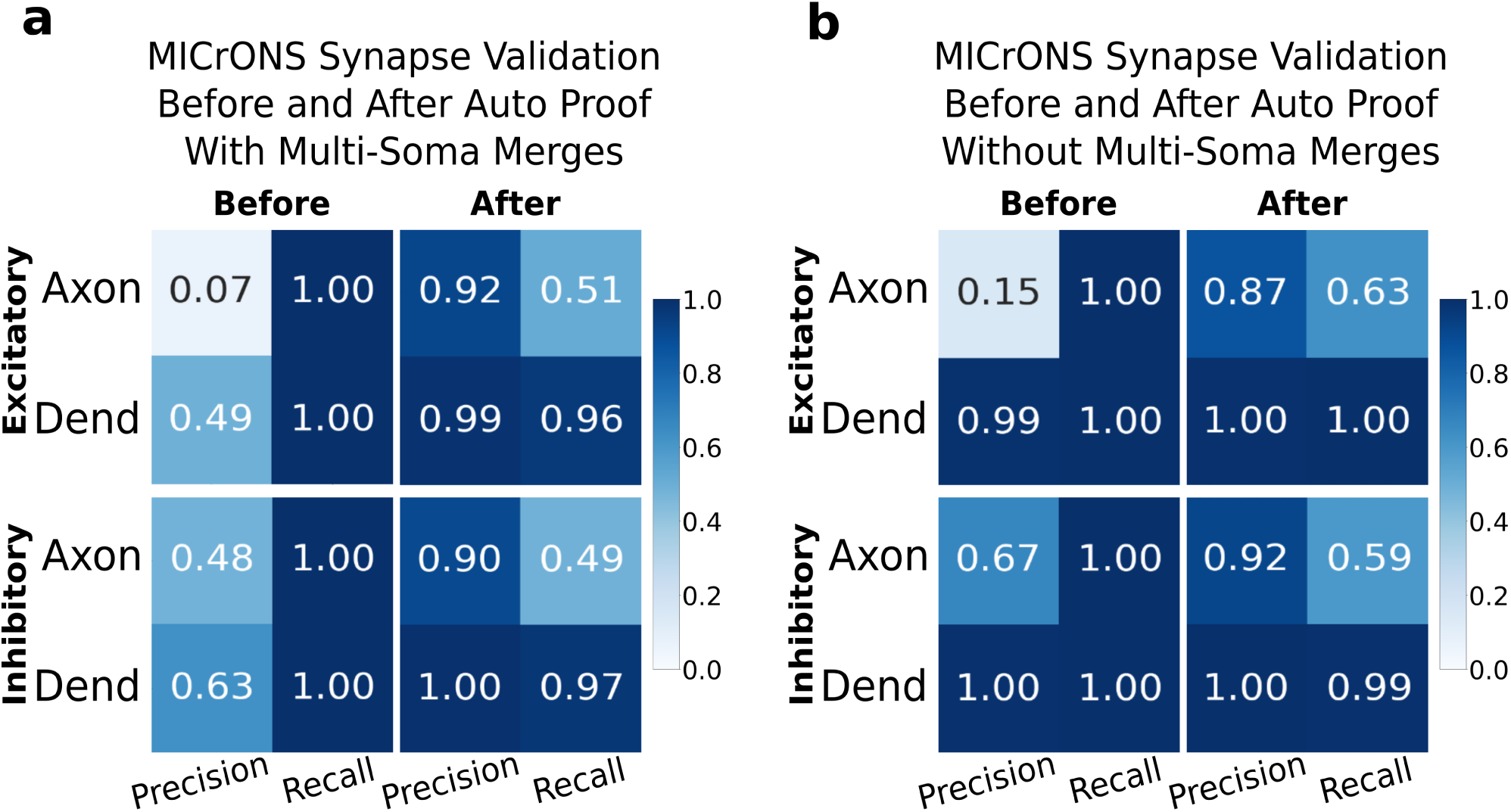
Supplemental MICrONS Multi-Soma Auto Proofreading Validation. The precision/recall metrics comparing the synapse counts of cells before and after automatic proofreading for the Axon/Dendrite compartments and for different excitatory/inhibitory cell types when compared to human proofreading. **a** Validation when only considering neurons with at least one soma merge to the main segment (19 excitatory, 12 inhibitory). **b** Validation when only considering neurons with no soma merged to the main segment (103 excitatory, 63 inhibitory).

**Fig. 15.**
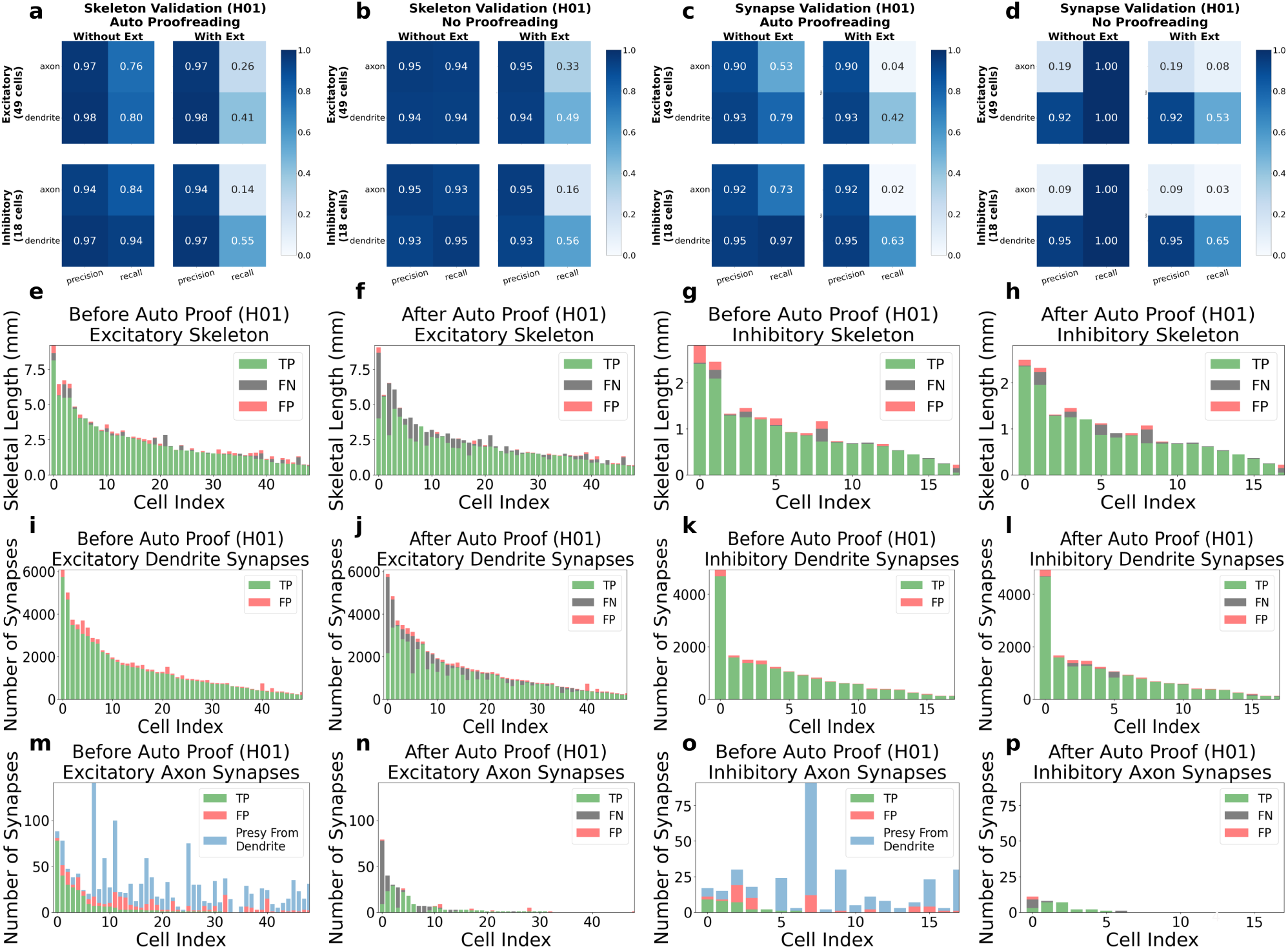
Supplemental H01 Auto Proofreading Validation. Validation metrics and visualizations of the automatic proofreading step when comparing the edits made by the automatic proofreading algorithm to edits made by human proofreaders. The two different columns (“With Ext” and “Without Ext”) for the confusion matrices in panels a - d represent comparisons with two possible sources of human proofreading ground truth. The “With Ext” column refers to the skeleton or synapse state after automatic proofreading (panels a,c) or in the raw un-proofread segmentation (panels b,d) compared to the state after a human proofreader both cleaned the existing cell of merge errors and added back missing axon and dendrite segments. The “Without Ext” column performs this same comparison, but assuming that human proofreaders ONLY cleaned merge errors (without performing any extensions). The number of cells in the test set were 49 excitatory and 18 inhibitory. Histograms (panels e - p) give a visual representation of the metrics reported in the precision/recall tables (panels a - d). FN classifications can exist before automatic proofreading because of dropping axon/dendritic segments in the mesh and graph processing pipeline prior to the automatic proofreading step. Note: While perfectly extending all axonal and dendritic processes is not yet possible, the extent to which neurons were extended in the manually proofread set from the H01 dataset are much less extensively extended in comparison to those of the MICrONS dataset; therefore, the recall numbers for the “With Ext” categories in the H01 validation are much more likely an over-estimate in comparison with those of the MICrONS dataset. **a** The precision/recall metrics comparing the skeleton length of cells after automatic proofreading for the Axon/Dendrite compartments and for different exc/inh cell types when compared to human proofreading with extension and without. **b** The precision/recall metrics comparing the skeleton length of cells with no automatic proofreading. **c** The precision/recall metrics comparing the synapse counts of cells with automatic proofreading. **d** The precision/recall metrics comparing the synapse counts of cells with no proofreading. **e - h** TP/FN/FP classification of each test cell’s skeletons before and after automatic proofreading for both excitatory and inhibitory cells, demonstrating a large percentage of the FP skeleton segments are removed after the process. **i - l** TP/FN/FP classification of each test cell’s dendrite synapses (postsyns) before and after automatic proofreading for both excitatory and inhibitory cells, demonstrating a large percentage of the FP postsyns are removed after the application of dendrite proofreading heuristics. **m - p** TP/FN/FP classification of each test cell’s axon synapses (presyn) before and after automatic proofreading for both excitatory and inhibitory cells, demonstrating a large percentage of the FP presyns are removed after the application of axon proofreading heuristics. Those axon presyns located not on the main axon but on dendritic segments are filtered away and designated as “Presyn From Dendrite”, which does not include the heuristic rule of “Axon on Dendrite” but instead just filters away any presyns located on dendritic segments that were not filtered away using the heuristic rules.

**Fig. 16.**
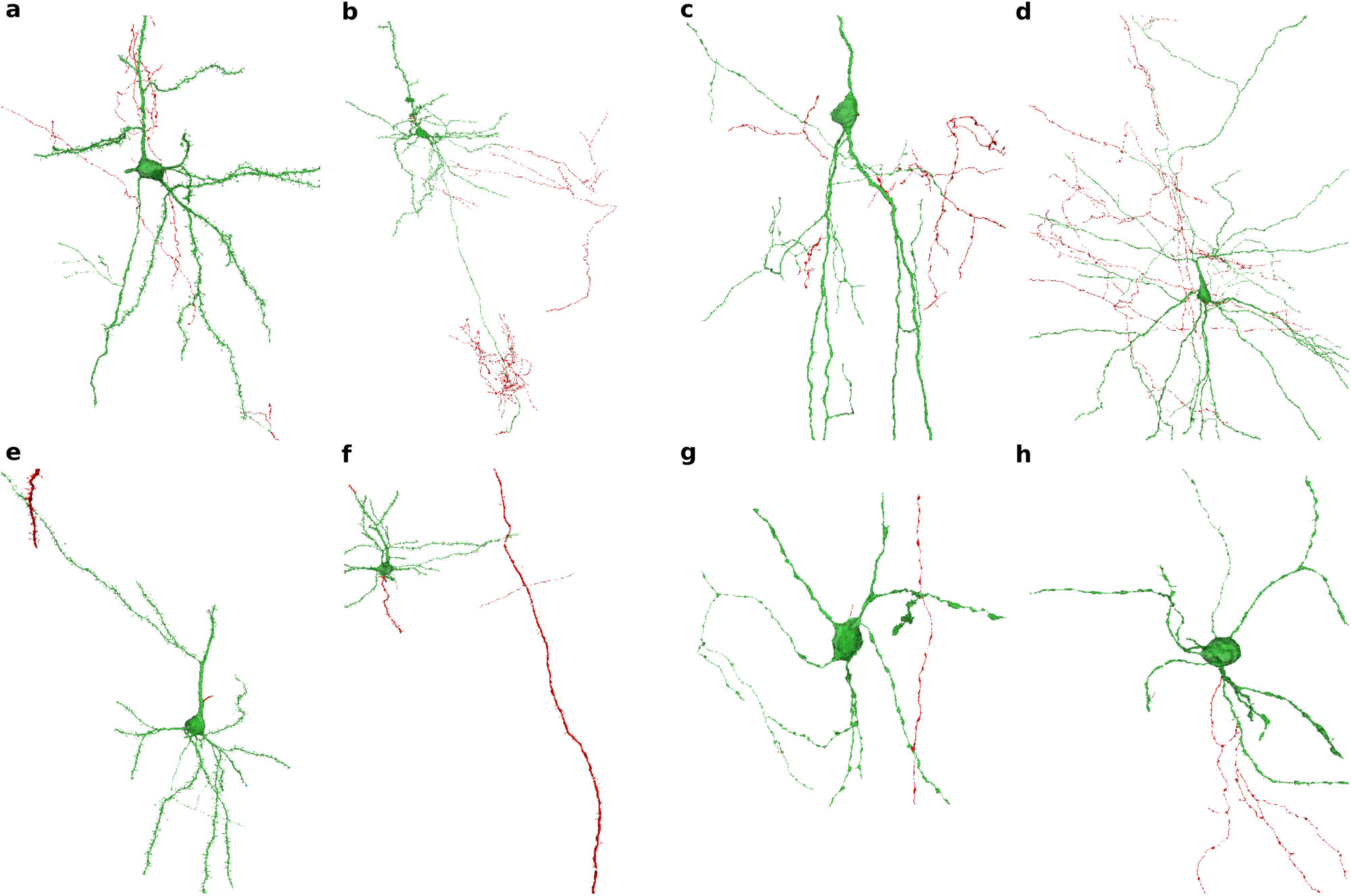
Proofread Neuron Examples with Merge Errors Labeled. Examples of excitatory and inhibitory neurons from both the MICrONS and H01 dataset after automatic proofreading (green) with the removed merge errors shown (red). **a,b** (MICrONS) Example excitatory cells in the 50th and 90th percentile of merge error skeletal length removed. **c,d** (MICrONS) Example inhibitory cells in the 50th and 90th percentile of merge error skeletal length removed. **e,f** (H01) Example excitatory cells in the 50th and 90th percentile of merge error skeletal length removed. **g,h** (H01) Example inhibitory cells in the 50th and 90th percentile of merge error skeletal length removed.

**Fig. 17.**
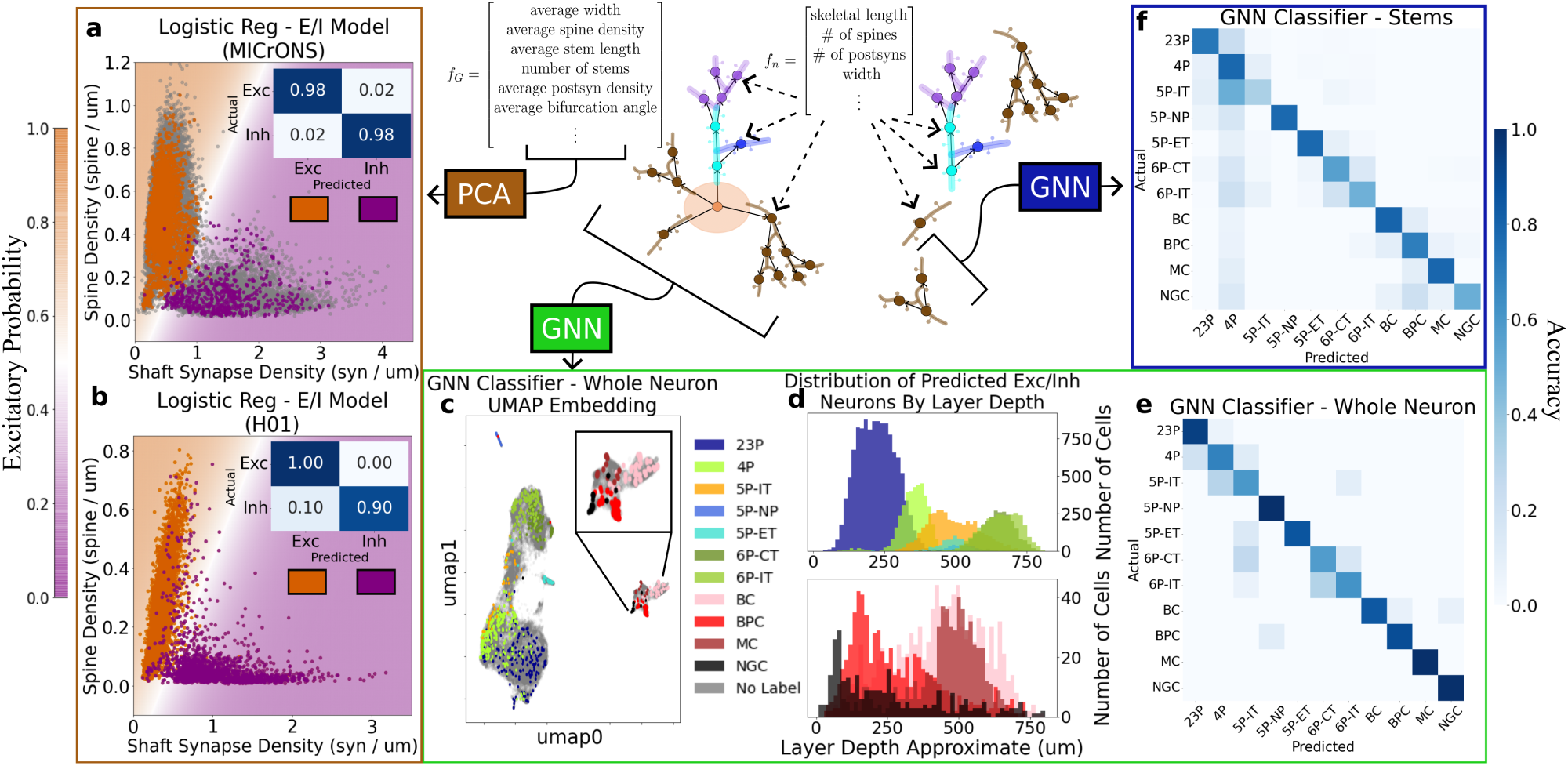
Graph decomposition enables cell-type classification. **a,b** Two interpretable features identified by PCA were highly informative for excitatory/inhibitory classification: spine density (number of spines per *µ*m of skeletal length) and shaft synapse density (number of synapses not on a spine per *µ*m of skeletal length). Consistent with previous studies (Azouz et al., 1997; Kawaguchi et al., 2006), a logistic regression model trained on just these two features enabled linear discrimination of excitatory and inhibitory cells with high accuracy (same parameters for logistic regression model used for both MICrONS and H01 dataset). **c** A Graph Convolutional Neural Network (GCN) trained on manually-annotated cell types produces an embedding space with a continuum of excitatory neurons progressing from the top layers down to the bottom layers while keeping inhibitory neurons and some excitatory neurons with distinct morphology (5P-NP and 5P-ET) clearly separated (see Table 1 for the cell-type abbreviation glossary). **d** The depth of predicted cell types outside of the training volume are consistent with their expected laminar distribution even though no coordinate features are used in the GCN classifier. **e** Confusion Matrix of the test dataset for the neuron GCN classifier tested on n=178 held out neurons: 23P (n=33), 4P (n=51), 5P-IT (n=10), 5P-NP (n=4), 5P-ET (n=7), 6P-CT (n=19), 6P-IT (n=29), BC (n=13), BPC (n=9), MC (n=1), NGC (n=2). **f** Cell classes could also be determined using a dendritic subgraph of only one stem (branching segment connected to the soma) nearly as well as when using the entire dendritic tree, suggesting that the GCN identifies somewhat local features that enable classification. GCN tested on n=1023 test stems: 23P (n=230), 4P (n=301), 5P-IT (n=48), 5P-NP (n=18), 5P-ET (n=45), 6P-CT (n=116), 6P-IT (n=137), BC (n=65), BPC (n=30), MC (n=19), NGC (n=14).

**Fig. 18.**
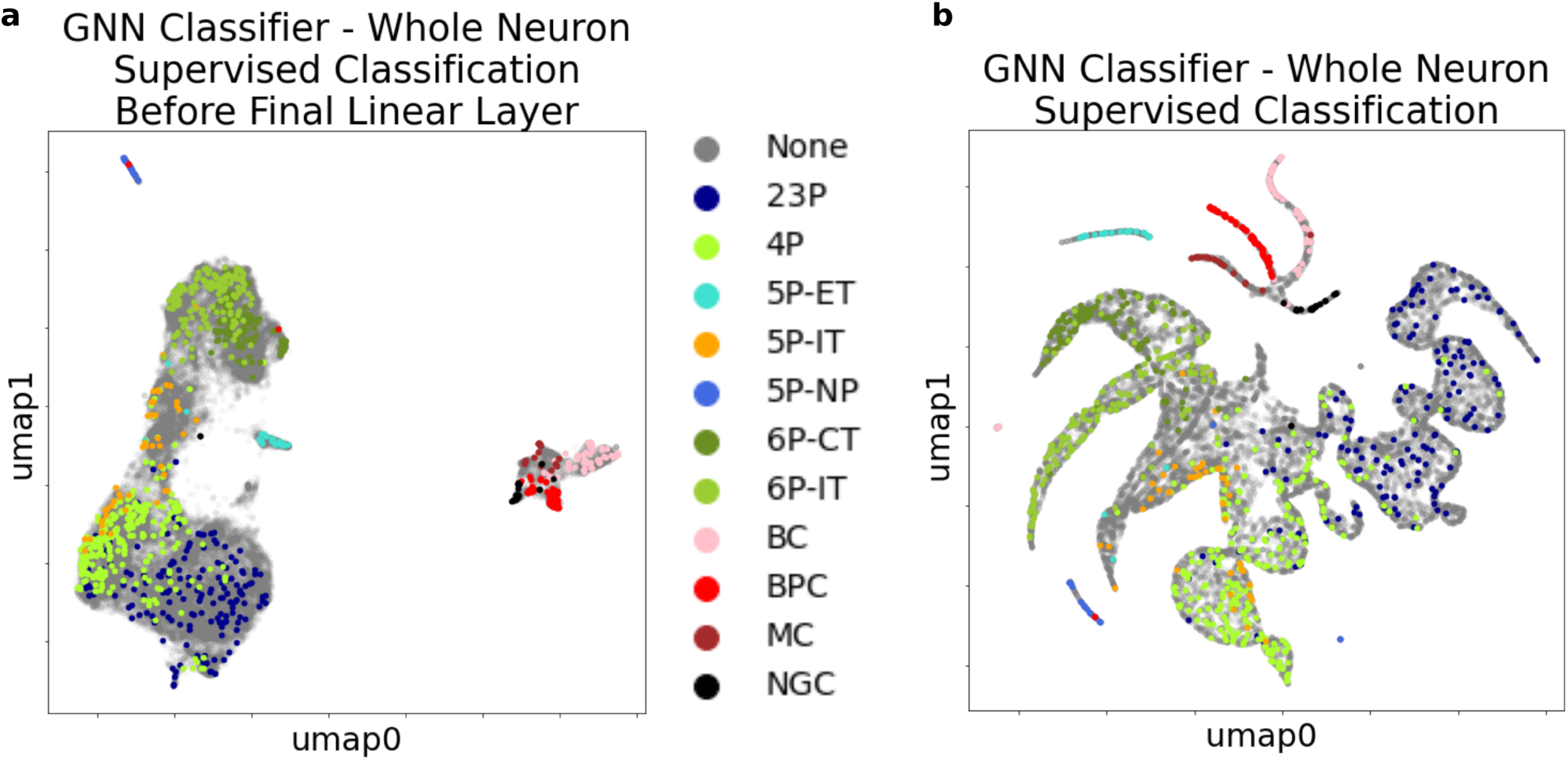
GNN Classifier Whole Neuron UMAP Embeddings. **a** Embeddings before the final linear layer and softmax function with hand labeled cells from (Schneider-Mizell et al., 2023) overlaid (these labels were used for the training and validation process of the GNN). Cell-type separation is evident at this stage indicating that the classifier has learned useful features prior to the readout. See Table 1 for the cell-type abbreviation glossary. **b** Embeddings after the final linear layer with cell type labels from (Schneider-Mizell et al., 2023).

**Fig. 19.**
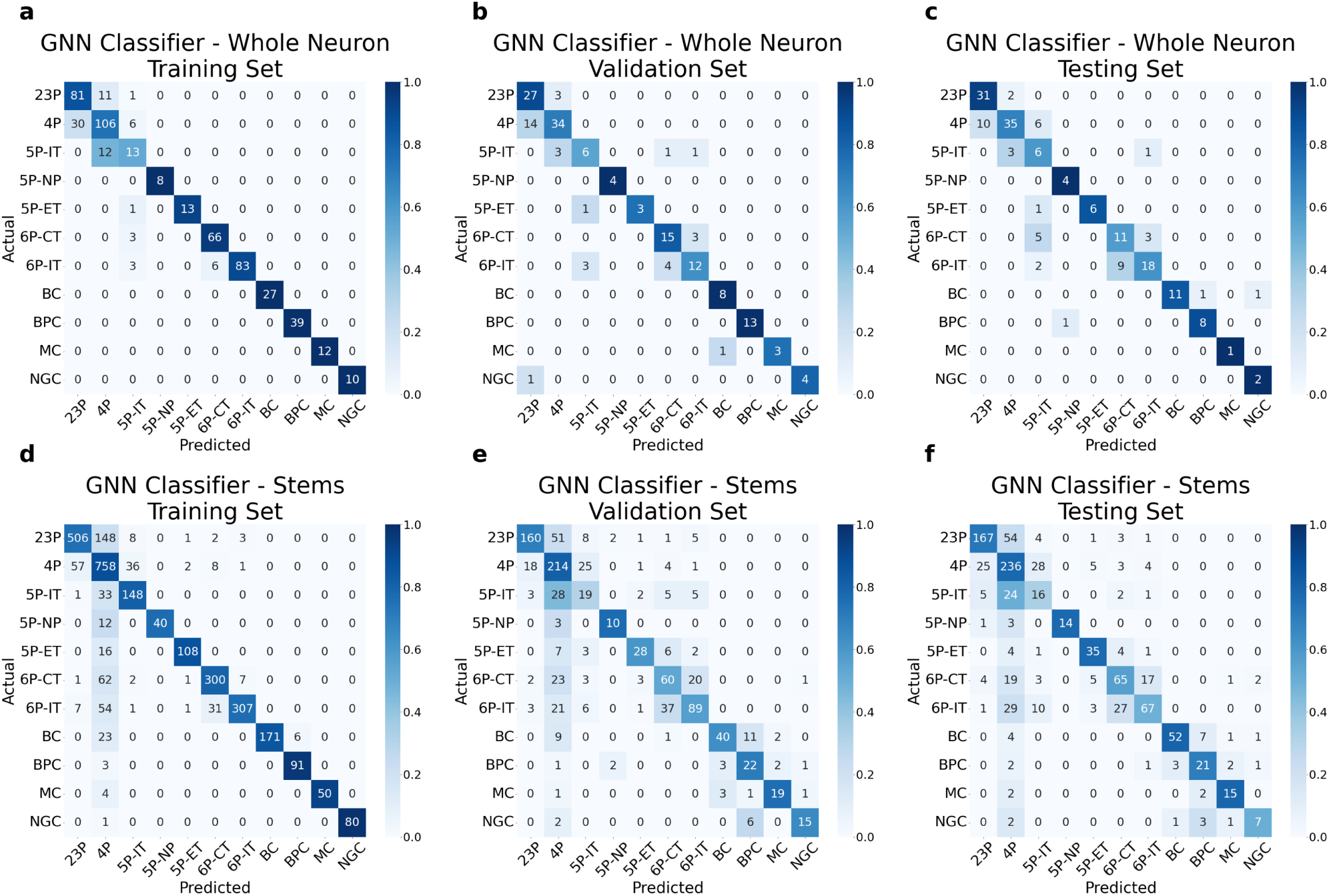
GNN Classifier Train/Validation/Test Confusion. Confusion matrix of predicted and actual neuron counts from each of the 60:20:20 Train/Validation/Test splits used in the supervised training process of the Graph Neural Network cell type classifier. All ground truth labels were from hand-annotated classes described in (Schneider-Mizell et al., 2023). Color intensities are from normalized values in reference to the summation of a given row. See Table 1 for the cell-type abbreviation glossary. **a-c** Confusion matrix (neuron counts) for the GNN Classifier using the entire neuron graph including the somatic root node. **d-f** Confusion matrix (stem counts) for the GNN Classifier applied to a single stem (a single projection from the soma), without any information about the soma itself.

**Fig. 20.**
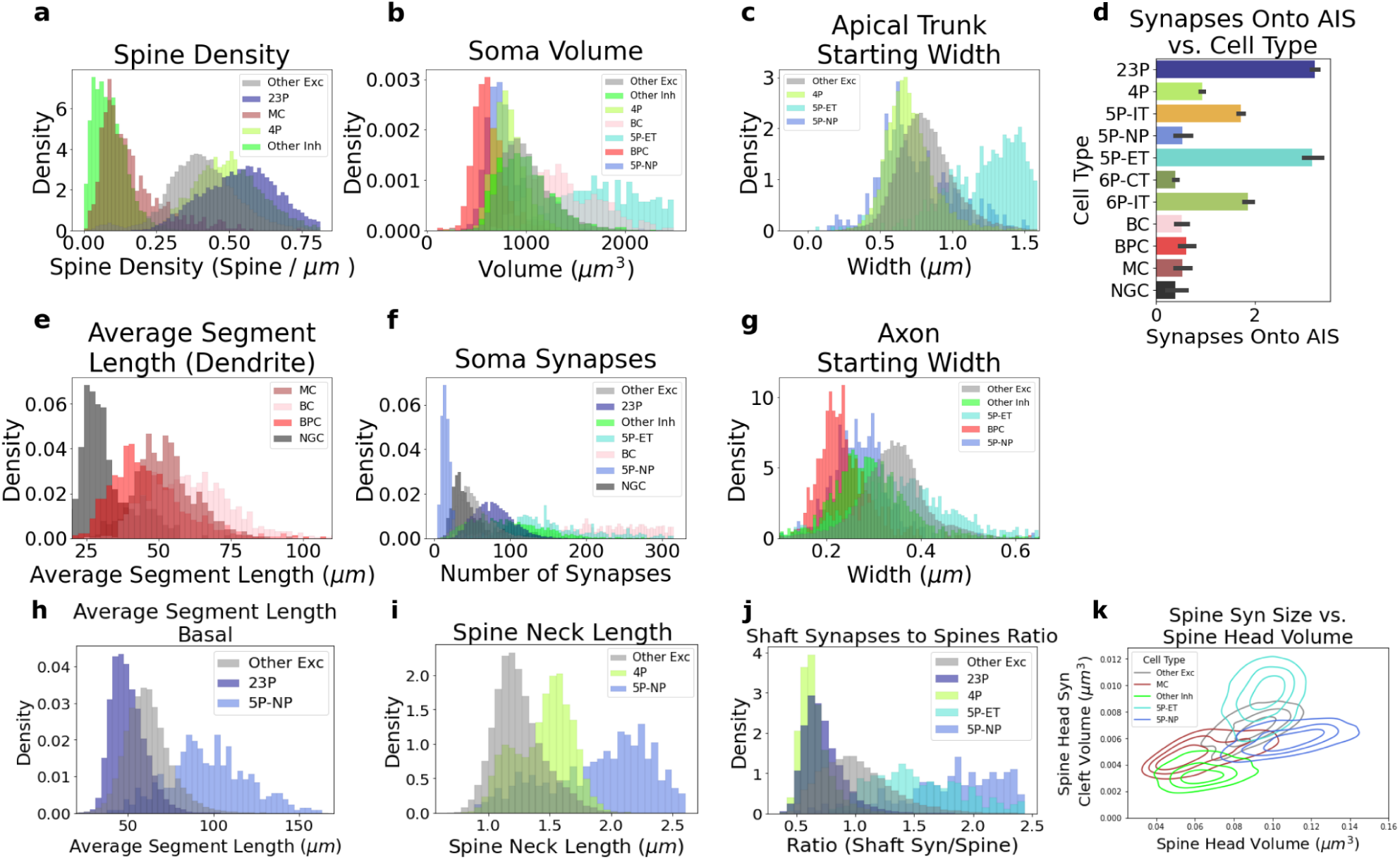
Various morphological features computed by NEURD. Histograms and bar graphs of a variety of salient features computed by NEURD, colored by the labels generated from the GNN classifier. Some plots are replicating previous work from (Elabbady et al., 2022). See Table 1 for the cell-type abbreviation glossary. **a** Spine density (number of spines per *µ*m of skeletal length) distributions from automatic spine detection. As has been previously reported (Azouz et al., 1997), layer 2/3 pyramidal cells are more densely spiny than layer 4, and MC spine density is higher than other inhibitory cells. **b** Soma volume computed during the soma detection step in the NEURD mesh processing pipeline. As expected, 5P-NP, 4P and BPC generally have smaller somas than other cells from their same excitatory or inhibitory class while 5P-ET and BC are larger than other cells in the same class (Elabbady et al., 2022). **c** Width measurements generated from the average distance of the inner skeleton to the mesh surface (radius approximation) at the beginning of the apical trunk protrusion. Compared to other cell types, 4P and 5P-NP cells have smaller trunks, while 5P-ET are larger. **d** Average number of synapses onto the axon initial segment (AIS) for different cell types. As expected, 23P, 5P-ET, and 5P-IT cell types are more densely innervated on their AIS (Schneider-Mizell et al., 2021). AIS is defined as within 10 *µ*m - 40 *µ*m skeletal distance of the soma, and error bars are standard deviation. **e** Average skeletal length of non-branching dendritic segments for stems of different cell type subclasses, illustrating that NGC have significantly shorter distances between branch points in their dendrites than other inhibitory cells. **f** Distributions of synapses onto the soma illustrating the expected larger average number of soma synapses for 5P-ET and BC and smaller numbers for 5P-NP and NGC (Elabbady et al., 2022). **g** Distributions of radius approximation for the start of the axon protrusion from either a dendrite or the soma, showing smaller typical widths for 5P-NP and BPC and larger starting widths for 5P-ET. **h** Distribution of the average skeletal length of non-branching dendritic segments for stems of different cell-type subclasses. **i** Distribution of average spine neck length of different excitatory cell type subclasses. As expected, 5P-NP cells have the longest average neck lengths, but 4P cells also display significantly longer necks than other excitatory cell types. **j** The ratio of non-spine synapses to spine counts varies across cell types. **k** KDE plots (kernel density estimation, estimates the shape and distribution of the discrete dataset, quartile levels shown) of spines on different cell type subclasses, comparing each spine head’s volume and the size of the largest synapse on that spine head. These plots reveal differences in both the distribution and scaling of synapse sizes and spine heads across cell types. For example Martinotti cells (red) have larger spine head synapses than other inhibitory spines (light green) and the head volume and synapse size scale at a rate more similar to other excitatory cells (grey).

**Fig. 21.**
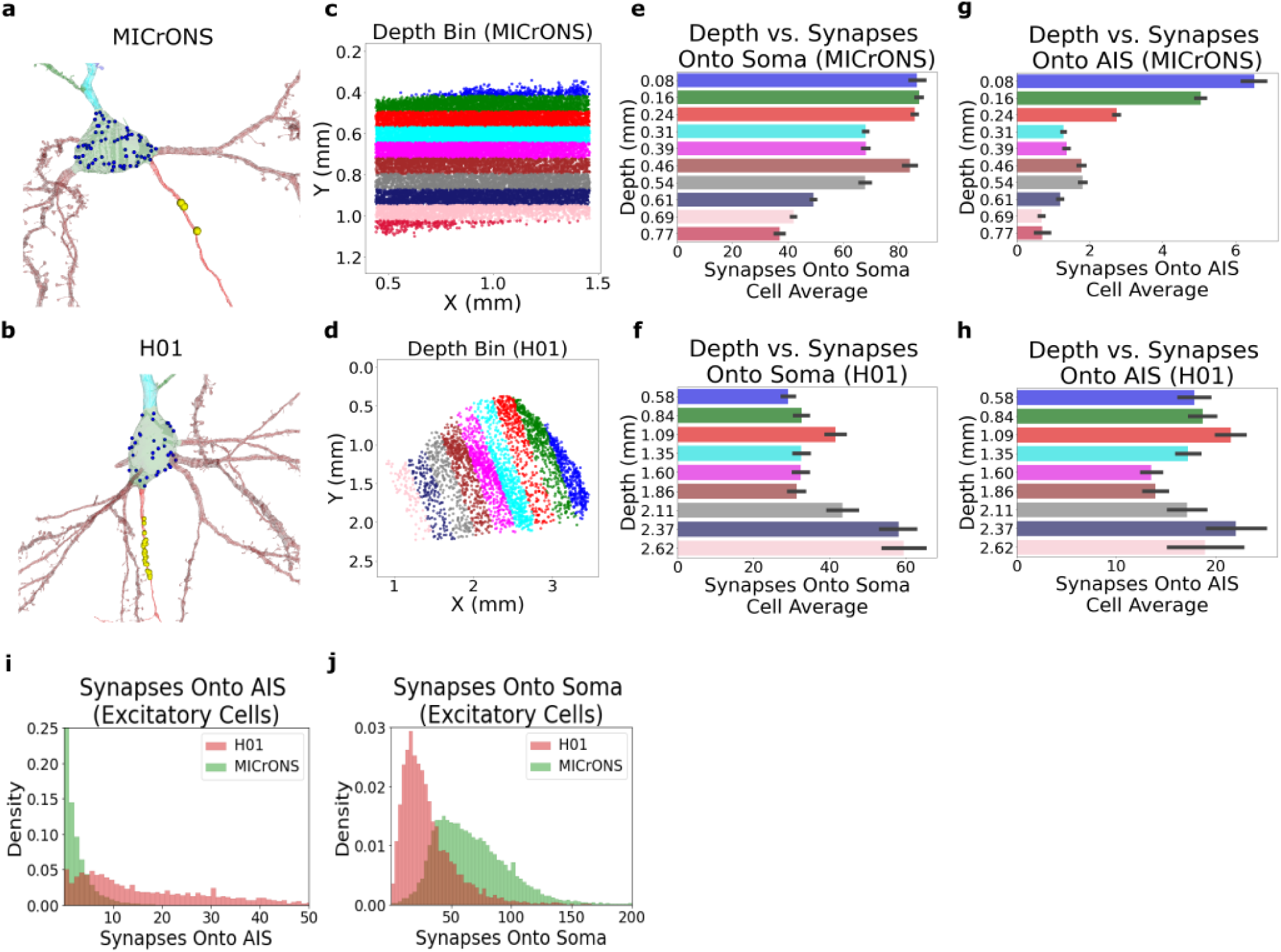
Comparison of Synapses onto AIS and Soma. **a-b** Example neurons with synapses onto the axon (AIS synapses) in yellow and synapses onto soma in blue. Example neurons are both in the 75th percentile of the number of AIS synapses in their respective volumes. **c,d** Depth bins used for analysis of both synapses onto AIS vs depth and synapses onto soma vs. depth (this figure, panels e-h and Main Fig. 4a). Note, for the MICrONS dataset plots there is an offset of approximately 300 *µ*m between the depth value and the y coordinate. **e,f** Average number of synapses onto the soma of cells varies across depth (mean +/- std), decreasing in deeper layers of the MICrONS volume, but increasing in deeper layers of the H01 dataset. **g,h** Average number of synapses onto the axon initial segment (AIS) of cells at different laminar depths (mean +/- std) for the MICrONS and H01 volume. **i** Distribution of the number of AIS synapses per cell compared across datasets, emphasizing the increased innervation of the AIS for neurons in the H01 dataset in comparison to MICrONS. **j** Distribution of the number of soma synapses per cell. As expected, neurons in the MICrONS volume have more identified synapses onto their soma, despite the smaller surface area of mouse somas compared to human (Wildenberg et al., 2021).

**Fig. 22.**
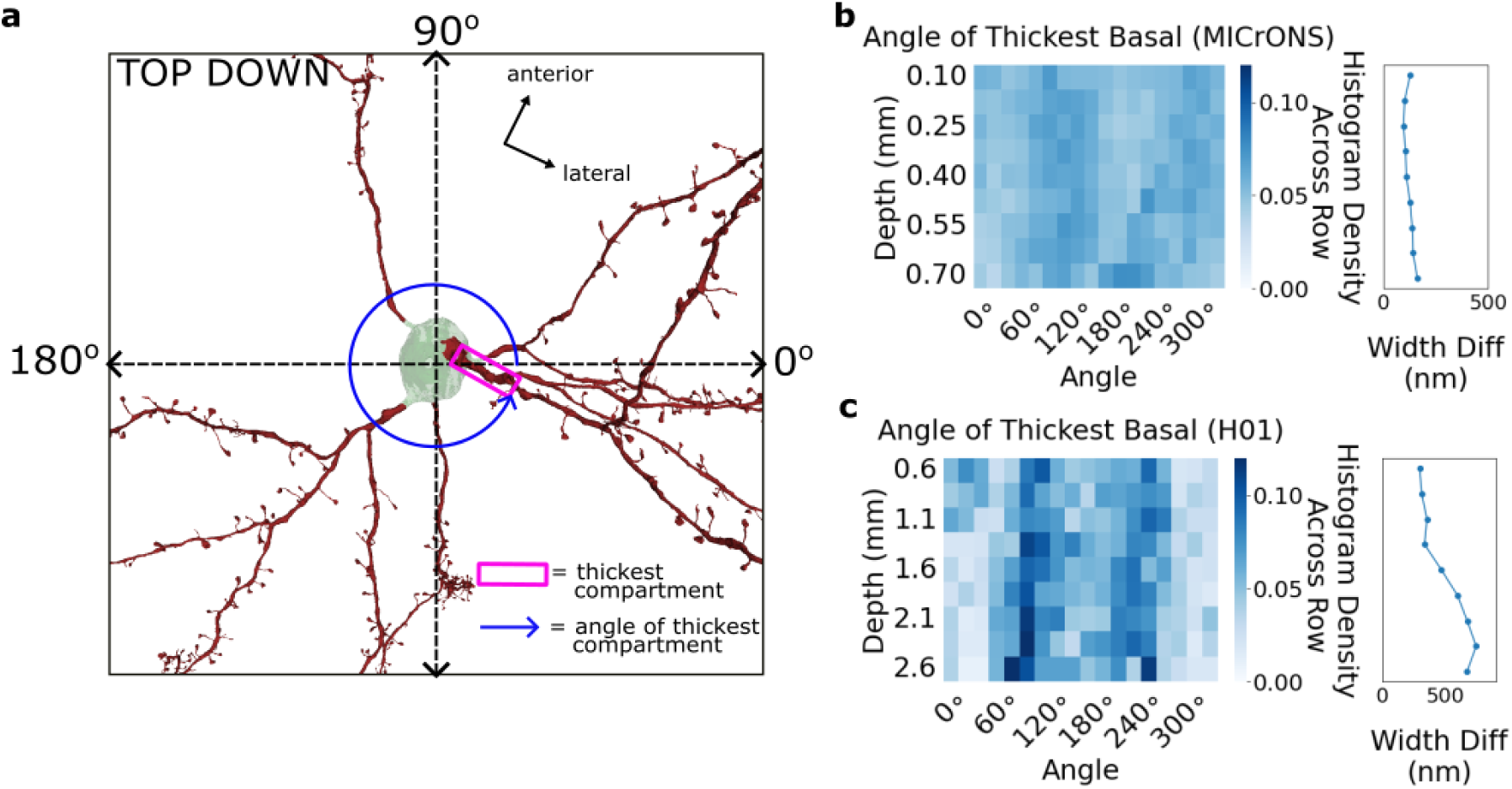
Thickest Basal Geometric Analysis. Using the easily accessible geometric features of neurons after decomposition, the thickest basal branch is identified and the distribution of xz projection angles for all neurons in the volume is compared across depths for the two volumes. **a** Top down view of a neuron to illustrate the geometric analysis: the thickest basal branch is boxed in pink and the xz angle of these branches are indicated with a blue circular angle marker. **b,c** Histograms showing the distribution of mean skeletal angle of the thickest basal stem. Each row is a normalized histogram for a specific depth bin. The H01 dataset shows more bimodal structure (especially in the deeper layers) than the MICrONS dataset, consistent with a previous report (Shapson-Coe et al., 2021). We find that this pattern is also visible in more superficial layers of H01, but is less obvious because the width difference between the widest and the second-widest basal branch is much larger in deep layers (“Width Diff” plots to the right of the heat maps).

**Fig. 23.**
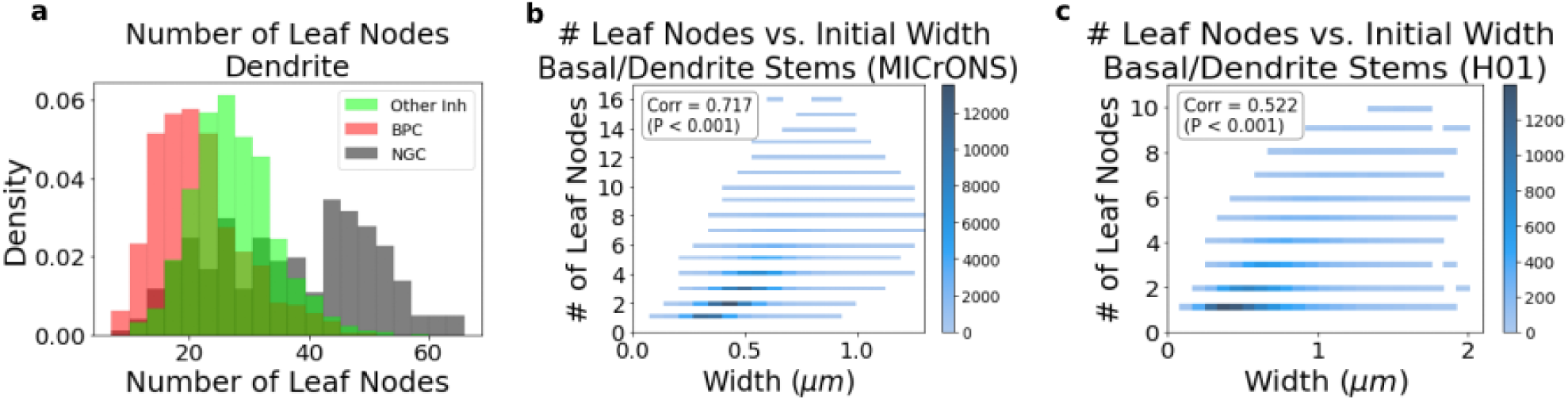
Neuron Dendritic Branching Characteristics. Measurements related to leaf nodes (terminating ends of the dendritic stem) excluding apical dendrites. **a** Distributions of the number of total leaf nodes for the non-apical dendrites of each neuron separated by inhibitory cell type. As expected, NGC cells have the most leaf nodes of any inhibitory cell type, while BPC have fewer leaf nodes compared to other interneurons. **b-c** Histogram for all the non-apical dendritic stems of every neuron in the volume comparing the initial width of the stem to the number of leaf nodes. For both the MICrONS (b) and H01 (c), there is a significant positive correlation.

**Fig. 24.**
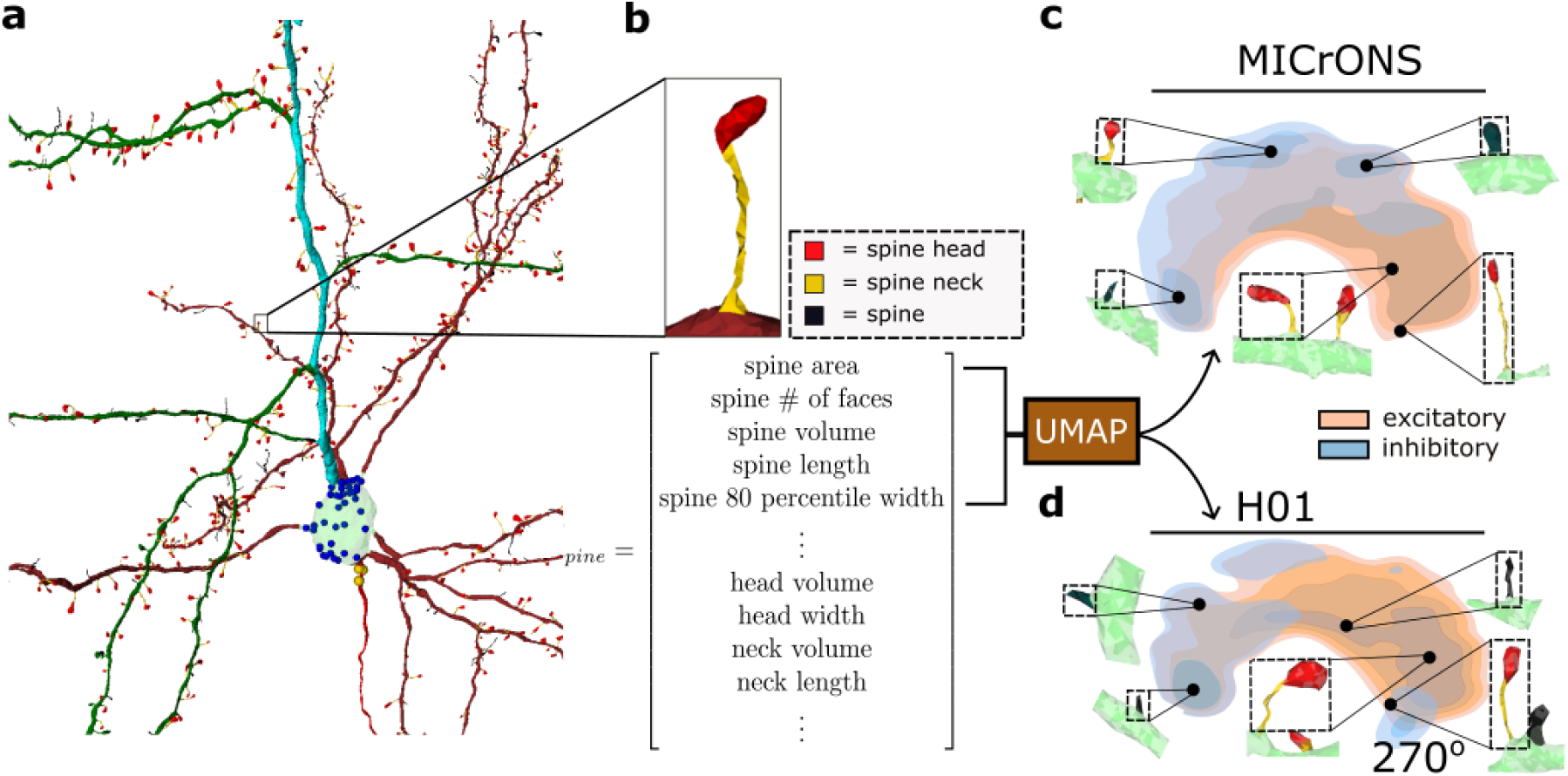
Spine Feature Extraction and UMAP distribution. **a** Cleaned and annotated neuron mesh; soma synapses in blue, axon initial segment (AIS) synapses in yellow, basal dendrite in brown, apical trunk in aqua, oblique branches in green. Spine heads in red, spine necks in yellow and non-segmented spine in black along dendritic segments. An random spine is identified with the black rectangle and expanded in the next panel. **b** Example spine submesh from the spine identification with the head and neck mesh segmentation shown and a vector of features extracted from the spine submesh. Most spines are annotated with interpretable features such as head volume, spine skeletal length, and spine neck length while some spines (shown in black) which are smaller (typically under 0.7 um) or unable to be segmented, lack the head/neck features. **c,d** Kernel density estimation of UMAP embedding (quartile levels shown) of spines sampled from MICrONS and H01 dataset using spine features from panel **b** (without head or neck features). The embeddings show a similar embedding structure between the two datasets in terms of spine shape and inhibitory/excitatory class, similar to previous work clustering a non-parametric representation of postsynaptic shapes (Seshamani et al., 2020).

**Fig. 25.**
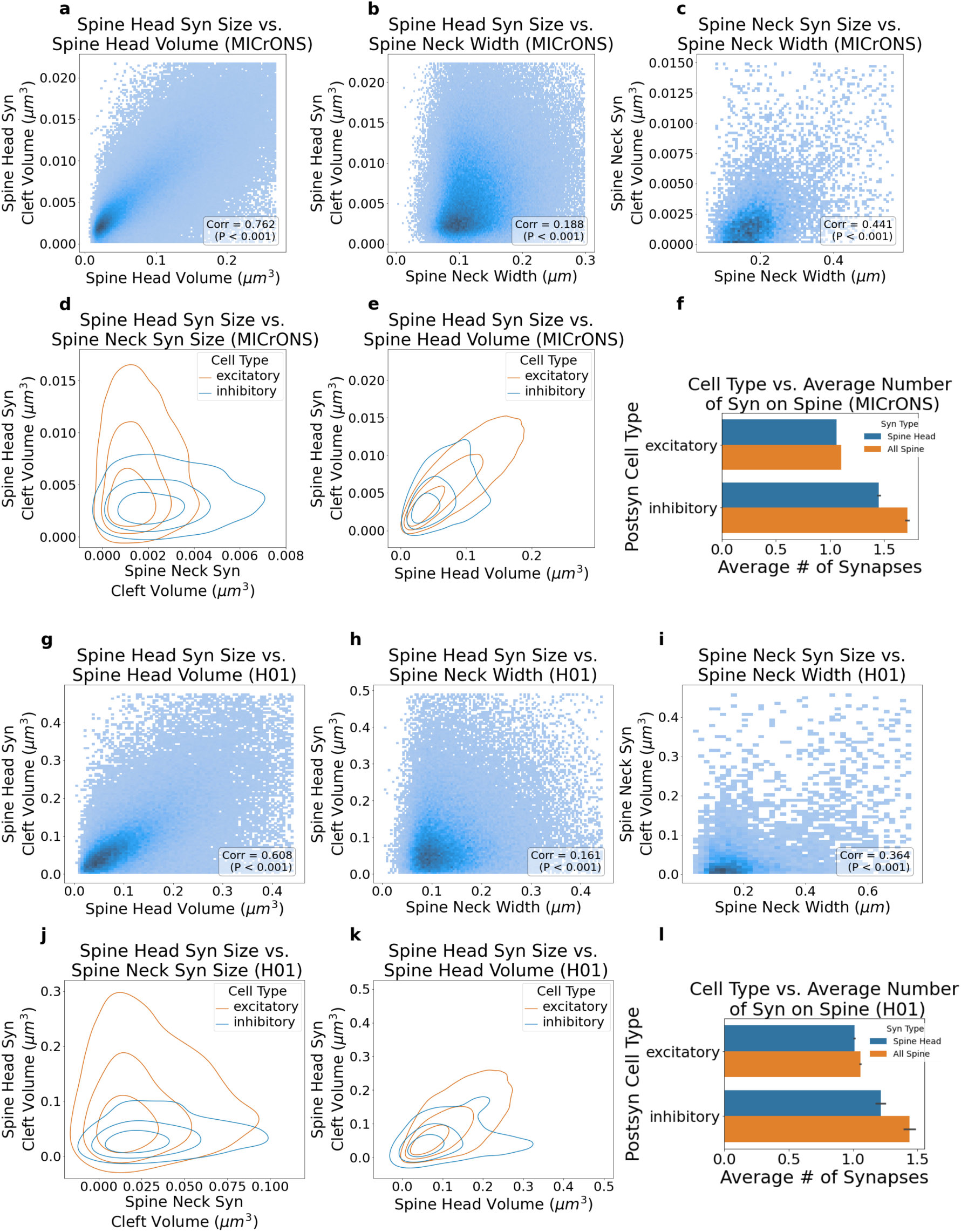
Postsynaptic Spine Feature Analysis. Here we compare the distributions and correlations of certain spine features; replicating and expanding on previous work. The MICrONS dataset is analyzed in **a-f** and the same analysis is repeated for H01 in **g-l**. **a-b,g-h** As expected, for synapses onto the spine **head**, the size of the synaptic cleft and the volume of the spine head mesh are strongly positively correlated, while cleft size and neck width are positively but more weakly correlated (Harris and Stevens, 1989; Arellano et al., 2007). **c,i** For synapses onto the spine **neck**, the width of the spine neck and the synaptic cleft volume of synapses are positively correlated. **d,j** KDE of the joint distribution (quartile levels shown) of the spine neck synaptic cleft volume with spine head synaptic cleft volume for different postsynaptic cell types (exc/inh), illustrating the different joint distributions for each cell type. For synapses onto spine **heads**, synaptic size has a wider range for excitatory cell spines than inhibitory cell spines in both volumes. For synapses onto spine **necks**, the range of synaptic size is larger for inhibitory cells in the MICrONS volume, but similar in the H01 dataset. **e,k** Spine head volume is positively correlated with spine head synaptic cleft volume for both excitatory and inhibitory neurons in both datasets. **f,l** Average number of synapses on all spines and spine heads for different cell types, indicating that in both datasets inhibitory spines receive more synapses per spine than excitatory spines.

**Fig. 26.**
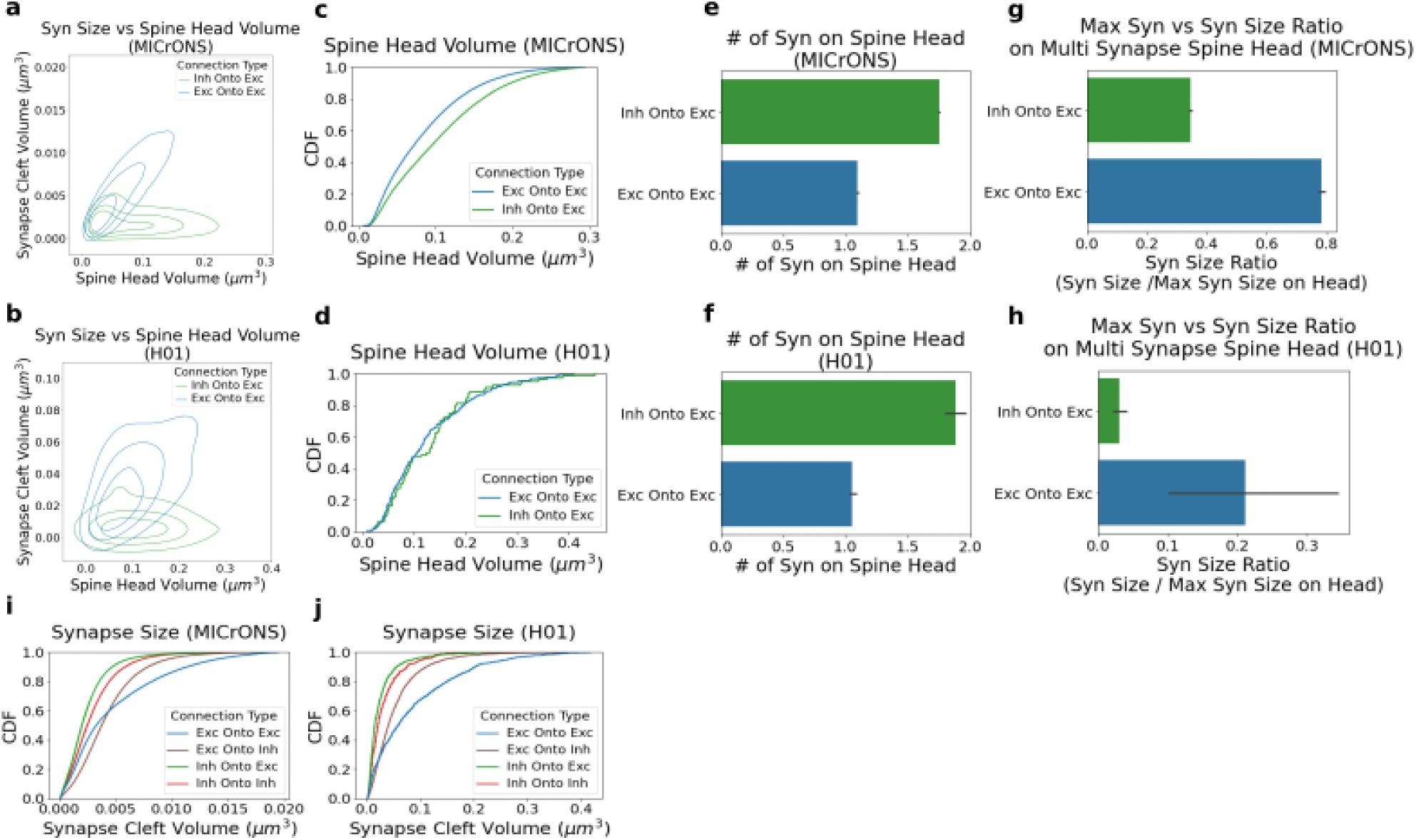
Spine and Synapse Connectivity Analysis. We revisited the spine analysis in Fig. 25 taking into account information about the identity of the presynaptic neuron for each synapse. **a-b** KDE distribution (quartile levels shown) relating the postsynaptic spine head volume and the synapse cleft volume for synapses onto excitatory cells given different presynaptic cell types. In both datasets a significant positive correlation between spine head volume and synapse size is observed only when the source cell is excitatory but not when the source is inhibitory. **c-d** CDF of the spine head volume for postsynaptic excitatory cells given different presynaptic cell types. For the MICrONS dataset, inhibitory presynaptic cells typically target larger spine heads but this trend is not significant in H01. **e-f** As a possible explanation of why inhibitory cells target larger spine heads, a plot of the average number of synapses on a spine head conditioned on the presynaptic cell type shows that spine heads targeted by inhibitory neurons generally have two synapses as opposed to a mean closer to one synapse per spine for excitatory synapses. **g,h** Expanding on the observation that inhibitory cells typically synapse onto spines with more than one synapse, for spines with multiple synapses, we plot the relative size of a spine head synapse to the size of the largest synapse on that same spine head (mean +/- std) given different presynaptic cell types. We observe that the synapse from an inhibitory source is typically much smaller than the largest synapse on the spine head in both the MICrONS and H01 dataset. **i,j** CDF of the distribution of synapse cleft volumes for different connections types show a similar trend between the MICrONS and H01 dataset where the synapses with excitatory presynaptic cells are typically larger than inhibitory cells and synapses onto inhibitory cells are typically smaller than those onto excitatory.

**Fig. 27.**
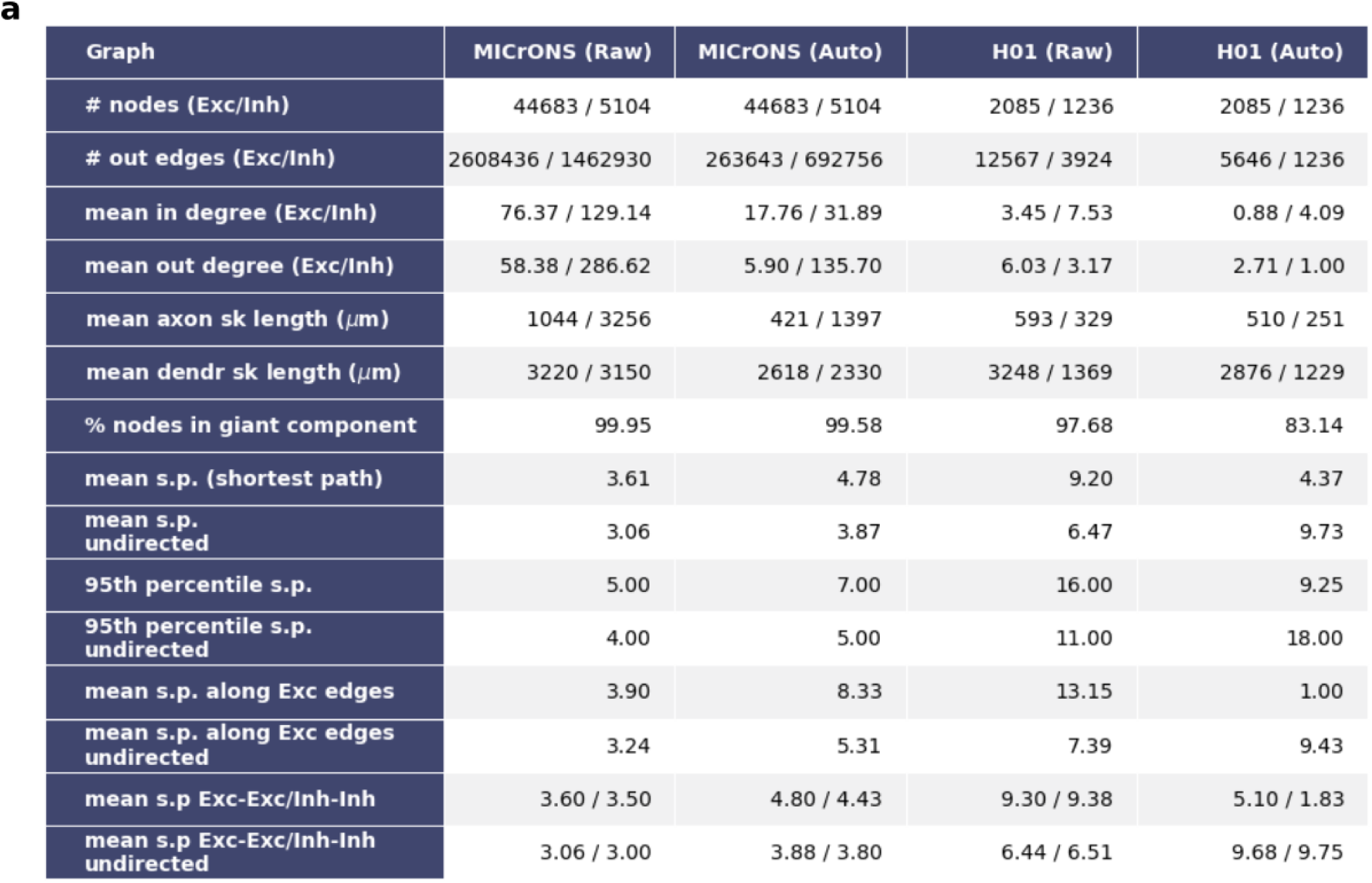
Connectome Network Statistics Table. **a** Network statistics of the MICrONS and H01 connectomes where the nodes are entire single soma neurons and the edges are the synapses between them (neurons with manual proofreading and their associated synapses are excluded). The term “raw” refers to synaptic data before any processing with NEURD, “giant component” refers to the largest connected subgraph, “auto” refers to the connectome produced after the decomposition pipeline and automated proofreading, “sk” abbreviates “skeletal” when referring to skeletal distances, and “sp” abbreviates “shortest path”. Once all neurons were cleaned and annotated from the NEURD decomposition process there were on the order of 10^8^ and 10^7^ individual presynaptic or postsynaptic connections for the MICrONS and H01 dataset respectively. However, in order for a connection to be included in the connectome, both the presynaptic and postsynaptic information must be present in the data, which was not the case for a majority of connections and is why the number of edges in the connectome is much smaller than number of individual synapses available in the cleaned datasets.

**Fig. 28.**
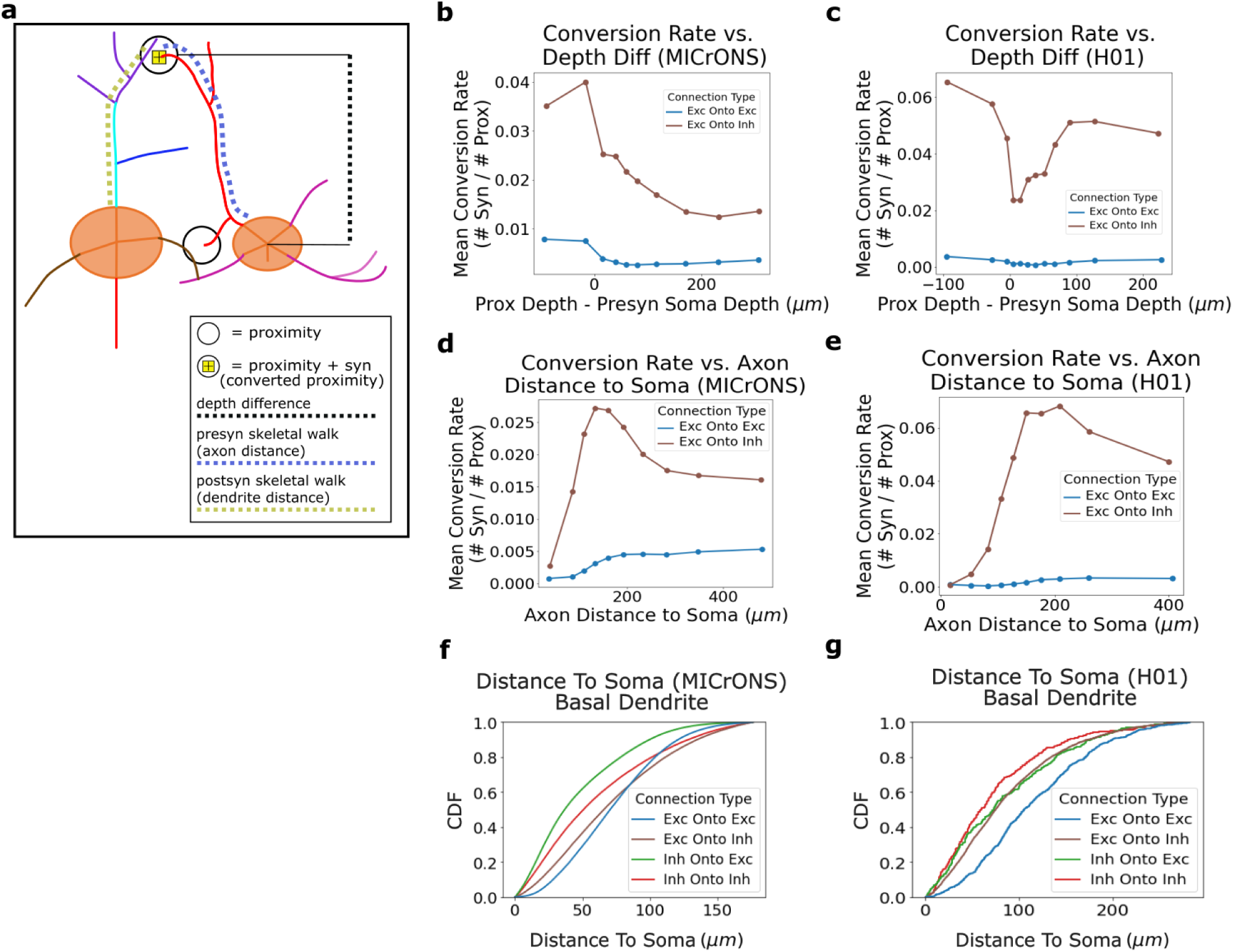
Path distances and relation to conversion conversion rate. Features computed by the NEURD preprocessing pipeline make available path lengths and distances along x/y/z dimension or dendritic/axonal walk between presynaptic or postsynaptic soma centers and their synapse or proximity locations. For calculating conversion rates, proximities are binned (approximate equal depth bins) in terms of their relative depth to the presynaptic soma center (proximity depth - presynaptic soma depth) or axonal path distance and then the mean conversion rate (number of synapses / number of proximities) for that bin is computed for different connection types **a** Illustration showing a pair of cells with both a proximity with a converted synapse and one without from an inhibitory cell to an excitatory cell. Each of the paths or distances used in the later panels is shown with a dotted color path: depth difference (proximity depth - presynaptic soma depth; the example drawn would have a negative depth difference because the presynaptic soma is deeper in the volume than the proximity), presynaptic skeletal walk (axonal skeletal distance from the presynaptic soma center to the synapse or proximity), postsynaptic skeletal walk (dendritic skeletal distance from the postsynaptic soma center to the synapse or proximity). **b,c** Conversion rate as a function of relative proximity depth. In the MICrONS volume, the plot demonstrates that the conversion rate for excitatory connections onto both excitatory and inhibitory postsynaptic cells is greater when the proximity is above the soma (for both connection types); in the H01 volume, the plot demonstrates a greater conversion rate above the soma than below, but with an additional reduction in conversion rates close to the soma that is not seen in MICrONS. **d,e** Mean conversion rate as a function of distance from the synapse to the presynaptic cell along the axon. In both the MICrONS and H01 dataset the conversion rate peaks farther away from the soma and then gradually decreases when moving farther downstream. **f,g** Cumulative density function (CDF) of the postsynaptic skeletal walk distance distribution for different exc/inh connection combinations (apical and soma synapses are excluded). In both datasets, excitatory inputs are further along the dendrite from the soma.

**Fig. 29.**
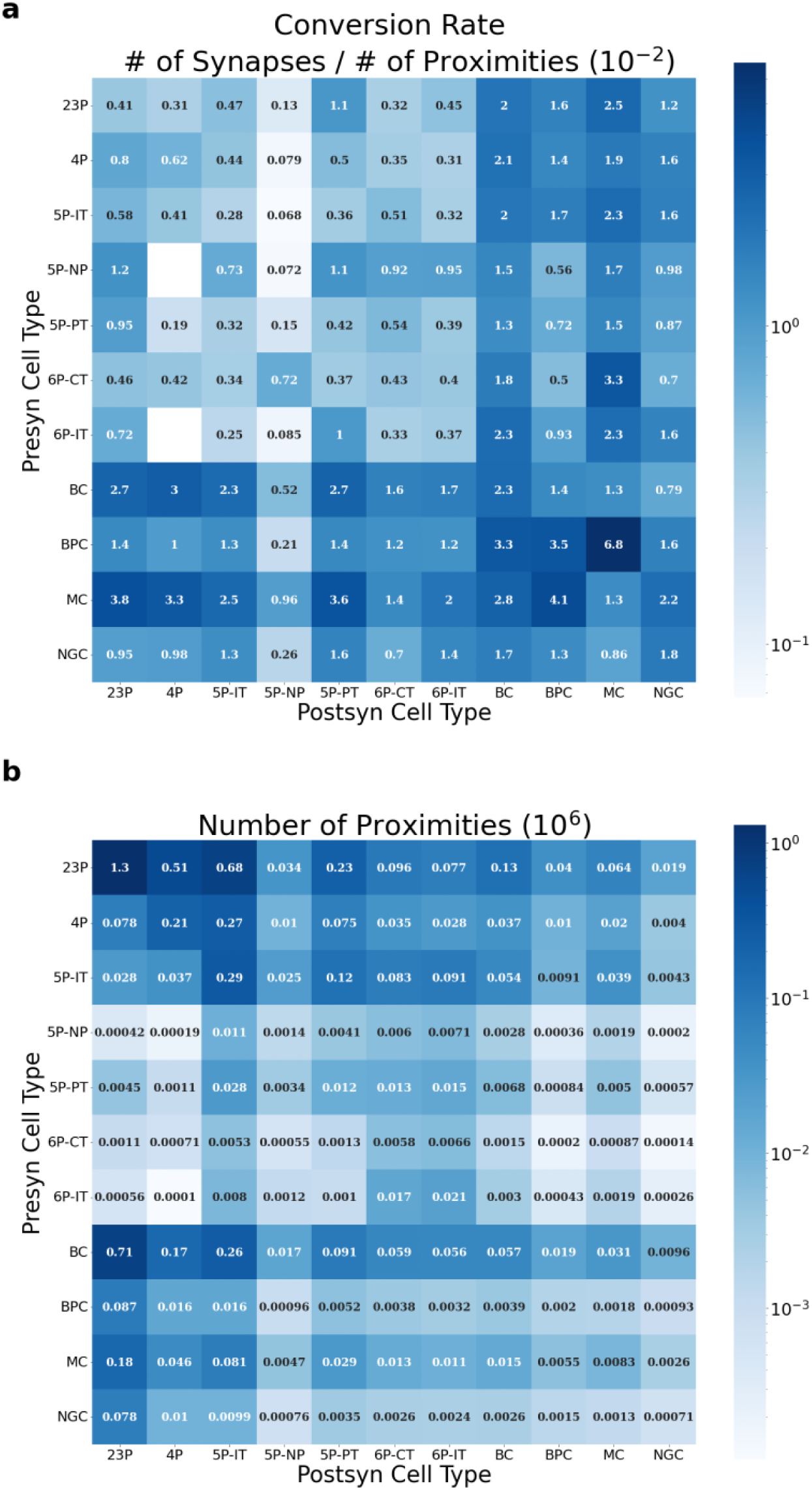
Conversion Rate Cell Type Matrix. The conversion rates (number of synapses / number of proximities) for different presynaptic and postsynaptic cell type pairs. The cell type labels are determined by the GNN whole neuron classifier. Proximities are filtered to only include those with the following features: less than 3 *µ*m proximity distance, dendrite only postsynaptic compartment, presynaptic proximity width less than 130 nm (to exclude myelinated axon), presynaptic and postsynaptic cell type labels with at least a 70% confidence for each from the GNN classifier. **a** Conversion rate for different cell type presynaptic and postsynaptic combinations **b** Number of proximities in dataset used to calculate the conversion rate.

**Fig. 30.**
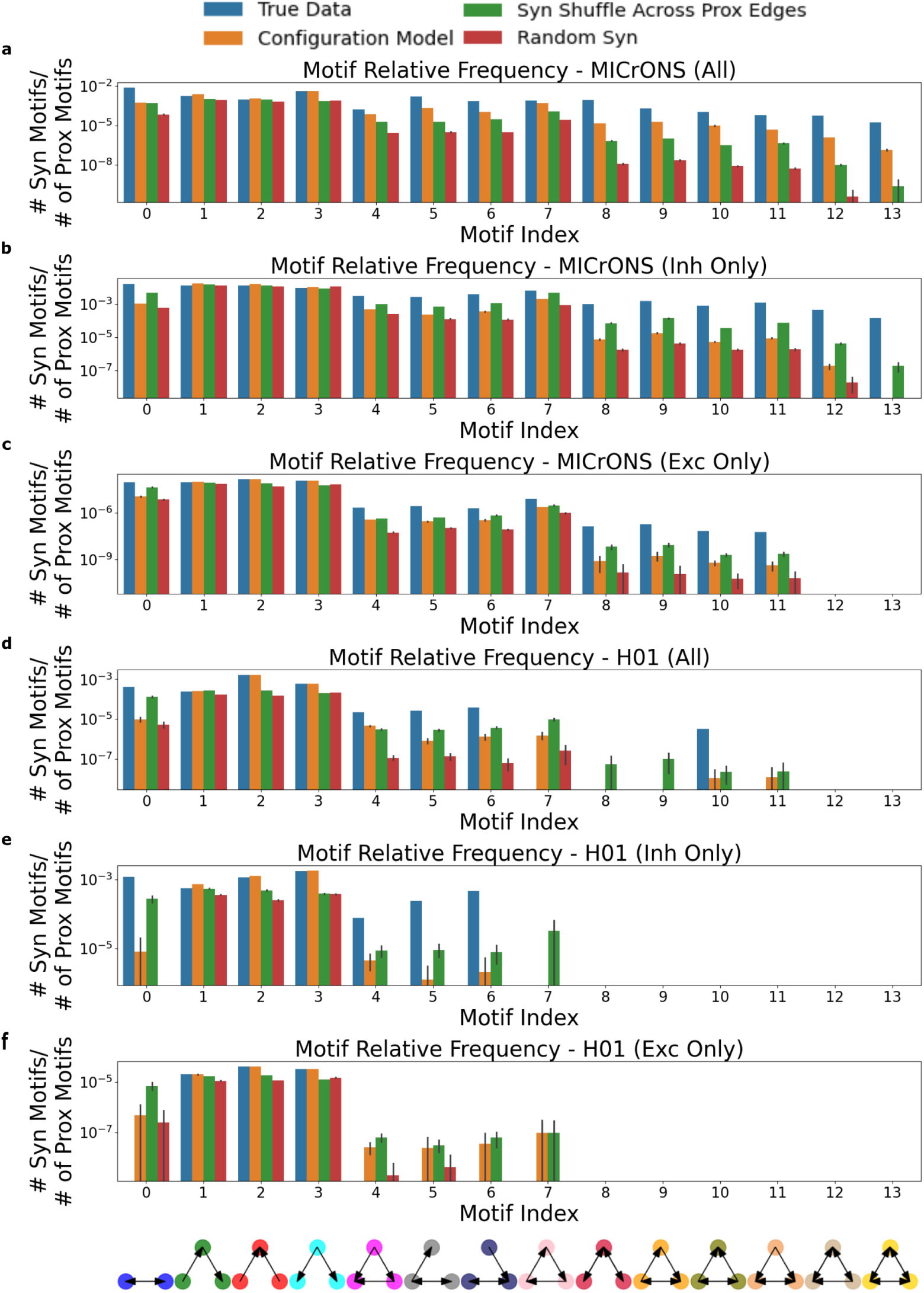
Higher-order Triangle Motif Analysis. Across all automatically-proofread neurons, we count the number of reciprocal connections and directed triangle motifs in the synaptic and proximity connectome and compare the observed ratios to null ratios from three different models: first, a model where synaptic degree distribution is held the same but edges are shuffled (configuration model), second, a model where the synaptic edges are shuffled only between neurons with an existing proximity edge, or third, a model where synapses are randomly shuffled between neurons regardless of proximity. In addition to testing on the entire connectome (“All”), we also performed the same set of comparisons on the inhibitory (“Inh Only”) and excitatory (“Exc Only”) subgraphs to test whether higher-order motif frequencies were consistent in these subpopulations. **a-c** MICrONS dataset relative frequencies (duplicated from Fig. 5g) showing that the relative frequency of higher-order motifs in the synaptic connectome decreases as the number of edges in the motif increases (more higher-order), but are consistently higher than the null model controls (250 random graph samples for each null distribution comparison) for all subgraphs. **d-f** H01 dataset relative frequencies showing that the relative frequency of higher motifs in the synaptic connectome decreases as the number of edges in the motif increases. The observed motif frequencies are again higher than the null models (except in some more edge-dense 3 node motifs in the inhibitory and excitatory only subgraphs), but many of the motifs with more than three directed edges are not observed due to the more incomplete reconstruction of neurons in H01 (400 random graph samples for each null distribution comparison, more samples were computed than MICrONS because computation was faster with a smaller connectome).

**Fig. 31.**
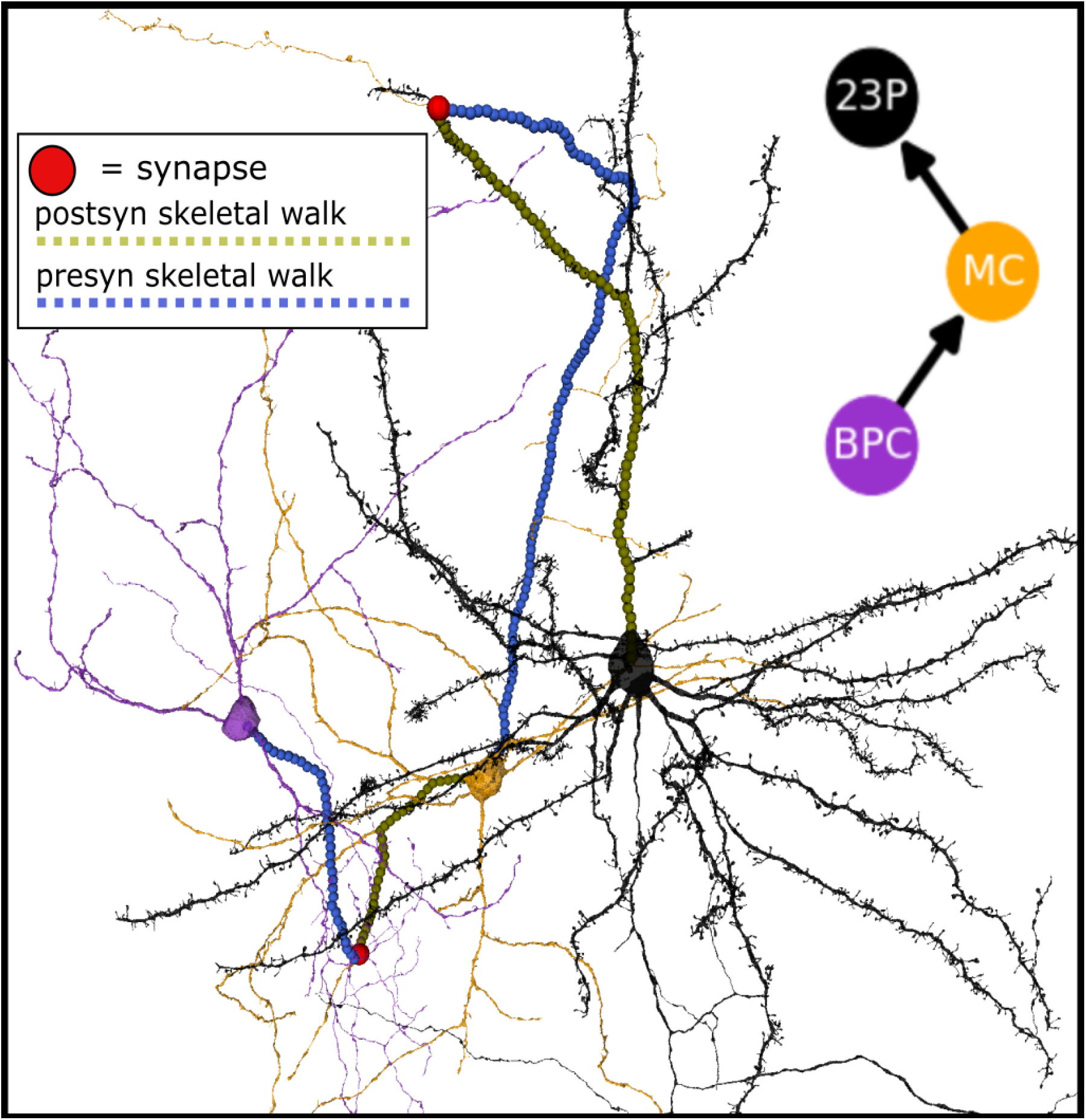
Higher-Order Cell Type Motif Search. Cell-type specific connections and motifs in the MICrONS dataset can be found by querying the annotated connectivity graph. NEURD allows for visualization of these connection paths and motifs so they can be quickly inspected.

**Fig. 32.**
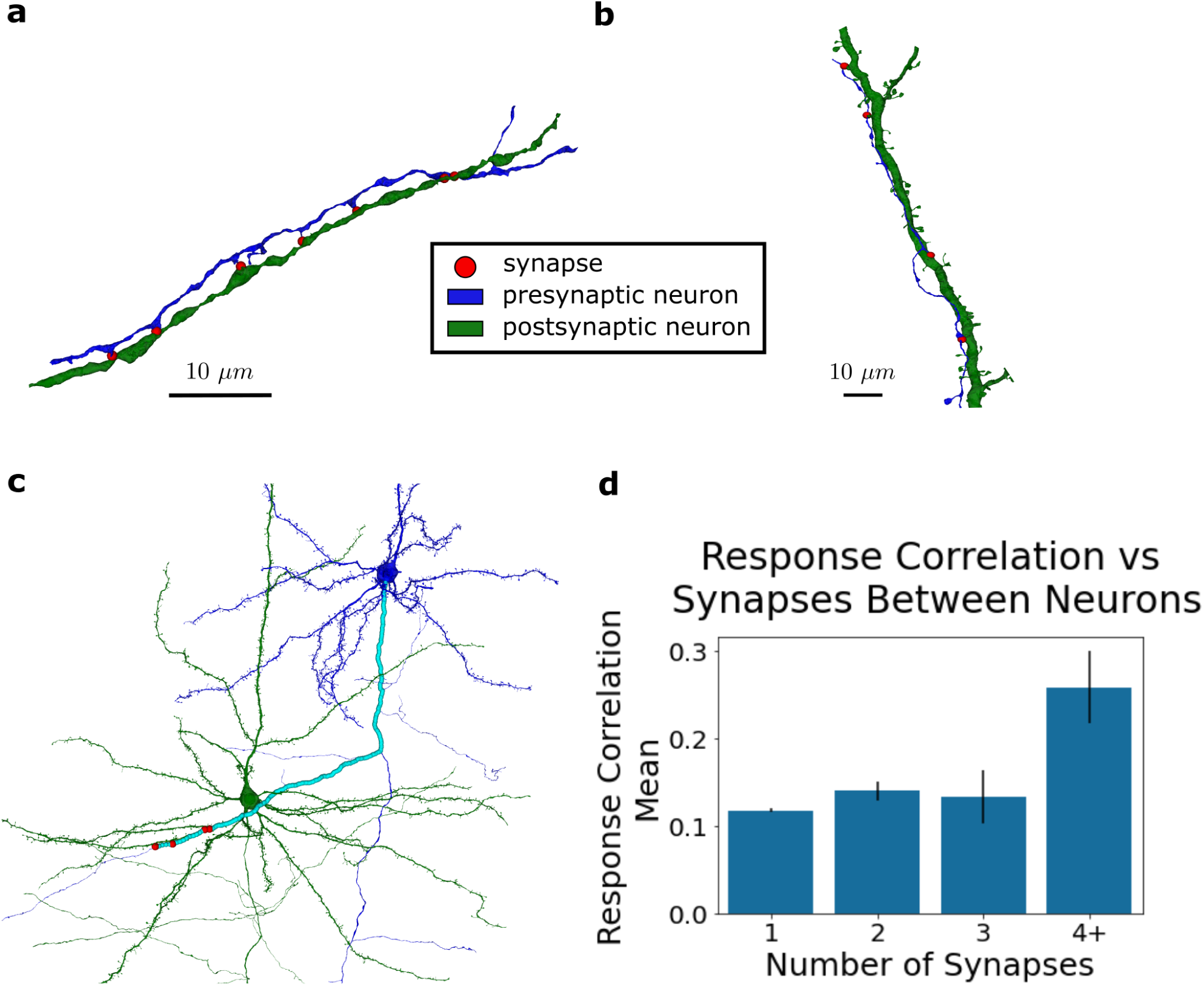
Functional Connectomics Illustration: High-degree cell pairs. **a** Example multi-synaptic connection (n=7 synapses) from an excitatory to inhibitory neuron in the H01 dataset **b** Example multi-synaptic connection (n=4 synapses) from excitatory to excitatory neuron in the MICrONS dataset. **c** Example of a highly spatially clustered multisynaptic connection (n=4 synapses) on a postsynaptic basal dendrite from a neuron cleaned with automated proofreading in the MICrONS dataset (presynaptic skeletal walk shown in aqua, synapses shown in red) **d** Distribution of response correlation between pairs of functionally matched excitatory neurons in the MICrONS dataset. Response correlation is significantly larger for pairs of neurons with 4 or more synapses connecting them (n=11 pairs) compared to those with 1, 2, or 3 synapses (n=5350, 280, 34 pairs respectively).

**Fig. 33.**
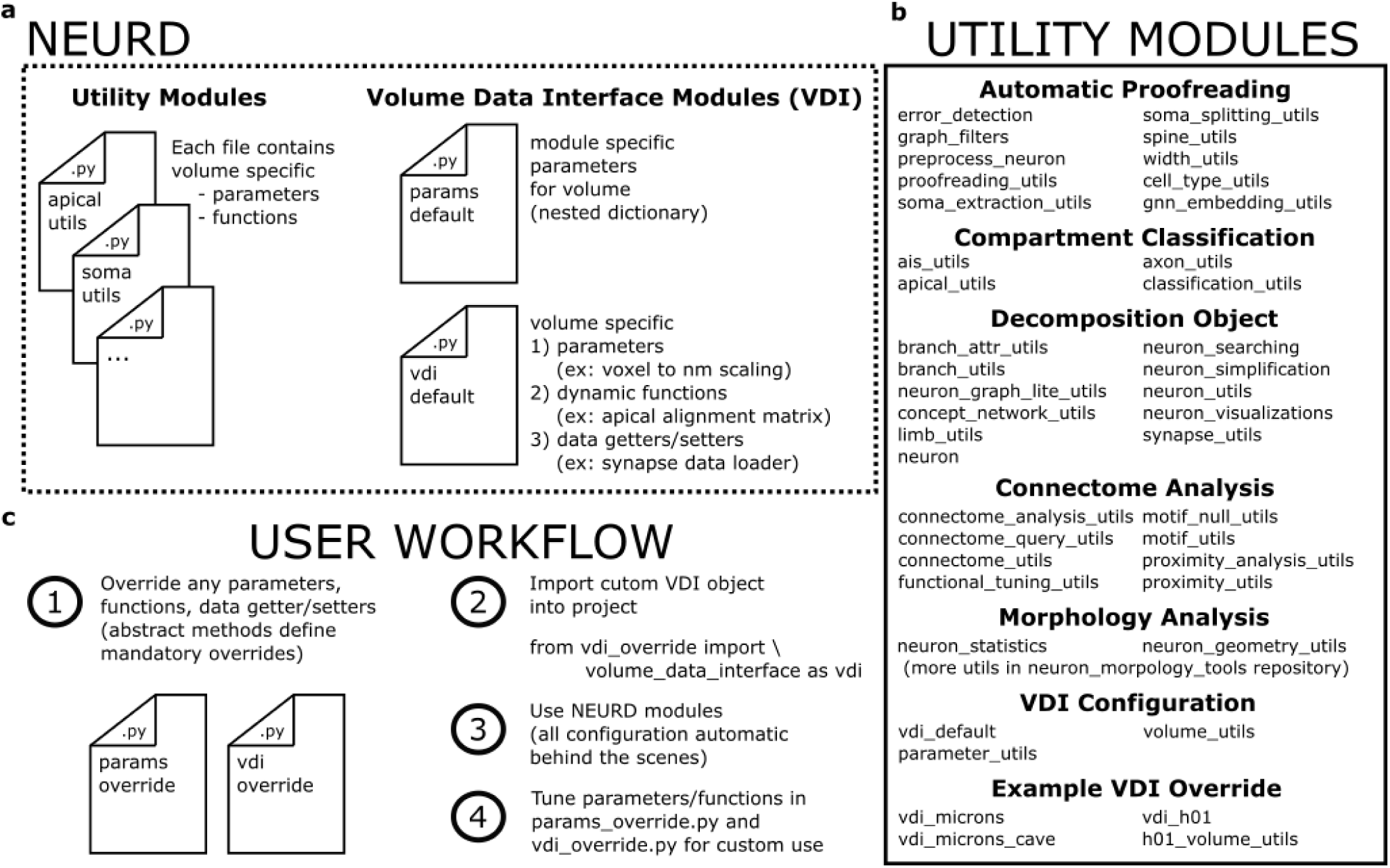
NEURD Code Package Overview. **a** The general structure of the code package modules. Most of the modules (.py files) are utility functions that define the functionality pertaining to a specific processing step (ex: *error_dection.py* ) or neuron feature (ex: *spine_utils.py* ). There are then two .py files that each define a “Volume Data Interface” (VDI). Together they set the user’s settings/parameters for NEURD and facilitate data setting and fetching. The VDI modules have default implementations, but NEURD users can override these files for custom implementations (examples of how to override these VDI files are provided in the code documentation). **b** List of utility modules available in NEURD at the time of publication **c** Step by step procedure for how a user would override/configure, import and use the NEURD package

**Fig. 34.**
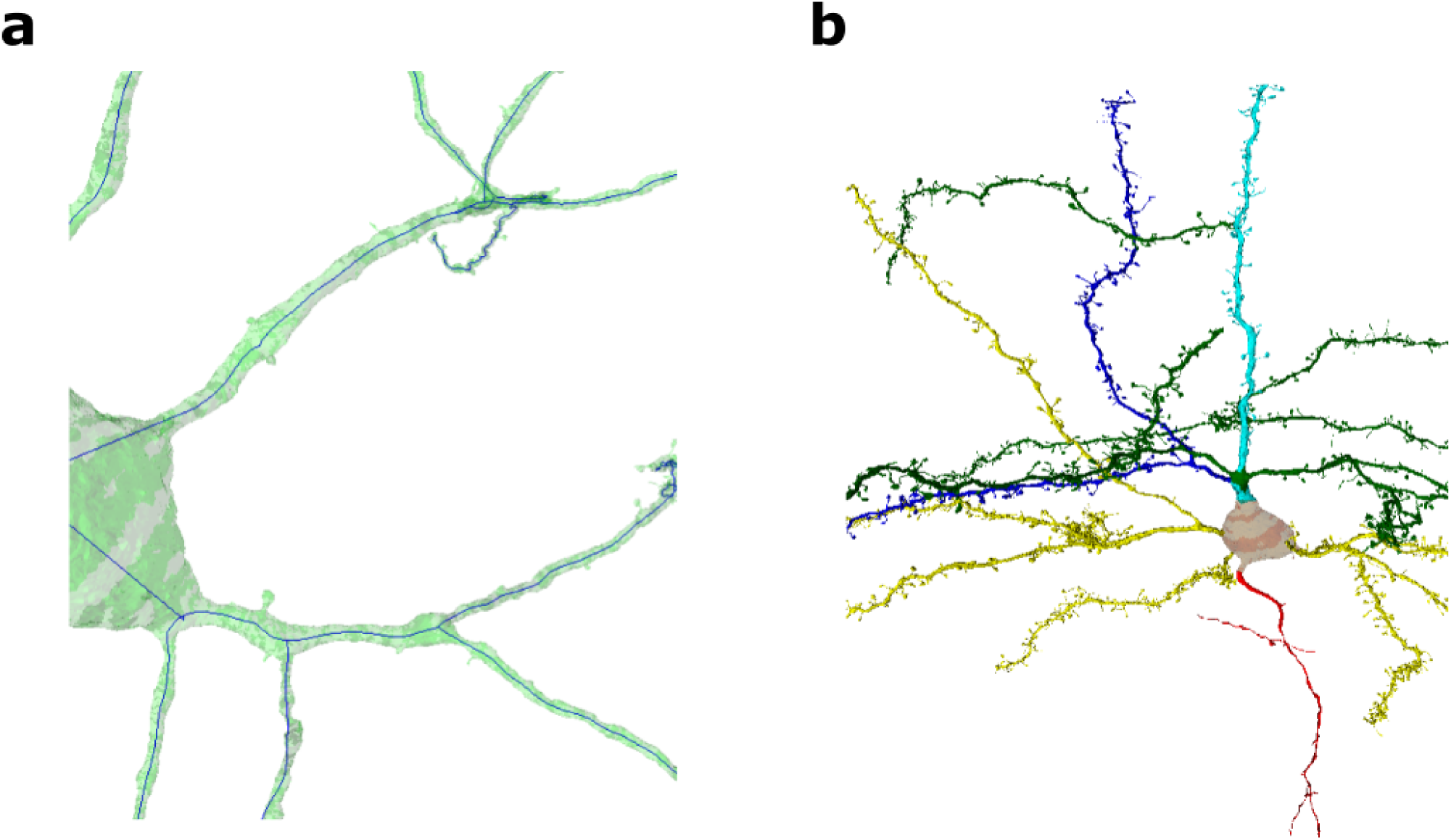
Skeleton and Compartment Visualization. **a** A general example of a skeleton generated by the NEURD package where most thick dendritic segments are represented by a skeleton that tracks the inside middle of the segment, and thinner segments are represented with surface skeletons. The inhibitory neuron in this example is available for 3-D view and closer inspection using the NEURD tutorials on the GitHub repository. **b** A general example of compartment labeling of a neuron’s mesh and skeleton. The excitatory neuron in this example is available for 3-D view and closer inspection using the NEURD tutorials on the NEURD GitHub repository.

**Fig. 35.**
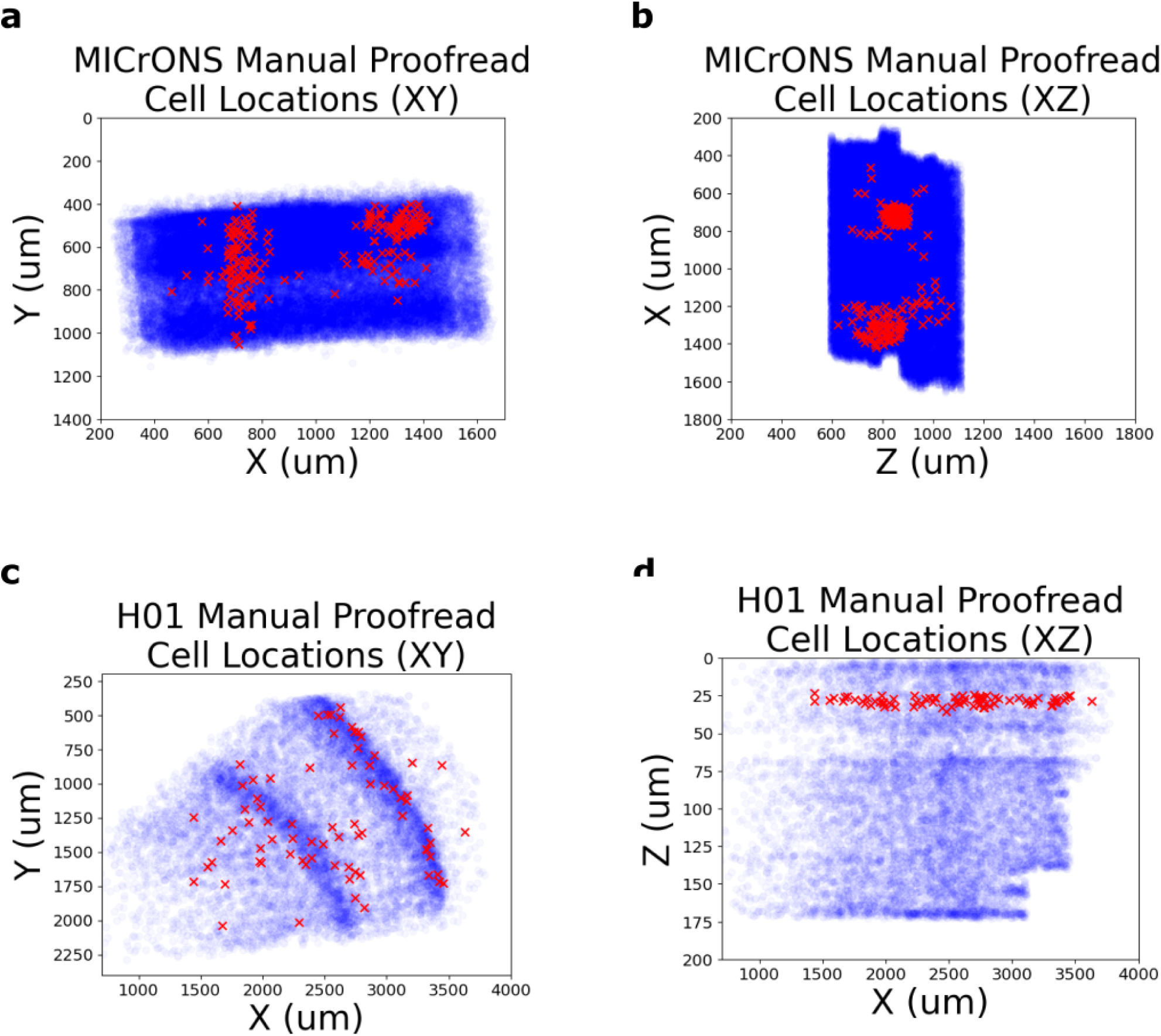
Manual Proofread Cell Locations. **a,b** Locations in the volume (indicated by red “x” markers) of all manually proofread cells used in the MICrONS validation test set. All cells in the volume are plotted in the background in blue. **c,d** Locations in the volume (indicated by red “x” markers) of all manually proofread cells used in the H01 validation test set. All cells in the volume are plotted in the background in blue.

**Appendix Table 1:**
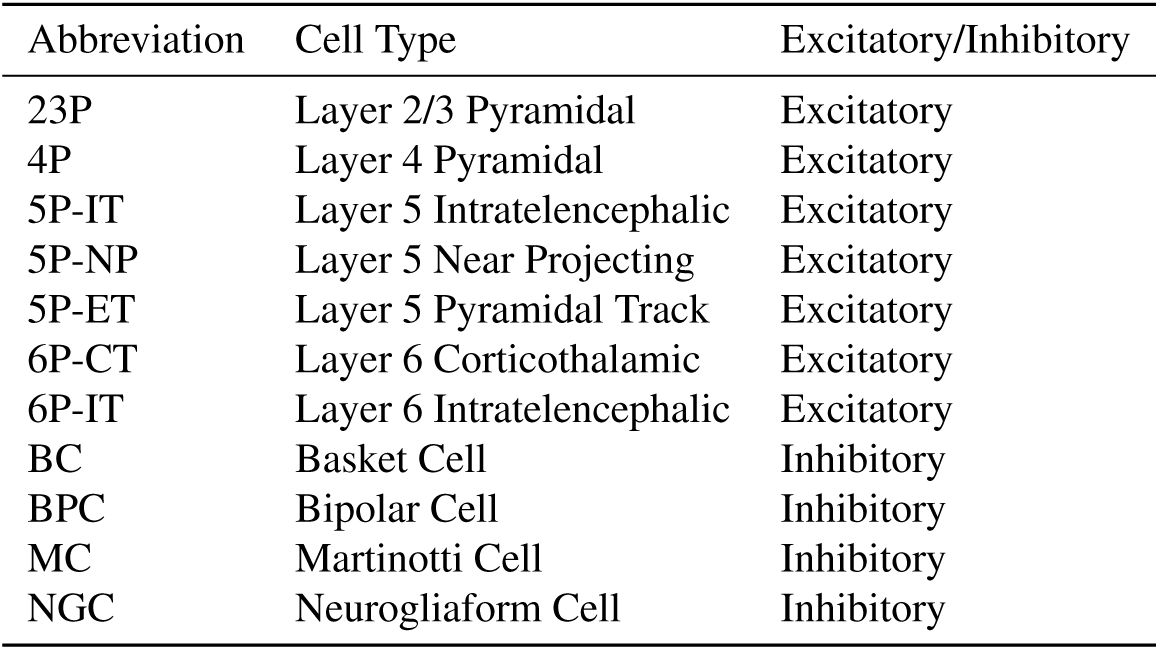
Cell Type Subclass Glossary.

**Appendix Table 2:**
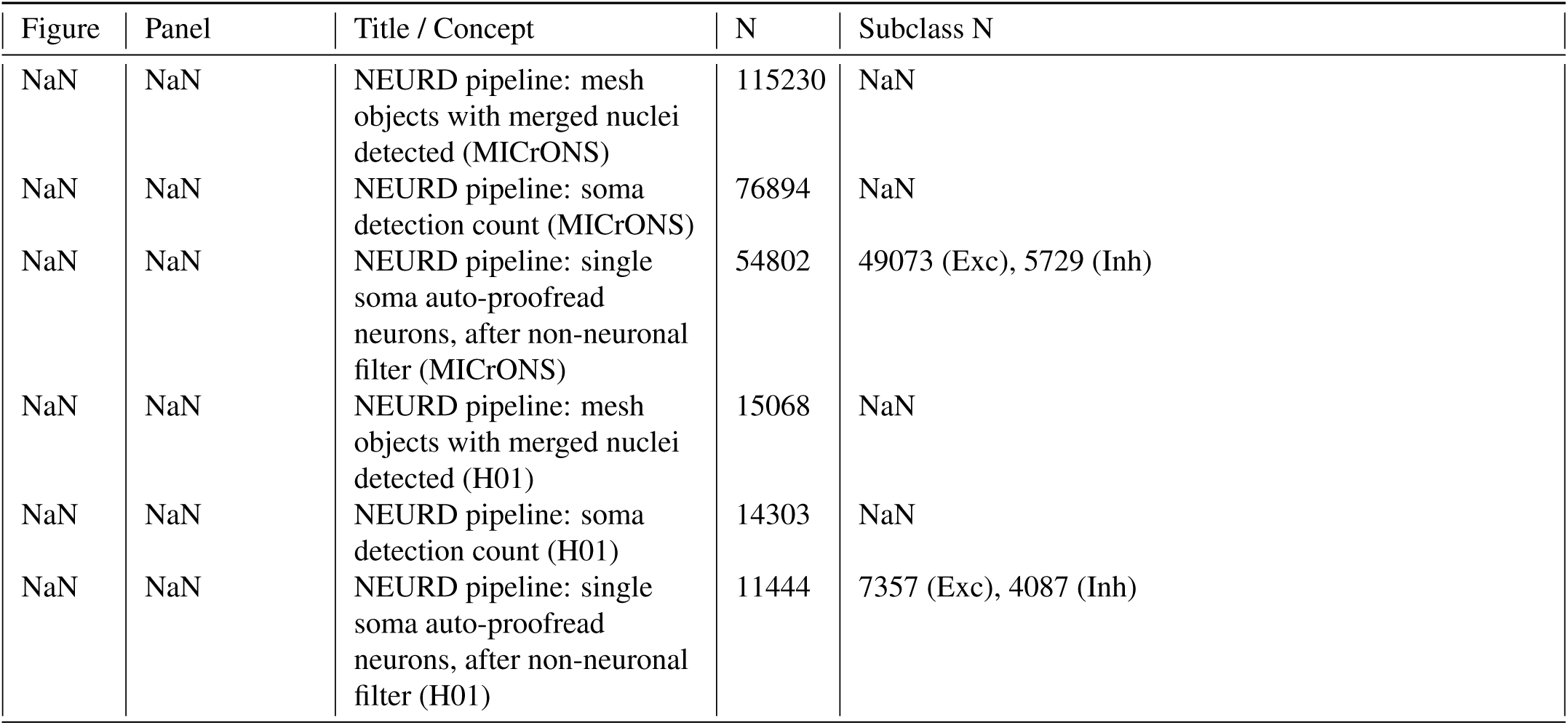
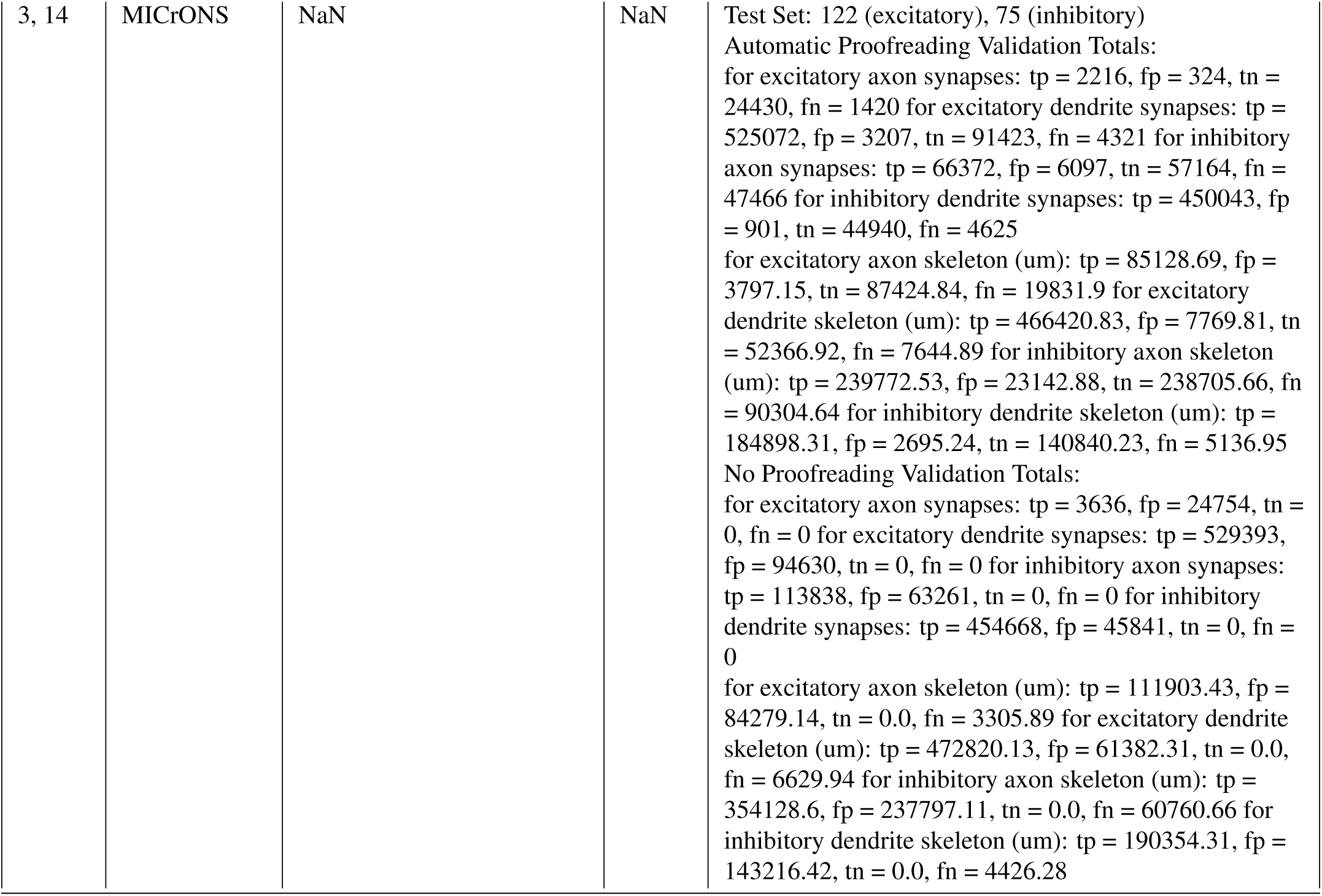
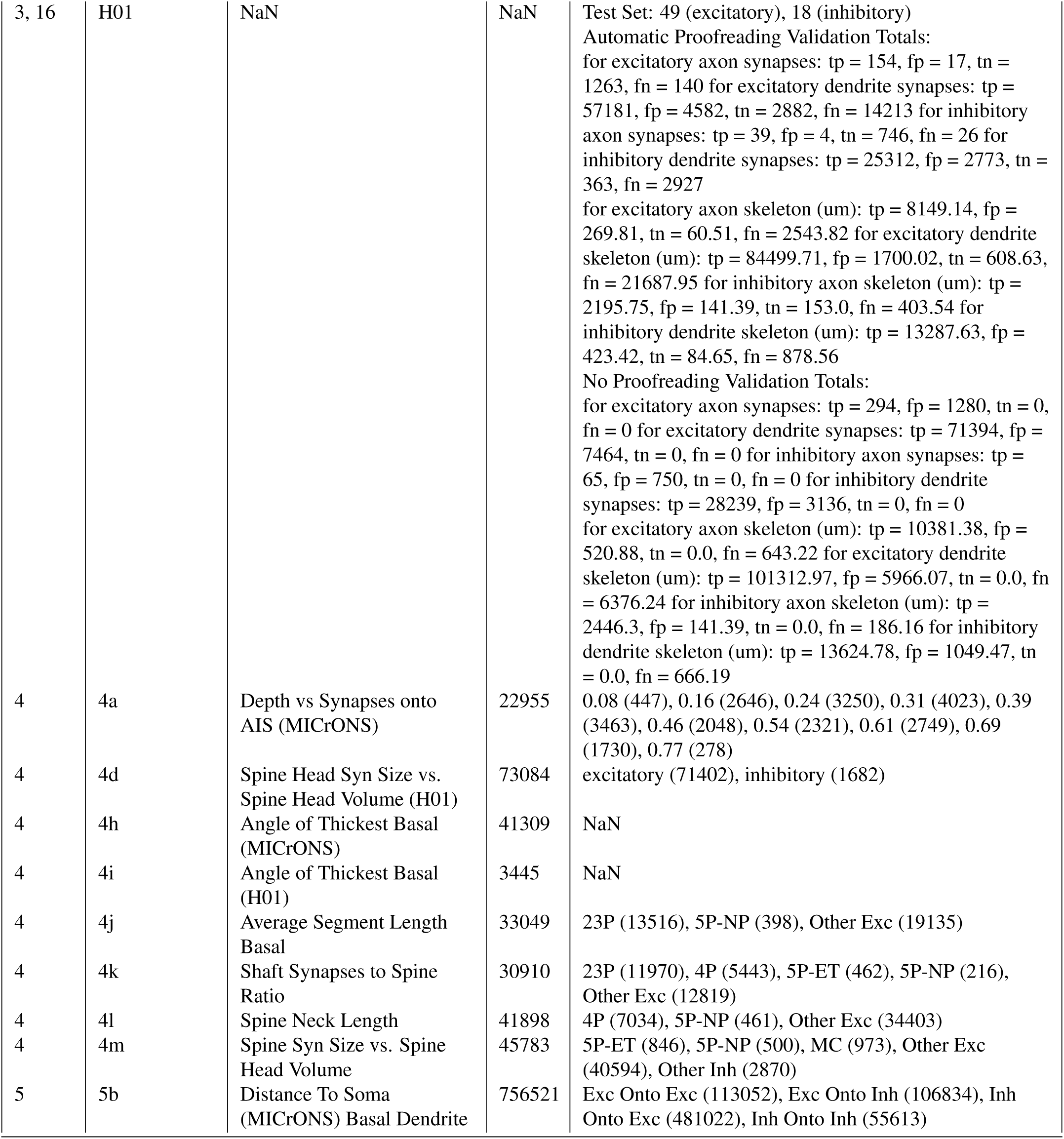
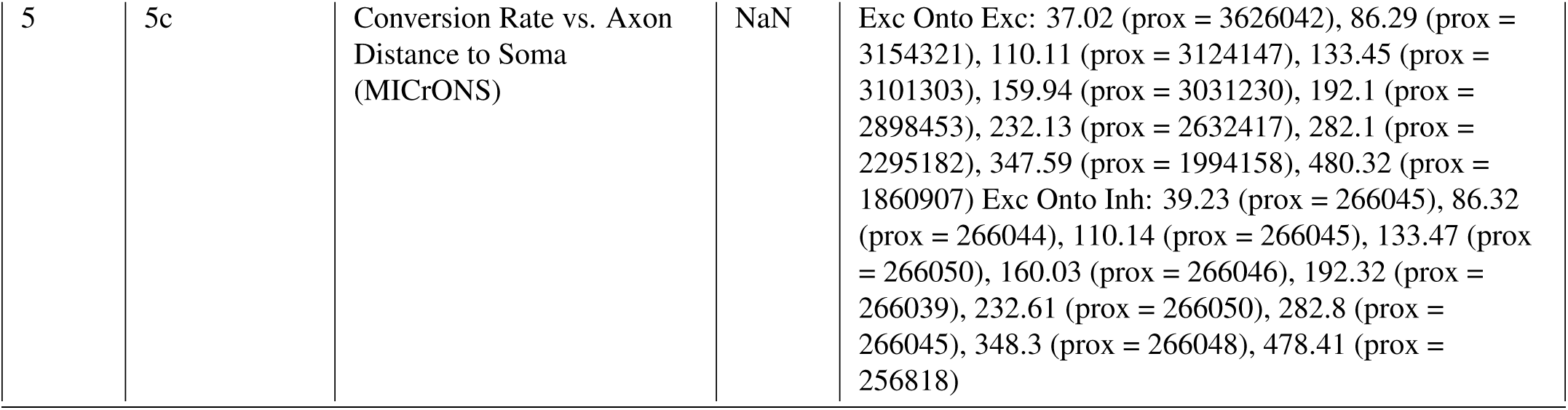
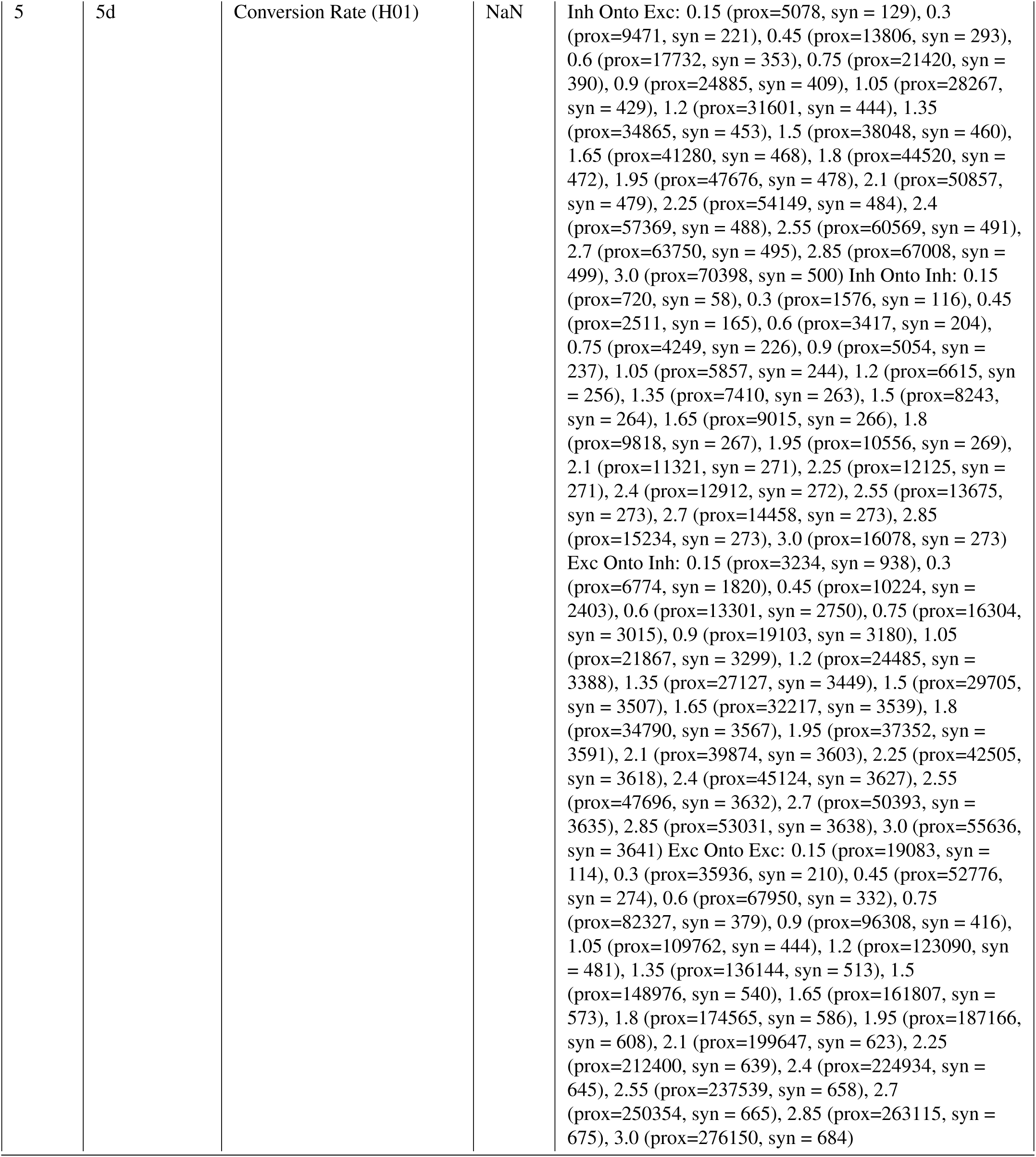
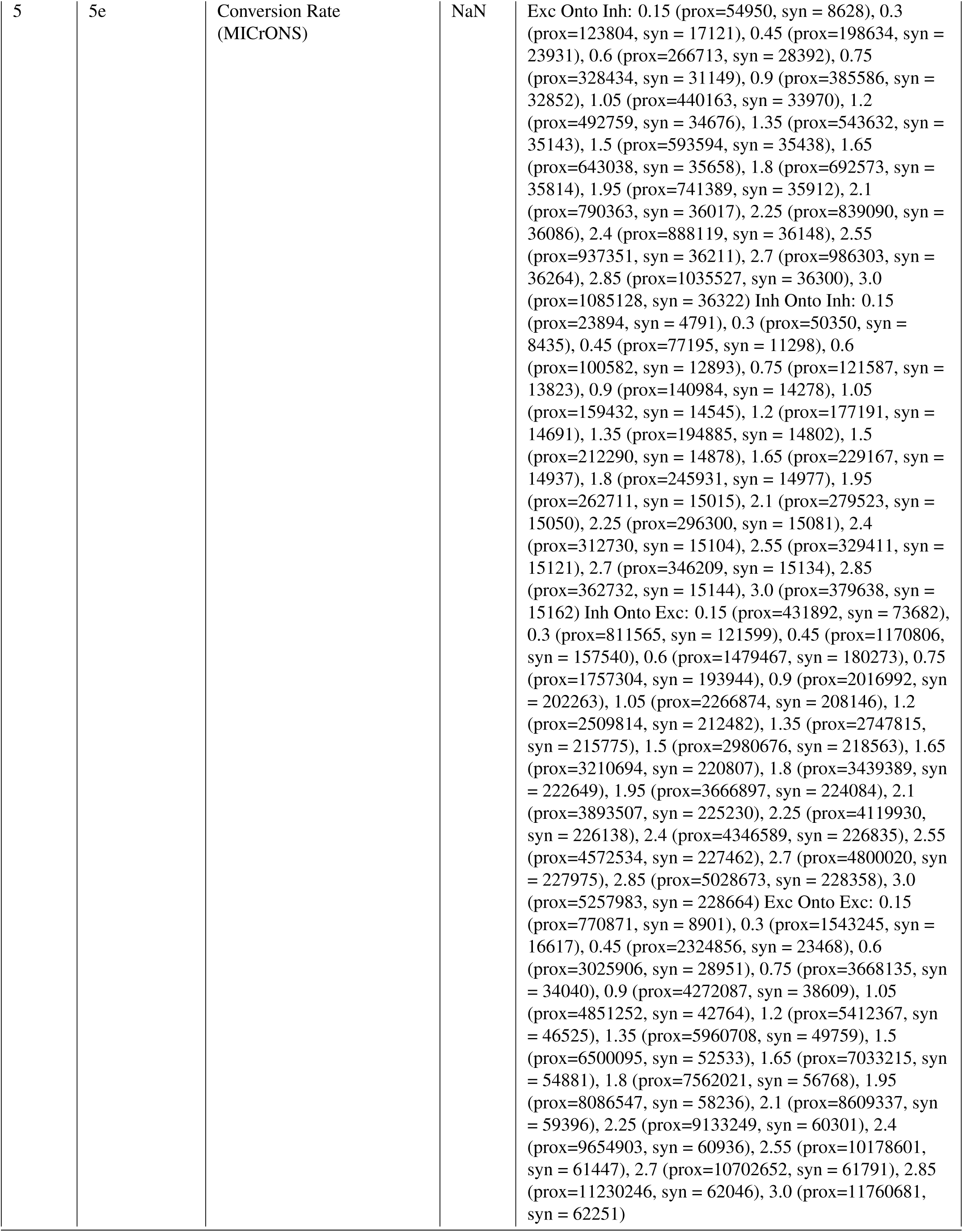
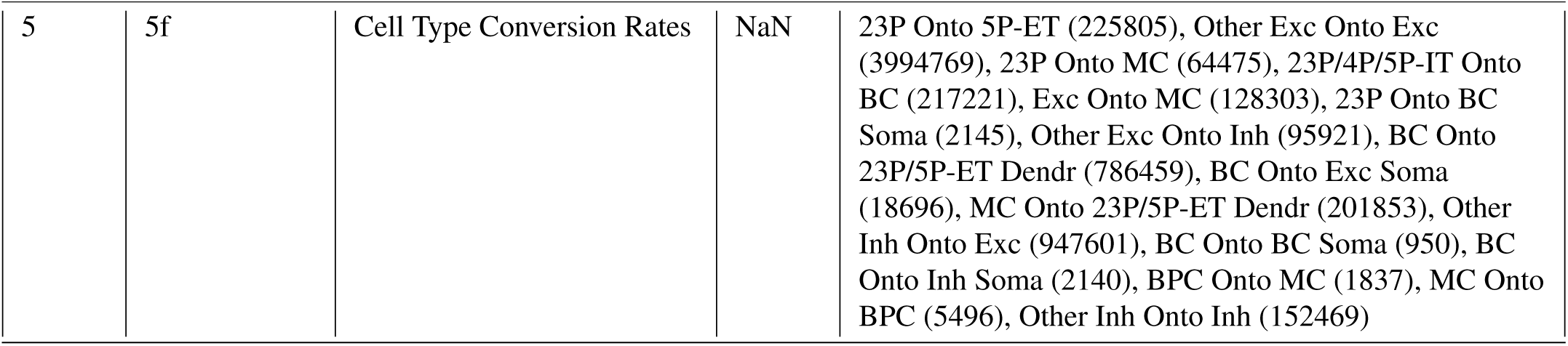
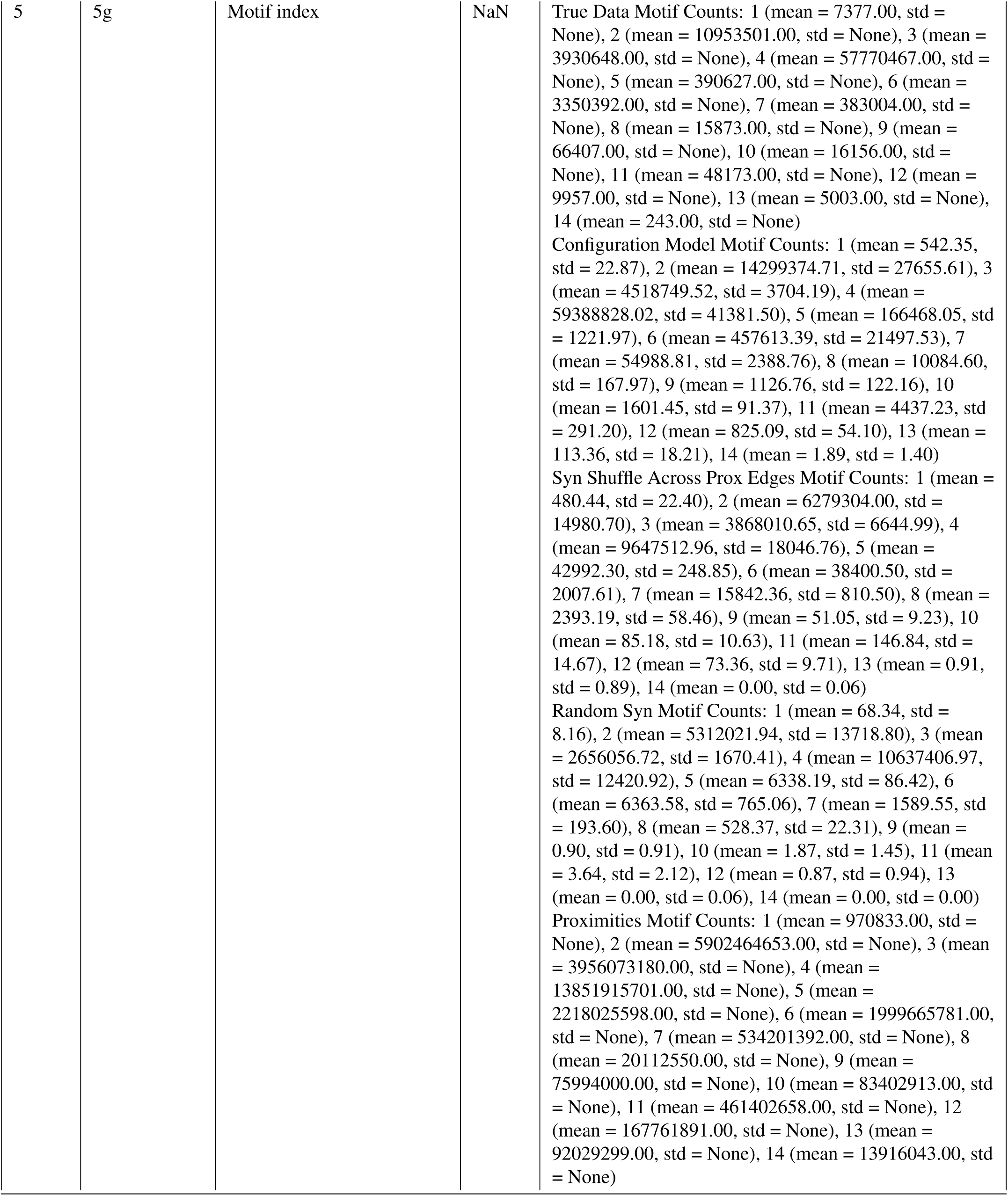
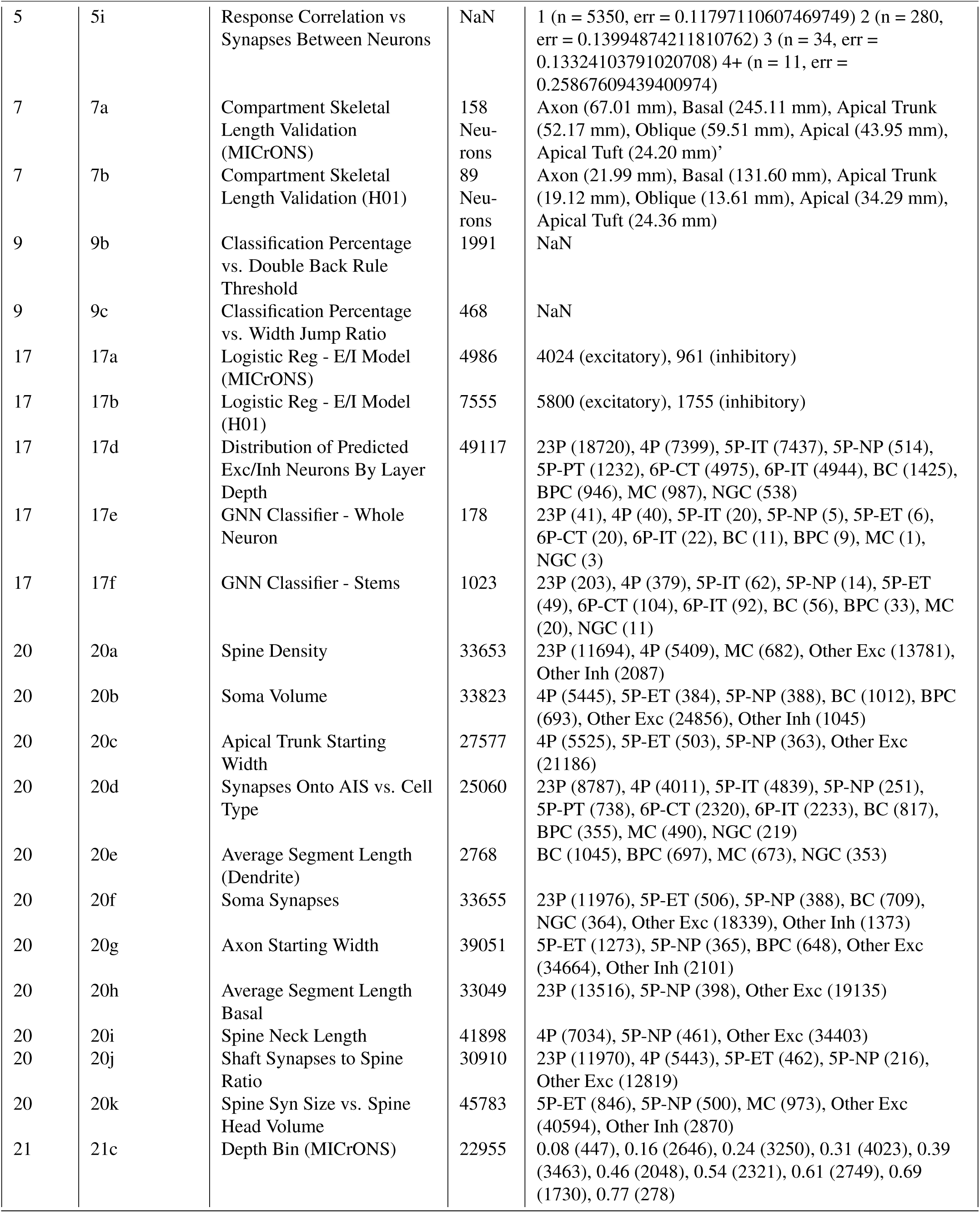
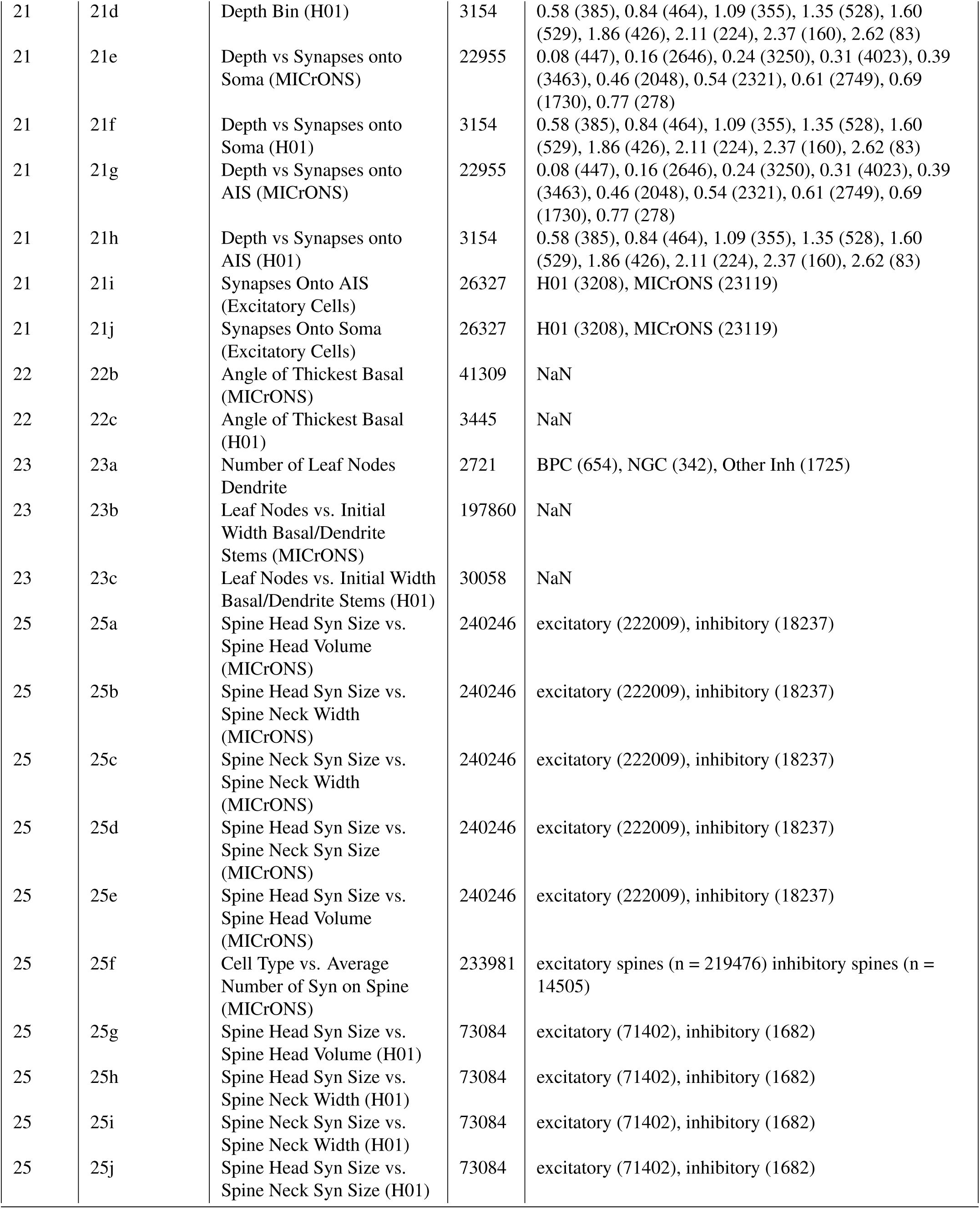
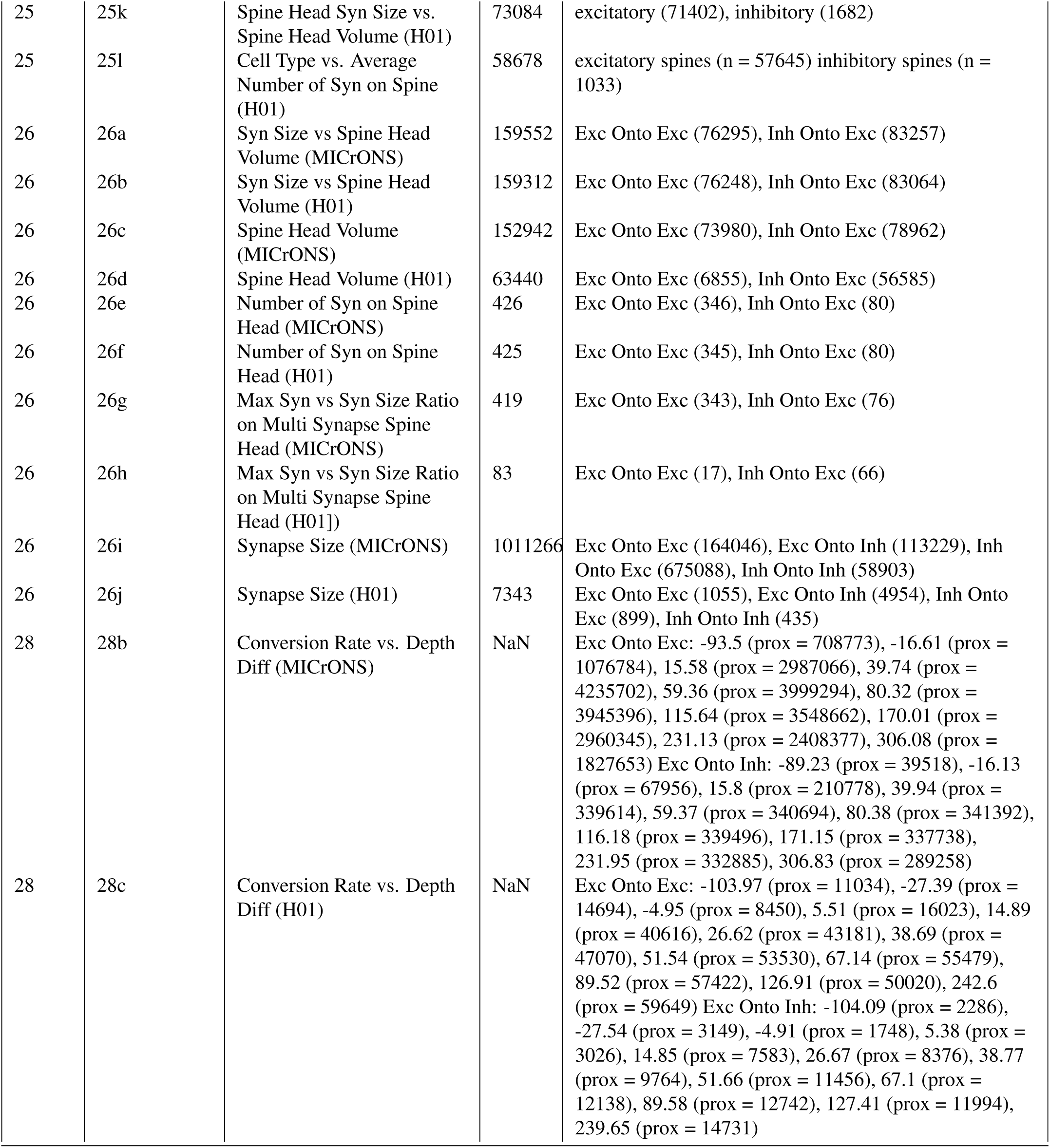
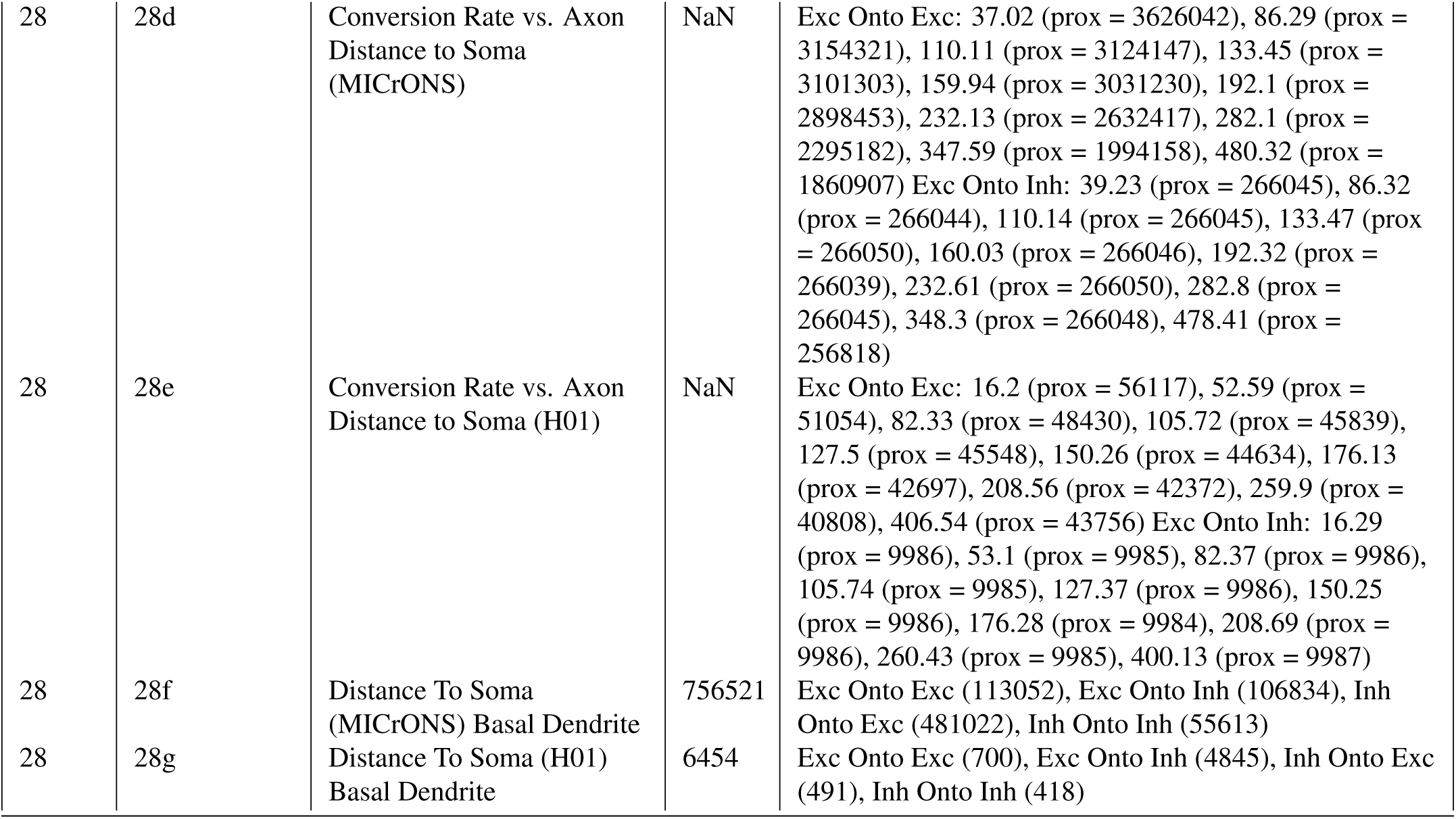
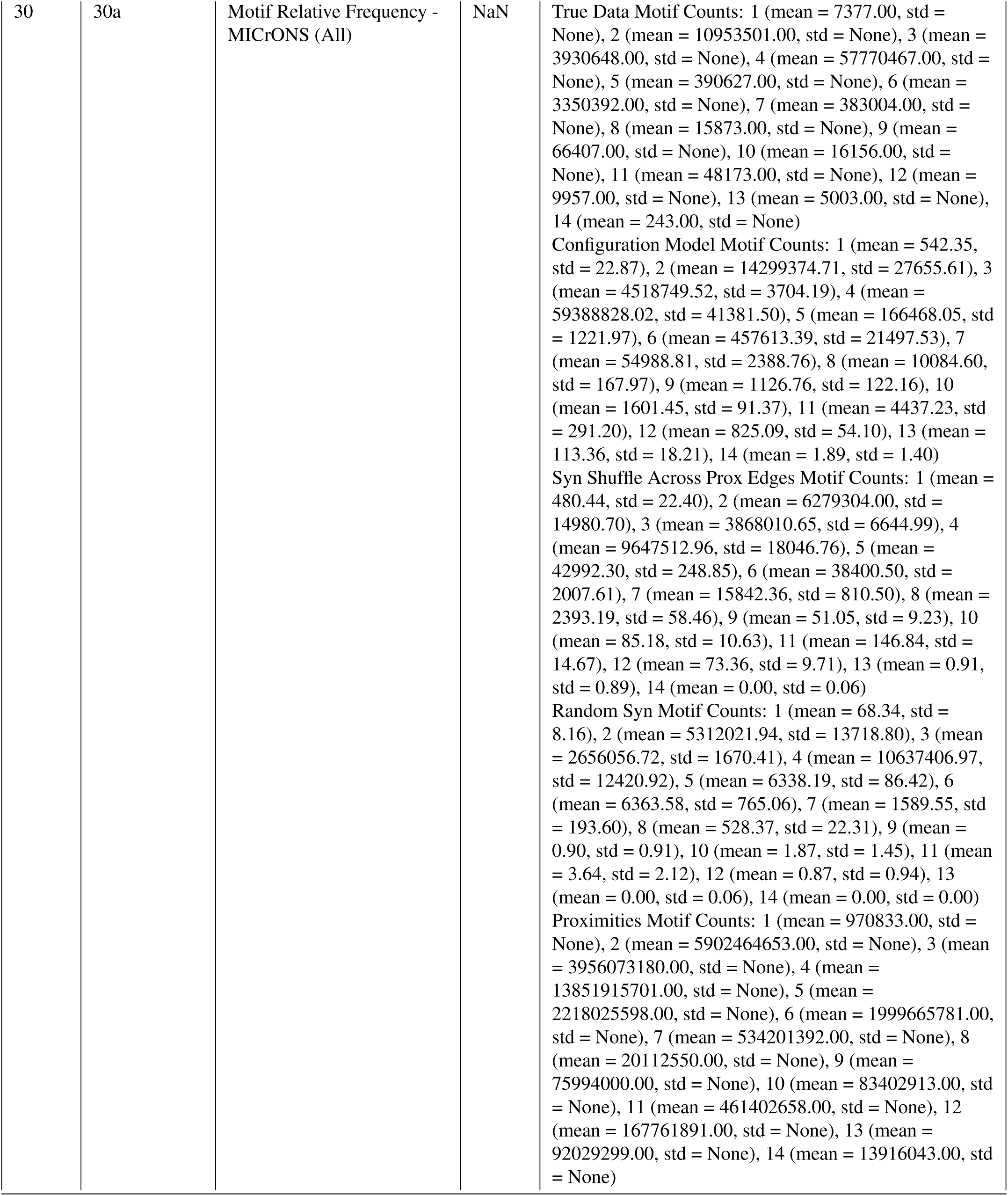
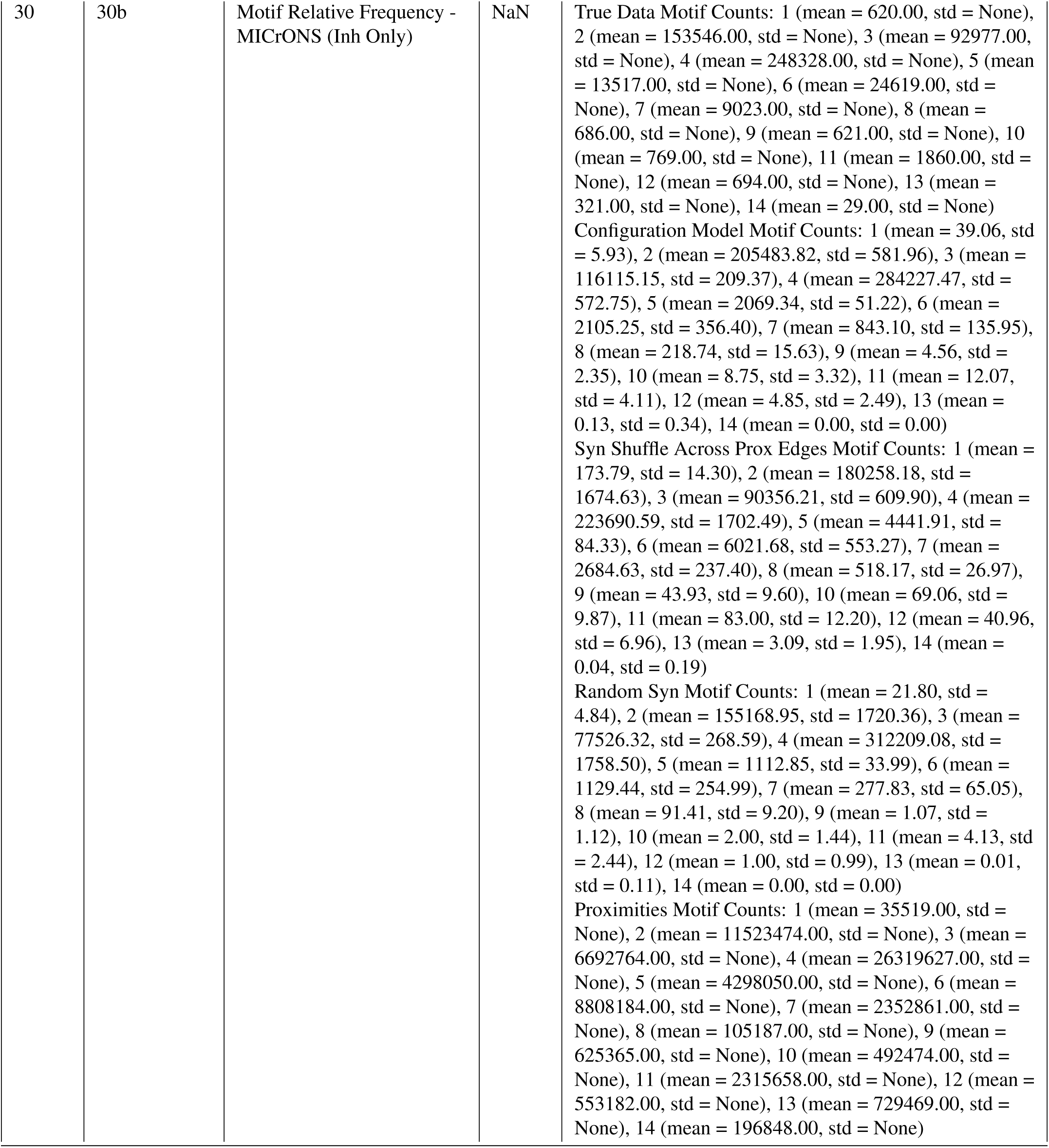
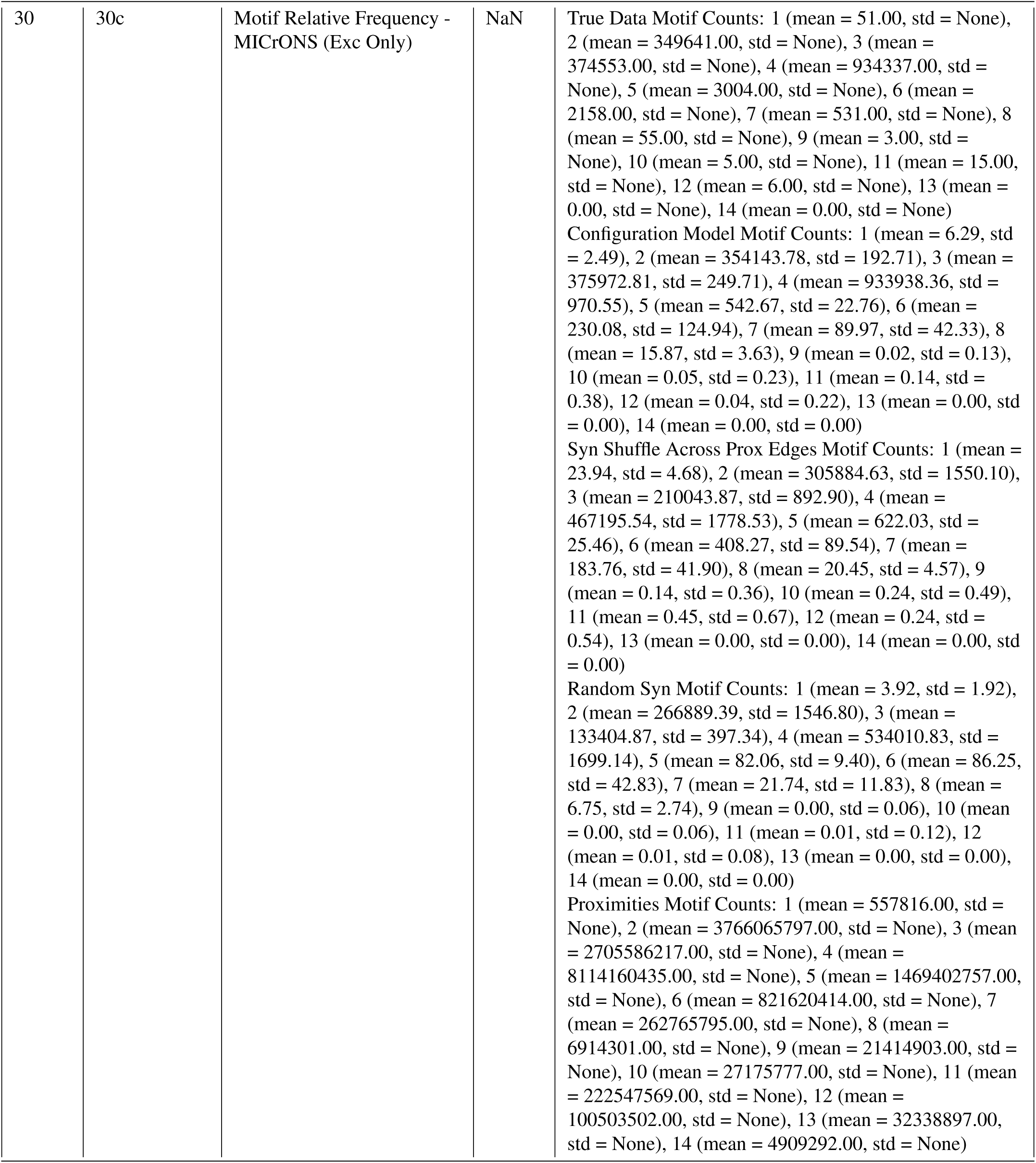
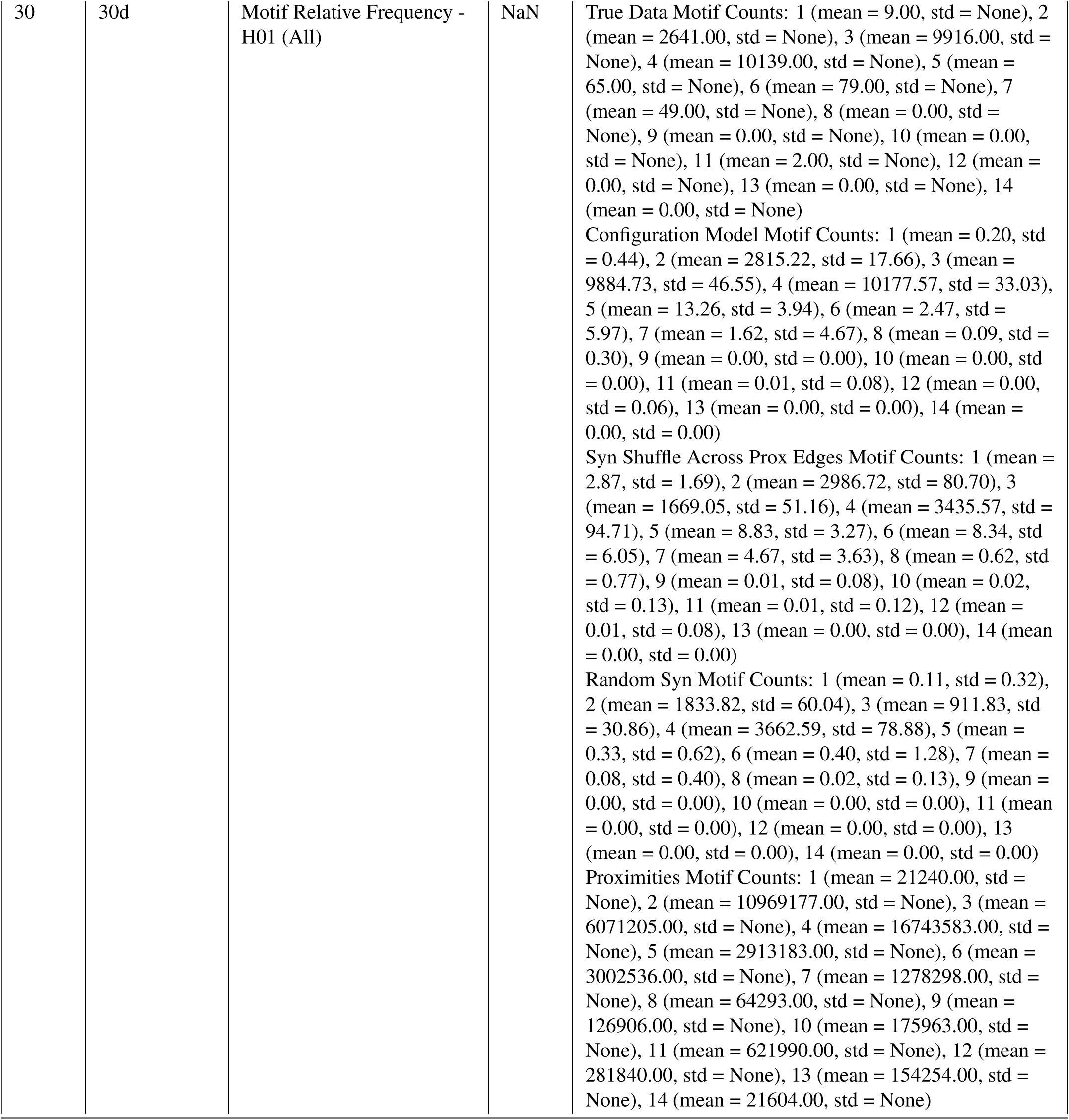
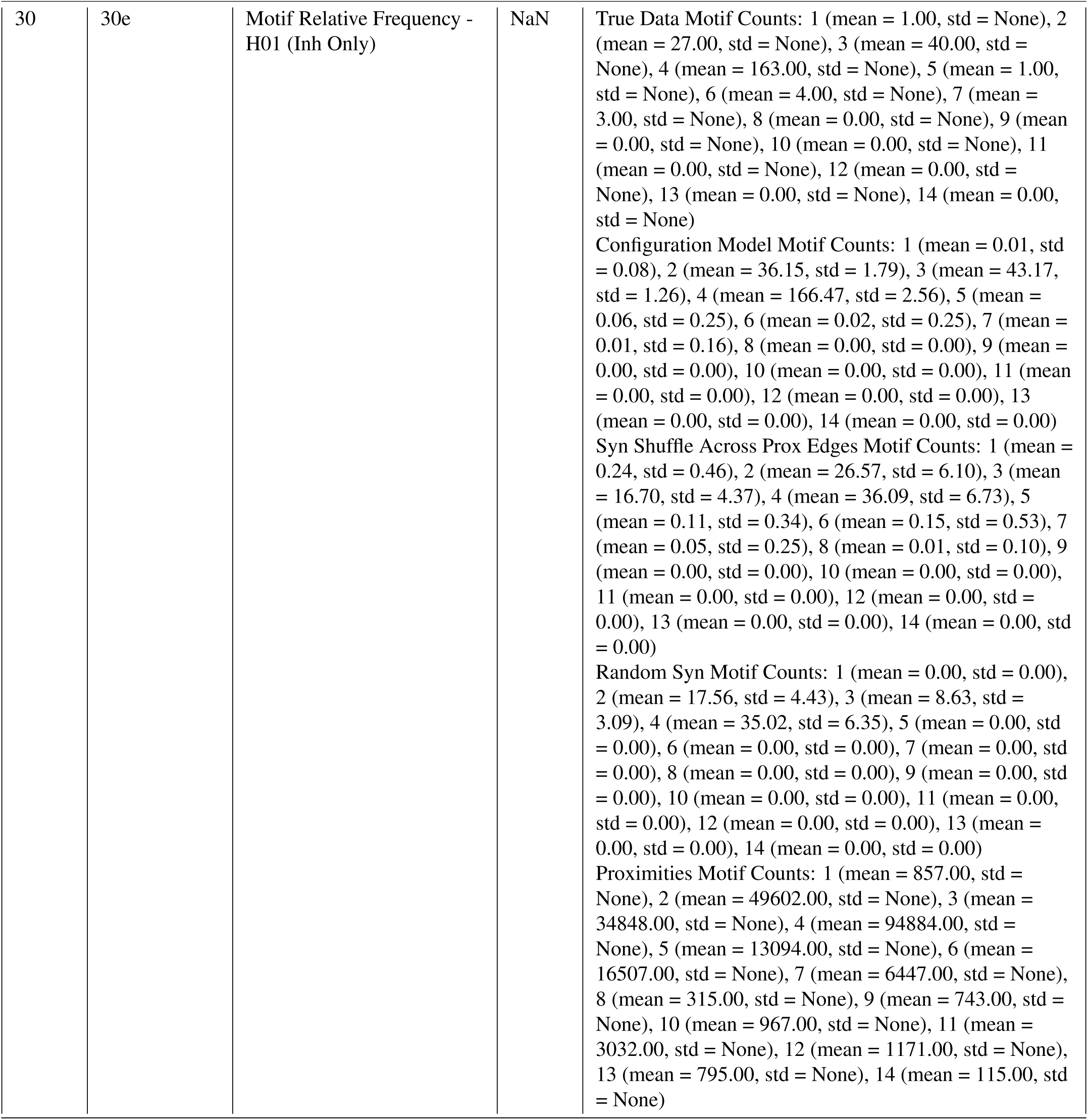
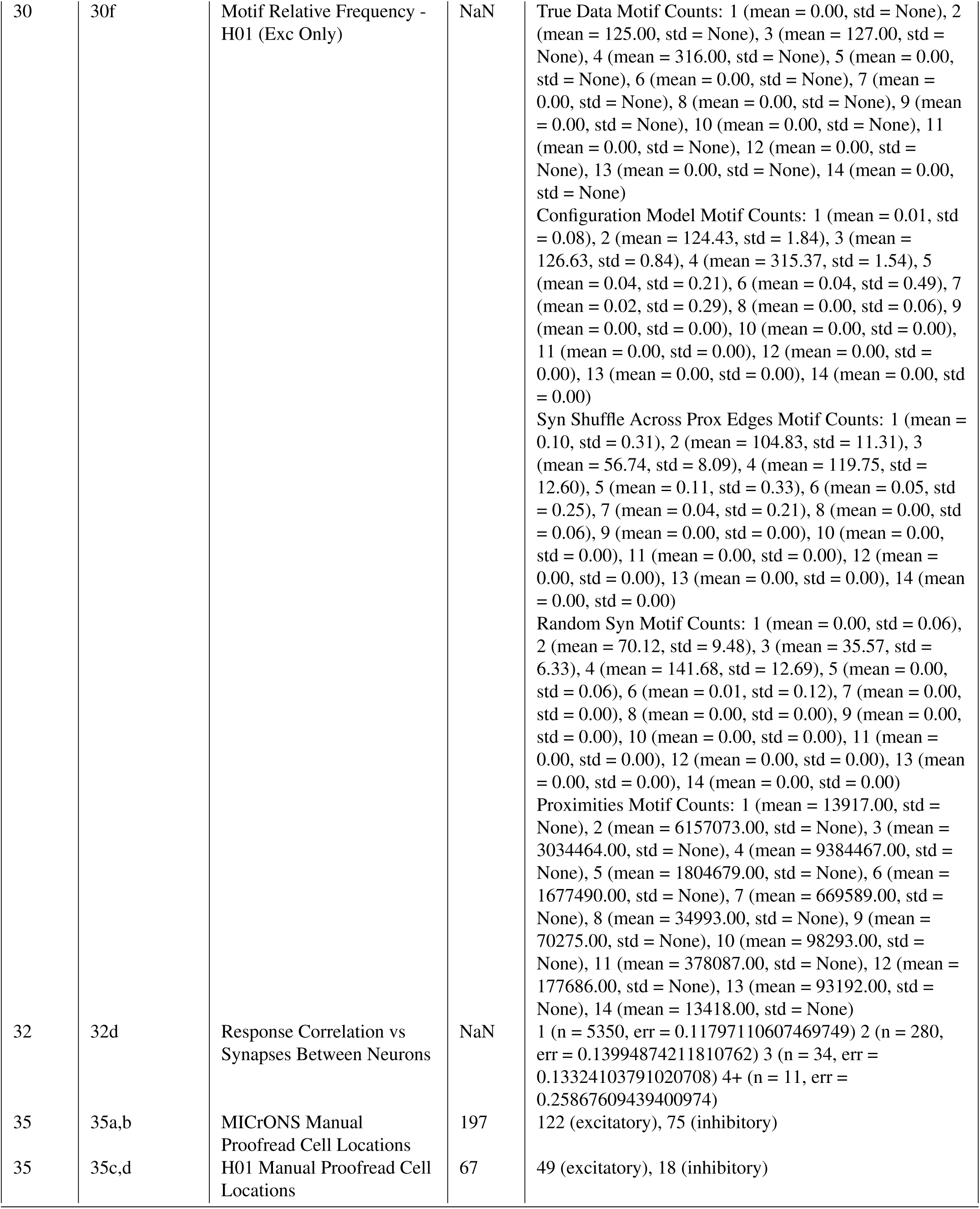
Figure Data Size. **(N)**

| Figure | Panel | Title / Concept | N | Subclass N |
| --- | --- | --- | --- | --- |
| NaN | NaN | NEURD pipeline: mesh objects with merged nuclei detected (MICrONS) | 115230 | NaN |
| NaN | NaN | NEURD pipeline: soma detection count (MICrONS) | 76894 | NaN |
| NaN | NaN | NEURD pipeline: single soma auto-proofread neurons, after non-neuronal filter (MICrONS) | 54802 | 49073 (Exc), 5729 (Inh) |
| NaN | NaN | NEURD pipeline: mesh objects with merged nuclei detected (H01) | 15068 | NaN |
| NaN | NaN | NEURD pipeline: soma detection count (H01) | 14303 | NaN |
| NaN | NaN | NEURD pipeline: single soma auto-proofread neurons, after non-neuronal filter (H01) | 11444 | 7357 (Exc), 4087 (Inh) |
| 3, 14 | MICrONS | NaN | NaN | Test Set: 122 (excitatory), 75 (inhibitory)<br>Automatic Proofreading Validation Totals:<br>for excitatory axon synapses: tp = 2216, fp = 324, tn = 24430, fn = 1420 for excitatory dendrite synapses: tp = 525072, fp = 3207, tn = 91423, fn = 4321 for inhibitory axon synapses: tp = 66372, fp = 6097, tn = 57164, fn = 47466 for inhibitory dendrite synapses: tp = 450043, fp = 901, tn = 44940, fn = 4625<br>for excitatory axon skeleton (um): tp = 85128.69, fp = 3797.15, tn = 87424.84, fn = 19831.9 for excitatory dendrite skeleton (um): tp = 466420.83, fp = 7769.81, tn = 52366.92, fn = 7644.89 for inhibitory axon skeleton (um): tp = 239772.53, fp = 23142.88, tn = 238705.66, fn = 90304.64 for inhibitory dendrite skeleton (um): tp = 184898.31, fp = 2695.24, tn = 140840.23, fn = 5136.95<br>No Proofreading Validation Totals:<br>for excitatory axon synapses: tp = 3636, fp = 24754, tn = 0, fn = 0 for excitatory dendrite synapses: tp = 529393, fp = 94630, tn = 0, fn = 0 for inhibitory axon synapses: tp = 113838, fp = 63261, tn = 0, fn = 0 for inhibitory dendrite synapses: tp = 454668, fp = 45841, tn = 0, fn = 0<br>for excitatory axon skeleton (um): tp = 111903.43, fp = 84279.14, tn = 0.0, fn = 3305.89 for excitatory dendrite skeleton (um): tp = 472820.13, fp = 61382.31, tn = 0.0, fn = 6629.94 for inhibitory axon skeleton (um): tp = 354128.6, fp = 237797.11, tn = 0.0, fn = 60760.66 for inhibitory dendrite skeleton (um): tp = 190354.31, fp = 143216.42, tn = 0.0, fn = 4426.28 |
| 3, 16 | H01 | NaN | NaN | Test Set: 49 (excitatory), 18 (inhibitory)<br>Automatic Proofreading Validation Totals:<br>for excitatory axon synapses: tp = 154, fp = 17, tn = 1263, fn = 140 for excitatory dendrite synapses: tp = 57181, fp = 4582, tn = 2882, fn = 14213 for inhibitory axon synapses: tp = 39, fp = 4, tn = 746, fn = 26 for inhibitory dendrite synapses: tp = 25312, fp = 2773, tn = 363, fn = 2927<br>for excitatory axon skeleton (um): tp = 8149.14, fp = 269.81, tn = 60.51, fn = 2543.82 for excitatory dendrite skeleton (um): tp = 84499.71, fp = 1700.02, tn = 608.63, fn = 21687.95 for inhibitory axon skeleton (um): tp = 2195.75, fp = 141.39, tn = 153.0, fn = 403.54 for inhibitory dendrite skeleton (um): tp = 13287.63, fp = 423.42, tn = 84.65, fn = 878.56<br>No Proofreading Validation Totals:<br>for excitatory axon synapses: tp = 294, fp = 1280, tn = 0, fn = 0 for excitatory dendrite synapses: tp = 71394, fp = 7464, tn = 0, fn = 0 for inhibitory axon synapses: tp = 65, fp = 750, tn = 0, fn = 0 for inhibitory dendrite synapses: tp = 28239, fp = 3136, tn = 0, fn = 0 for excitatory axon skeleton (um): tp = 10381.38, fp = 520.88, tn = 0.0, fn = 643.22 for excitatory dendrite skeleton (um): tp = 101312.97, fp = 5966.07, tn = 0.0, fn = 6376.24 for inhibitory axon skeleton (um): tp = 2446.3, fp = 141.39, tn = 0.0, fn = 186.16 for inhibitory dendrite skeleton (um): tp = 13624.78, fp = 1049.47, tn = 0.0, fn = 666.19 |
| 4 | 4a | Depth vs Synapses onto AIS (MICrONS) | 22955 | 0.08 (447), 0.16 (2646), 0.24 (3250), 0.31 (4023), 0.39 (3463), 0.46 (2048), 0.54 (2321), 0.61 (2749), 0.69 (1730), 0.77 (278) |
| 4 | 4d | Spine Head Syn Size vs. Spine Head Volume (H01) | 73084 | excitatory (71402), inhibitory (1682) |
| 4 | 4h | Angle of Thickest Basal (MICrONS) | 41309 | NaN |
| 4 | 4i | Angle of Thickest Basal (H01) | 3445 | NaN |
| 4 | 4j | Average Segment Length Basal | 33049 | 23P (13516), 5P-NP (398), Other Exc (19135) |
| 4 | 4k | Shaft Synapses to Spine Ratio | 30910 | 23P (11970), 4P (5443), 5P-ET (462), 5P-NP (216), Other Exc (12819) |
| 4 | 4l | Spine Neck Length | 41898 | 4P (7034), 5P-NP (461), Other Exc (34403) |
| 4 | 4m | Spine Syn Size vs. Spine Head Volume | 45783 | 5P-ET (846), 5P-NP (500), MC (973), Other Exc (40594), Other Inh (2870) |
| 5 | 5b | Distance To Soma (MICrONS) Basal Dendrite | 756521 | Exc Onto Exc (113052), Exc Onto Inh (106834), Inh Onto Exc (481022), Inh Onto Inh (55613) |
| 5 | 5c | Conversion Rate vs. Axon Distance to Soma (MICrONS) | NaN | Exc Onto Exc: 37.02 (prox = 3626042), 86.29 (prox = 3154321), 110.11 (prox = 3124147), 133.45 (prox = 3101303), 159.94 (prox = 3031230), 192.1 (prox = 2898453), 232.13 (prox = 2632417), 282.1 (prox = 2295182), 347.59 (prox = 1994158), 480.32 (prox = 1860907) Exc Onto Inh: 39.23 (prox = 266045), 86.32 (prox = 266044), 110.14 (prox = 266045), 133.47 (prox = 266050), 160.03 (prox = 266046), 192.32 (prox = 266039), 232.61 (prox = 266050), 282.8 (prox = 266045), 348.3 (prox = 266048), 478.41 (prox = 256818) |
| 5 | 5d | Conversion Rate (H01) | NaN | Inh Onto Exc: 0.15 (prox=5078, syn = 129), 0.3 (prox=9471, syn = 221), 0.45 (prox=13806, syn = 293), 0.6 (prox=17732, syn = 353), 0.75 (prox=21420, syn = 390), 0.9 (prox=24885, syn = 409), 1.05 (prox=28267, syn = 429), 1.2 (prox=31601, syn = 444), 1.35 (prox=34865, syn = 453), 1.5 (prox=38048, syn = 460), 1.65 (prox=41280, syn = 468), 1.8 (prox=44520, syn = 472), 1.95 (prox=47676, syn = 478), 2.1 (prox=50857, syn = 479), 2.25 (prox=54149, syn = 484), 2.4 (prox=57369, syn = 488), 2.55 (prox=60569, syn = 491), 2.7 (prox=63750, syn = 495), 2.85 (prox=67008, syn = 499), 3.0 (prox=70398, syn = 500)<br>Inh Onto Inh: 0.15 (prox=720, syn = 58), 0.3 (prox=1576, syn = 116), 0.45 (prox=2511, syn = 165), 0.6 (prox=3417, syn = 204), 0.75 (prox=4249, syn = 226), 0.9 (prox=5054, syn = 237), 1.05 (prox=5857, syn = 244), 1.2 (prox=6615, syn = 256), 1.35 (prox=7410, syn = 263), 1.5 (prox=8243, syn = 264), 1.65 (prox=9015, syn = 266), 1.8 (prox=9818, syn = 267), 1.95 (prox=10556, syn = 269), 2.1 (prox=11321, syn = 271), 2.25 (prox=12125, syn = 271), 2.4 (prox=12912, syn = 272), 2.55 (prox=13675, syn = 273), 2.7 (prox=14458, syn = 273), 2.85 (prox=15234, syn = 273), 3.0 (prox=16078, syn = 273)<br>Exc Onto Inh: 0.15 (prox=3234, syn = 938), 0.3 (prox=6774, syn = 1820), 0.45 (prox=10224, syn = 2403), 0.6 (prox=13301, syn = 2750), 0.75 (prox=16304, syn = 3015), 0.9 (prox=19103, syn = 3180), 1.05 (prox=21867, syn = 3299), 1.2 (prox=24485, syn = 3388), 1.35 (prox=27127, syn = 3449), 1.5 (prox=29705, syn = 3507), 1.65 (prox=32217, syn = 3539), 1.8 (prox=34790, syn = 3567), 1.95 (prox=37352, syn = 3591), 2.1 (prox=39874, syn = 3603), 2.25 (prox=42505, syn = 3618), 2.4 (prox=45124, syn = 3627), 2.55 (prox=47696, syn = 3632), 2.7 (prox=50393, syn = 3635), 2.85 (prox=53031, syn = 3638), 3.0 (prox=55636, syn = 3641)<br>Exc Onto Exc: 0.15 (prox=19083, syn = 114), 0.3 (prox=35936, syn = 210), 0.45 (prox=52776, syn = 274), 0.6 (prox=67950, syn = 332), 0.75 (prox=82327, syn = 379), 0.9 (prox=96308, syn = 416), 1.05 (prox=109762, syn = 444), 1.2 (prox=123090, syn = 481), 1.35 (prox=136144, syn = 513), 1.5 (prox=148976, syn = 540), 1.65 (prox=161807, syn = 573), 1.8 (prox=174565, syn = 586), 1.95 (prox=187166, syn = 608), 2.1 (prox=199647, syn = 623), 2.25 (prox=212400, syn = 639), 2.4 (prox=224934, syn = 645), 2.55 (prox=237539, syn = 658), 2.7 (prox=250354, syn = 665), 2.85 (prox=263115, syn = 675), 3.0 (prox=276150, syn = 684) |
| 5 | 5e | Conversion Rate (MICrONS) | NaN | Exc Onto Inh: 0.15 (prox=54950, syn = 8628), 0.3 (prox=123804, syn = 17121), 0.45 (prox=198634, syn = 23931), 0.6 (prox=266713, syn = 28392), 0.75 (prox=328434, syn = 31149), 0.9 (prox=385586, syn = 32852), 1.05 (prox=440163, syn = 33970), 1.2 (prox=492759, syn = 34676), 1.35 (prox=543632, syn = 35143), 1.5 (prox=593594, syn = 35438), 1.65 (prox=643038, syn = 35658), 1.8 (prox=692573, syn = 35814), 1.95 (prox=741389, syn = 35912), 2.1 (prox=790363, syn = 36017), 2.25 (prox=839090, syn = 36086), 2.4 (prox=888119, syn = 36148), 2.55 (prox=937351, syn = 36211), 2.7 (prox=986303, syn = 36264), 2.85 (prox=1035527, syn = 36300), 3.0 (prox=1085128, syn = 36322) Inh Onto Inh: 0.15 (prox=23894, syn = 4791), 0.3 (prox=50350, syn = 8435), 0.45 (prox=77195, syn = 11298), 0.6 (prox=100582, syn = 12893), 0.75 (prox=121587, syn = 13823), 0.9 (prox=140984, syn = 14278), 1.05 (prox=159432, syn = 14545), 1.2 (prox=177191, syn = 14691), 1.35 (prox=194885, syn = 14802), 1.5 (prox=212290, syn = 14878), 1.65 (prox=229167, syn = 14937), 1.8 (prox=245931, syn = 14977), 1.95 (prox=262711, syn = 15015), 2.1 (prox=279523, syn = 15050), 2.25 (prox=296300, syn = 15081), 2.4 (prox=312730, syn = 15104), 2.55 (prox=329411, syn = 15121), 2.7 (prox=346209, syn = 15134), 2.85 (prox=362732, syn = 15144), 3.0 (prox=379638, syn = 15162) Inh Onto Exc: 0.15 (prox=431892, syn = 73682), 0.3 (prox=811565, syn = 121599), 0.45 (prox=1170806, syn = 157540), 0.6 (prox=1479467, syn = 180273), 0.75 (prox=1757304, syn = 193944), 0.9 (prox=2016992, syn = 202263), 1.05 (prox=2266874, syn = 208146), 1.2 (prox=2509814, syn = 212482), 1.35 (prox=2747815, syn = 215775), 1.5 (prox=2980676, syn = 218563), 1.65 (prox=3210694, syn = 220807), 1.8 (prox=3439389, syn = 222649), 1.95 (prox=3666897, syn = 224084), 2.1 (prox=3893507, syn = 225230), 2.25 (prox=4119930, syn = 226138), 2.4 (prox=4346589, syn = 226835), 2.55 (prox=4572534, syn = 227462), 2.7 (prox=4800020, syn = 227975), 2.85 (prox=5028673, syn = 228358), 3.0 (prox=5257983, syn = 228664) Exc Onto Exc: 0.15 (prox=770871, syn = 8901), 0.3 (prox=1543245, syn = 16617), 0.45 (prox=2324856, syn = 23468), 0.6 (prox=3025906, syn = 28951), 0.75 (prox=3668135, syn = 34040), 0.9 (prox=4272087, syn = 38609), 1.05 (prox=4851252, syn = 42764), 1.2 (prox=5412367, syn = 46525), 1.35 (prox=5960708, syn = 49759), 1.5 (prox=6500095, syn = 52533), 1.65 (prox=7033215, syn = 54881), 1.8 (prox=7562021, syn = 56768), 1.95 (prox=8086547, syn = 58236), 2.1 (prox=8609337, syn = 59396), 2.25 (prox=9133249, syn = 60301), 2.4 (prox=9654903, syn = 60936), 2.55 (prox=10178601, syn = 61447), 2.7 (prox=10702652, syn = 61791), 2.85 (prox=11230246, syn = 62046), 3.0 (prox=11760681, syn = 62251) |
| 5 | 5f | Cell Type Conversion Rates | NaN | 23P Onto 5P-ET (225805), Other Exc Onto Exc (3994769), 23P Onto MC (64475), 23P/4P/5P-IT Onto BC (217221), Exc Onto MC (128303), 23P Onto BC Soma (2145), Other Exc Onto Inh (95921), BC Onto 23P/5P-ET Dendr (786459), BC Onto Exc Soma (18696), MC Onto 23P/5P-ET Dendr (201853), Other Inh Onto Exc (947601), BC Onto BC Soma (950), BC Onto Inh Soma (2140), BPC Onto MC (1837), MC Onto BPC (5496), Other Inh Onto Inh (152469) |
| 5 | 5g | Motif index | NaN | True Data Motif Counts: 1 (mean = 7377.00, std = None), 2 (mean = 10953501.00, std = None), 3 (mean = 3930648.00, std = None), 4 (mean = 57770467.00, std = None), 5 (mean = 390627.00, std = None), 6 (mean = 3350392.00, std = None), 7 (mean = 383004.00, std = None), 8 (mean = 15873.00, std = None), 9 (mean = 66407.00, std = None), 10 (mean = 16156.00, std = None), 11 (mean = 48173.00, std = None), 12 (mean = 9957.00, std = None), 13 (mean = 5003.00, std = None), 14 (mean = 243.00, std = None)<br>Configuration Model Motif Counts: 1 (mean = 542.35, std = 22.87), 2 (mean = 14299374.71, std = 27655.61), 3 (mean = 4518749.52, std = 3704.19), 4 (mean = 59388828.02, std = 41381.50), 5 (mean = 166468.05, std = 1221.97), 6 (mean = 457613.39, std = 21497.53), 7 (mean = 54988.81, std = 2388.76), 8 (mean = 10084.60, std = 167.97), 9 (mean = 1126.76, std = 122.16), 10 (mean = 1601.45, std = 91.37), 11 (mean = 4437.23, std = 291.20), 12 (mean = 825.09, std = 54.10), 13 (mean = 113.36, std = 18.21), 14 (mean = 1.89, std = 1.40)<br>Syn Shuffle Across Prox Edges Motif Counts: 1 (mean = 480.44, std = 22.40), 2 (mean = 6279304.00, std = 14980.70), 3 (mean = 3868010.65, std = 6644.99), 4 (mean = 9647512.96, std = 18046.76), 5 (mean = 42992.30, std = 248.85), 6 (mean = 38400.50, std = 2007.61), 7 (mean = 15842.36, std = 810.50), 8 (mean = 2393.19, std = 58.46), 9 (mean = 51.05, std = 9.23), 10 (mean = 85.18, std = 10.63), 11 (mean = 146.84, std = 14.67), 12 (mean = 73.36, std = 9.71), 13 (mean = 0.91, std = 0.89), 14 (mean = 0.00, std = 0.06)<br>Random Syn Motif Counts: 1 (mean = 68.34, std = 8.16), 2 (mean = 5312021.94, std = 13718.80), 3 (mean = 2656056.72, std = 1670.41), 4 (mean = 10637406.97, std = 12420.92), 5 (mean = 6338.19, std = 86.42), 6 (mean = 6363.58, std = 765.06), 7 (mean = 1589.55, std = 193.60), 8 (mean = 528.37, std = 22.31), 9 (mean = 0.90, std = 0.91), 10 (mean = 1.87, std = 1.45), 11 (mean = 3.64, std = 2.12), 12 (mean = 0.87, std = 0.94), 13 (mean = 0.00, std = 0.06), 14 (mean = 0.00, std = 0.00)<br>Proximities Motif Counts: 1 (mean = 970833.00, std = None), 2 (mean = 5902464653.00, std = None), 3 (mean = 3956073180.00, std = None), 4 (mean = 13851915701.00, std = None), 5 (mean = 2218025598.00, std = None), 6 (mean = 1999665781.00, std = None), 7 (mean = 534201392.00, std = None), 8 (mean = 20112550.00, std = None), 9 (mean = 75994000.00, std = None), 10 (mean = 83402913.00, std = None), 11 (mean = 461402658.00, std = None), 12 (mean = 167761891.00, std = None), 13 (mean = 92029299.00, std = None), 14 (mean = 13916043.00, std = None) |
| 5 | 5i | Response Correlation vs Synapses Between Neurons | NaN | 1 (n = 5350, err = 0.11797110607469749) 2 (n = 280, err = 0.13994874211810762) 3 (n = 34, err = 0.13324103791020708) 4+ (n = 11, err = 0.25867609439400974) |
| 7 | 7a | Compartment Skeletal Length Validation (MICrONS) | 158 Neurons | Axon (67.01 mm), Basal (245.11 mm), Apical Trunk (52.17 mm), Oblique (59.51 mm), Apical (43.95 mm), Apical Tuft (24.20 mm) |
| 7 | 7b | Compartment Skeletal Length Validation (H01) | 89 Neurons | Axon (21.99 mm), Basal (131.60 mm), Apical Trunk (19.12 mm), Oblique (13.61 mm), Apical (34.29 mm), Apical Tuft (24.36 mm) |
| 9 | 9b | Classification Percentage vs. Double Back Rule Threshold | 1991 | NaN |
| 9 | 9c | Classification Percentage vs. Width Jump Ratio | 468 | NaN |
| 17 | 17a | Logistic Reg - E/I Model (MICrONS) | 4986 | 4024 (excitatory), 961 (inhibitory) |
| 17 | 17b | Logistic Reg - E/I Model (H01) | 7555 | 5800 (excitatory), 1755 (inhibitory) |
| 17 | 17d | Distribution of Predicted Exc/Inh Neurons By Layer Depth | 49117 | 23P (18720), 4P (7399), 5P-IT (7437), 5P-NP (514), 5P-PT (1232), 6P-CT (4975), 6P-IT (4944), BC (1425), BPC (946), MC (987), NGC (538) |
| 17 | 17e | GNN Classifier - Whole Neuron | 178 | 23P (41), 4P (40), 5P-IT (20), 5P-NP (5), 5P-ET (6), 6P-CT (20), 6P-IT (22), BC (11), BPC (9), MC (1), NGC (3) |
| 17 | 17f | GNN Classifier - Stems | 1023 | 23P (203), 4P (379), 5P-IT (62), 5P-NP (14), 5P-ET (49), 6P-CT (104), 6P-IT (92), BC (56), BPC (33), MC (20), NGC (11) |
| 20 | 20a | Spine Density | 33653 | 23P (11694), 4P (5409), MC (682), Other Exc (13781), Other Inh (2087) |
| 20 | 20b | Soma Volume | 33823 | 4P (5445), 5P-ET (384), 5P-NP (388), BC (1012), BPC (693), Other Exc (24856), Other Inh (1045) |
| 20 | 20c | Apical Trunk Starting Width | 27577 | 4P (5525), 5P-ET (503), 5P-NP (363), Other Exc (21186) |
| 20 | 20d | Synapses Onto AIS vs. Cell Type | 25060 | 23P (8787), 4P (4011), 5P-IT (4839), 5P-NP (251), 5P-PT (738), 6P-CT (2320), 6P-IT (2233), BC (817), BPC (355), MC (490), NGC (219) |
| 20 | 20e | Average Segment Length (Dendrite) | 2768 | BC (1045), BPC (697), MC (673), NGC (353) |
| 20 | 20f | Soma Synapses | 33655 | 23P (11976), 5P-ET (506), 5P-NP (388), BC (709), NGC (364), Other Exc (18339), Other Inh (1373) |
| 20 | 20g | Axon Starting Width | 39051 | 5P-ET (1273), 5P-NP (365), BPC (648), Other Exc (34664), Other Inh (2101) |
| 20 | 20h | Average Segment Length Basal | 33049 | 23P (13516), 5P-NP (398), Other Exc (19135) |
| 20 | 20i | Spine Neck Length | 41898 | 4P (7034), 5P-NP (461), Other Exc (34403) |
| 20 | 20j | Shaft Synapses to Spine Ratio | 30910 | 23P (11970), 4P (5443), 5P-ET (462), 5P-NP (216), Other Exc (12819) |
| 20 | 20k | Spine Syn Size vs. Spine Head Volume | 45783 | 5P-ET (846), 5P-NP (500), MC (973), Other Exc (40594), Other Inh (2870) |
| 21 | 21c | Depth Bin (MICrONS) | 22955 | 0.08 (447), 0.16 (2646), 0.24 (3250), 0.31 (4023), 0.39 (3463), 0.46 (2048), 0.54 (2321), 0.61 (2749), 0.69 (1730), 0.77 (278) |
| 21 | 21d | Depth Bin (H01) | 3154 | 0.58 (385), 0.84 (464), 1.09 (355), 1.35 (528), 1.60 (529), 1.86 (426), 2.11 (224), 2.37 (160), 2.62 (83) |
| 21 | 21e | Depth vs Synapses onto Soma (MICrONS) | 22955 | 0.08 (447), 0.16 (2646), 0.24 (3250), 0.31 (4023), 0.39 (3463), 0.46 (2048), 0.54 (2321), 0.61 (2749), 0.69 (1730), 0.77 (278) |
| 21 | 21f | Depth vs Synapses onto Soma (H01) | 3154 | 0.58 (385), 0.84 (464), 1.09 (355), 1.35 (528), 1.60 (529), 1.86 (426), 2.11 (224), 2.37 (160), 2.62 (83) |
| 21 | 21g | Depth vs Synapses onto AIS (MICrONS) | 22955 | 0.08 (447), 0.16 (2646), 0.24 (3250), 0.31 (4023), 0.39 (3463), 0.46 (2048), 0.54 (2321), 0.61 (2749), 0.69 (1730), 0.77 (278) |
| 21 | 21h | Depth vs Synapses onto AIS (H01) | 3154 | 0.58 (385), 0.84 (464), 1.09 (355), 1.35 (528), 1.60 (529), 1.86 (426), 2.11 (224), 2.37 (160), 2.62 (83) |
| 21 | 21i | Synapses Onto AIS (Excitatory Cells) | 26327 | H01 (3208), MICrONS (23119) |
| 21 | 21j | Synapses Onto Soma (Excitatory Cells) | 26327 | H01 (3208), MICrONS (23119) |
| 22 | 22b | Angle of Thickest Basal (MICrONS) | 41309 | NaN |
| 22 | 22c | Angle of Thickest Basal (H01) | 3445 | NaN |
| 23 | 23a | Number of Leaf Nodes Dendrite | 2721 | BPC (654), NGC (342), Other Inh (1725) |
| 23 | 23b | Leaf Nodes vs. Initial Width Basal/Dendrite Stems (MICrONS) | 197860 | NaN |
| 23 | 23c | Leaf Nodes vs. Initial Width Basal/Dendrite Stems (H01) | 30058 | NaN |
| 25 | 25a | Spine Head Syn Size vs. Spine Head Volume (MICrONS) | 240246 | excitatory (222009), inhibitory (18237) |
| 25 | 25b | Spine Head Syn Size vs. Spine Neck Width (MICrONS) | 240246 | excitatory (222009), inhibitory (18237) |
| 25 | 25c | Spine Neck Syn Size vs. Spine Neck Width (MICrONS) | 240246 | excitatory (222009), inhibitory (18237) |
| 25 | 25d | Spine Head Syn Size vs. Spine Neck Syn Size (MICrONS) | 240246 | excitatory (222009), inhibitory (18237) |
| 25 | 25e | Spine Head Syn Size vs. Spine Head Volume (MICrONS) | 240246 | excitatory (222009), inhibitory (18237) |
| 25 | 25f | Cell Type vs. Average Number of Syn on Spine (MICrONS) | 233981 | excitatory spines (n = 219476) inhibitory spines (n = 14505) |
| 25 | 25g | Spine Head Syn Size vs. Spine Head Volume (H01) | 73084 | excitatory (71402), inhibitory (1682) |
| 25 | 25h | Spine Head Syn Size vs. Spine Neck Width (H01) | 73084 | excitatory (71402), inhibitory (1682) |
| 25 | 25i | Spine Neck Syn Size vs. Spine Neck Width (H01) | 73084 | excitatory (71402), inhibitory (1682) |
| 25 | 25j | Spine Head Syn Size vs. Spine Neck Syn Size (H01) | 73084 | excitatory (71402), inhibitory (1682) |
| 25 | 25k | Spine Head Syn Size vs. Spine Head Volume (H01) | 73084 | excitatory (71402), inhibitory (1682) |
| 25 | 25l | Cell Type vs. Average Number of Syn on Spine Head (H01) | 58678 | excitatory spines (n = 57645) inhibitory spines (n = 1033) |
| 26 | 26a | Syn Size vs Spine Head Volume (MICrONS) | 159552 | Exc Onto Exc (76295), Inh Onto Exc (83257) |
| 26 | 26b | Syn Size vs Spine Head Volume (H01) | 159312 | Exc Onto Exc (76248), Inh Onto Exc (83064) |
| 26 | 26c | Spine Head Volume (MICrONS) | 152942 | Exc Onto Exc (73980), Inh Onto Exc (78962) |
| 26 | 26d | Spine Head Volume (H01) | 63440 | Exc Onto Exc (6855), Inh Onto Exc (56585) |
| 26 | 26e | Number of Syn on Spine Head (MICrONS) | 426 | Exc Onto Exc (346), Inh Onto Exc (80) |
| 26 | 26f | Number of Syn on Spine Head (H01) | 425 | Exc Onto Exc (345), Inh Onto Exc (80) |
| 26 | 26g | Max Syn vs Syn Size Ratio on Multi Synapse Spine Head (MICrONS) | 419 | Exc Onto Exc (343), Inh Onto Exc (76) |
| 26 | 26h | Max Syn vs Syn Size Ratio on Multi Synapse Spine Head (H01) | 83 | Exc Onto Exc (17), Inh Onto Exc (66) |
| 26 | 26i | Synapse Size (MICrONS) | 1011266 | Exc Onto Exc (164046), Exc Onto Inh (113229), Inh Onto Exc (675088), Inh Onto Inh (58903) |
| 26 | 26j | Synapse Size (H01) | 7343 | Exc Onto Exc (1055), Exc Onto Inh (4954), Inh Onto Exc (899), Inh Onto Inh (435) |
| 28 | 28b | Conversion Rate vs. Depth Diff (MICrONS) | NaN | Exc Onto Exc: -93.5 (prox = 708773), -16.61 (prox = 1076784), 15.58 (prox = 2987066), 39.74 (prox = 4235702), 59.36 (prox = 3999294), 80.32 (prox = 3945396), 115.64 (prox = 3548662), 170.01 (prox = 2960345), 231.13 (prox = 2408377), 306.08 (prox = 1827653) Exc Onto Inh: -89.23 (prox = 39518), -16.13 (prox = 67956), 15.8 (prox = 210778), 39.94 (prox = 339614), 59.37 (prox = 340694), 80.38 (prox = 341392), 116.18 (prox = 339496), 171.15 (prox = 337738), 231.95 (prox = 332885), 306.83 (prox = 289258) |
| 28 | 28c | Conversion Rate vs. Depth Diff (H01) | NaN | Exc Onto Exc: -103.97 (prox = 11034), -27.39 (prox = 14694), -4.95 (prox = 8450), 5.51 (prox = 16023), 14.89 (prox = 40616), 26.62 (prox = 43181), 38.69 (prox = 47070), 51.54 (prox = 53530), 67.14 (prox = 55479), 89.52 (prox = 57422), 126.91 (prox = 50020), 242.6 (prox = 59649) Exc Onto Inh: -104.09 (prox = 2286), -27.54 (prox = 3149), -4.91 (prox = 1748), 5.38 (prox = 3026), 14.85 (prox = 7583), 26.67 (prox = 8376), 38.77 (prox = 9764), 51.66 (prox = 11456), 67.1 (prox = 12138), 89.58 (prox = 12742), 127.41 (prox = 11994), 239.65 (prox = 14731) |
| 28 | 28d | Conversion Rate vs. Axon Distance to Soma (MICrONS) | NaN | Exc Onto Exc: 37.02 (prox = 3626042), 86.29 (prox = 3154321), 110.11 (prox = 3124147), 133.45 (prox = 3101303), 159.94 (prox = 3031230), 192.1 (prox = 2898453), 232.13 (prox = 2632417), 282.1 (prox = 2295182), 347.59 (prox = 1994158), 480.32 (prox = 1860907) Exc Onto Inh: 39.23 (prox = 266045), 86.32 (prox = 266044), 110.14 (prox = 266045), 133.47 (prox = 266050), 160.03 (prox = 266046), 192.32 (prox = 266039), 232.61 (prox = 266050), 282.8 (prox = 266045), 348.3 (prox = 266048), 478.41 (prox = 256818) |
| 28 | 28e | Conversion Rate vs. Axon Distance to Soma (H01) | NaN | Exc Onto Exc: 16.2 (prox = 56117), 52.59 (prox = 51054), 82.33 (prox = 48430), 105.72 (prox = 45839), 127.5 (prox = 45548), 150.26 (prox = 44634), 176.13 (prox = 42697), 208.56 (prox = 42372), 259.9 (prox = 40808), 406.54 (prox = 43756) Exc Onto Inh: 16.29 (prox = 9986), 53.1 (prox = 9985), 82.37 (prox = 9986), 105.74 (prox = 9985), 127.37 (prox = 9986), 150.25 (prox = 9986), 176.28 (prox = 9984), 208.69 (prox = 9986), 260.43 (prox = 9985), 400.13 (prox = 9987) |
| 28 | 28f | Distance To Soma (MICrONS) Basal Dendrite | 756521 | Exc Onto Exc (113052), Exc Onto Inh (106834), Inh Onto Exc (481022), Inh Onto Inh (55613) |
| 28 | 28g | Distance To Soma (H01) Basal Dendrite | 6454 | Exc Onto Exc (700), Exc Onto Inh (4845), Inh Onto Exc (491), Inh Onto Inh (418) |
| 30 | 30a | Motif Relative Frequency - MICrONS (All) | NaN | True Data Motif Counts: 1 (mean = 7377.00, std = None), 2 (mean = 10953501.00, std = None), 3 (mean = 3930648.00, std = None), 4 (mean = 57770467.00, std = None), 5 (mean = 390627.00, std = None), 6 (mean = 3350392.00, std = None), 7 (mean = 383004.00, std = None), 8 (mean = 15873.00, std = None), 9 (mean = 66407.00, std = None), 10 (mean = 16156.00, std = None), 11 (mean = 48173.00, std = None), 12 (mean = 9957.00, std = None), 13 (mean = 5003.00, std = None), 14 (mean = 243.00, std = None)<br>Configuration Model Motif Counts: 1 (mean = 542.35, std = 22.87), 2 (mean = 14299374.71, std = 27655.61), 3 (mean = 4518749.52, std = 3704.19), 4 (mean = 59388828.02, std = 41381.50), 5 (mean = 166468.05, std = 1221.97), 6 (mean = 457613.39, std = 21497.53), 7 (mean = 54988.81, std = 2388.76), 8 (mean = 10084.60, std = 167.97), 9 (mean = 1126.76, std = 122.16), 10 (mean = 1601.45, std = 91.37), 11 (mean = 4437.23, std = 291.20), 12 (mean = 825.09, std = 54.10), 13 (mean = 113.36, std = 18.21), 14 (mean = 1.89, std = 1.40)<br>Syn Shuffle Across Prox Edges Motif Counts: 1 (mean = 480.44, std = 22.40), 2 (mean = 6279304.00, std = 14980.70), 3 (mean = 3868010.65, std = 6644.99), 4 (mean = 9647512.96, std = 18046.76), 5 (mean = 42992.30, std = 248.85), 6 (mean = 38400.50, std = 2007.61), 7 (mean = 15842.36, std = 810.50), 8 (mean = 2393.19, std = 58.46), 9 (mean = 51.05, std = 9.23), 10 (mean = 85.18, std = 10.63), 11 (mean = 146.84, std = 14.67), 12 (mean = 73.36, std = 9.71), 13 (mean = 0.91, std = 0.89), 14 (mean = 0.00, std = 0.06)<br>Random Syn Motif Counts: 1 (mean = 68.34, std = 8.16), 2 (mean = 5312021.94, std = 13718.80), 3 (mean = 2656056.72, std = 1670.41), 4 (mean = 10637406.97, std = 12420.92), 5 (mean = 6338.19, std = 86.42), 6 (mean = 6363.58, std = 765.06), 7 (mean = 1589.55, std = 193.60), 8 (mean = 528.37, std = 22.31), 9 (mean = 0.90, std = 0.91), 10 (mean = 1.87, std = 1.45), 11 (mean = 3.64, std = 2.12), 12 (mean = 0.87, std = 0.94), 13 (mean = 0.00, std = 0.06), 14 (mean = 0.00, std = 0.00)<br>Proximities Motif Counts: 1 (mean = 970833.00, std = None), 2 (mean = 5902464653.00, std = None), 3 (mean = 3956073180.00, std = None), 4 (mean = 13851915701.00, std = None), 5 (mean = 2218025598.00, std = None), 6 (mean = 1999665781.00, std = None), 7 (mean = 534201392.00, std = None), 8 (mean = 20112550.00, std = None), 9 (mean = 75994000.00, std = None), 10 (mean = 83402913.00, std = None), 11 (mean = 461402658.00, std = None), 12 (mean = 167761891.00, std = None), 13 (mean = 92029299.00, std = None), 14 (mean = 13916043.00, std = None) |
| 30 | 30b | Motif Relative Frequency - MICrONS (Inh Only) | NaN | True Data Motif Counts: 1 (mean = 620.00, std = None), 2 (mean = 153546.00, std = None), 3 (mean = 92977.00, std = None), 4 (mean = 248328.00, std = None), 5 (mean = 13517.00, std = None), 6 (mean = 24619.00, std = None), 7 (mean = 9023.00, std = None), 8 (mean = 686.00, std = None), 9 (mean = 621.00, std = None), 10 (mean = 769.00, std = None), 11 (mean = 1860.00, std = None), 12 (mean = 694.00, std = None), 13 (mean = 321.00, std = None), 14 (mean = 29.00, std = None)<br>Configuration Model Motif Counts: 1 (mean = 39.06, std = 5.93), 2 (mean = 205483.82, std = 581.96), 3 (mean = 116115.15, std = 209.37), 4 (mean = 284227.47, std = 572.75), 5 (mean = 2069.34, std = 51.22), 6 (mean = 2105.25, std = 356.40), 7 (mean = 843.10, std = 135.95), 8 (mean = 218.74, std = 15.63), 9 (mean = 4.56, std = 2.35), 10 (mean = 8.75, std = 3.32), 11 (mean = 12.07, std = 4.11), 12 (mean = 4.85, std = 2.49), 13 (mean = 0.13, std = 0.34), 14 (mean = 0.00, std = 0.00)<br>Syn Shuffle Across Prox Edges Motif Counts: 1 (mean = 173.79, std = 14.30), 2 (mean = 180258.18, std = 1674.63), 3 (mean = 90356.21, std = 609.90), 4 (mean = 223690.59, std = 1702.49), 5 (mean = 4441.91, std = 84.33), 6 (mean = 6021.68, std = 553.27), 7 (mean = 2684.63, std = 237.40), 8 (mean = 518.17, std = 26.97), 9 (mean = 43.93, std = 9.60), 10 (mean = 69.06, std = 9.87), 11 (mean = 83.00, std = 12.20), 12 (mean = 40.96, std = 6.96), 13 (mean = 3.09, std = 1.95), 14 (mean = 0.04, std = 0.19)<br>Random Syn Motif Counts: 1 (mean = 21.80, std = 4.84), 2 (mean = 155168.95, std = 1720.36), 3 (mean = 77526.32, std = 268.59), 4 (mean = 312209.08, std = 1758.50), 5 (mean = 1112.85, std = 33.99), 6 (mean = 1129.44, std = 254.99), 7 (mean = 277.83, std = 65.05), 8 (mean = 91.41, std = 9.20), 9 (mean = 1.07, std = 1.12), 10 (mean = 2.00, std = 1.44), 11 (mean = 4.13, std = 2.44), 12 (mean = 1.00, std = 0.99), 13 (mean = 0.01, std = 0.11), 14 (mean = 0.00, std = 0.00)<br>Proximities Motif Counts: 1 (mean = 35519.00, std = None), 2 (mean = 11523474.00, std = None), 3 (mean = 6692764.00, std = None), 4 (mean = 26319627.00, std = None), 5 (mean = 4298050.00, std = None), 6 (mean = 8808184.00, std = None), 7 (mean = 2352861.00, std = None), 8 (mean = 105187.00, std = None), 9 (mean = 625365.00, std = None), 10 (mean = 492474.00, std = None), 11 (mean = 2315658.00, std = None), 12 (mean = 553182.00, std = None), 13 (mean = 729469.00, std = None), 14 (mean = 196848.00, std = None) |
| 30 | 30c | Motif Relative Frequency - MICrONS (Exc Only) | NaN | True Data Motif Counts: 1 (mean = 51.00, std = None), 2 (mean = 349641.00, std = None), 3 (mean = 374553.00, std = None), 4 (mean = 934337.00, std = None), 5 (mean = 3004.00, std = None), 6 (mean = 2158.00, std = None), 7 (mean = 531.00, std = None), 8 (mean = 55.00, std = None), 9 (mean = 3.00, std = None), 10 (mean = 5.00, std = None), 11 (mean = 15.00, std = None), 12 (mean = 6.00, std = None), 13 (mean = 0.00, std = None), 14 (mean = 0.00, std = None)<br>Configuration Model Motif Counts: 1 (mean = 6.29, std = 2.49), 2 (mean = 354143.78, std = 192.71), 3 (mean = 375972.81, std = 249.71), 4 (mean = 933938.36, std = 970.55), 5 (mean = 542.67, std = 22.76), 6 (mean = 230.08, std = 124.94), 7 (mean = 89.97, std = 42.33), 8 (mean = 15.87, std = 3.63), 9 (mean = 0.02, std = 0.13), 10 (mean = 0.05, std = 0.23), 11 (mean = 0.14, std = 0.38), 12 (mean = 0.04, std = 0.22), 13 (mean = 0.00, std = 0.00), 14 (mean = 0.00, std = 0.00)<br>Syn Shuffle Across Prox Edges Motif Counts: 1 (mean = 23.94, std = 4.68), 2 (mean = 305884.63, std = 1550.10), 3 (mean = 210043.87, std = 892.90), 4 (mean = 467195.54, std = 1778.53), 5 (mean = 622.03, std = 25.46), 6 (mean = 408.27, std = 89.54), 7 (mean = 183.76, std = 41.90), 8 (mean = 20.45, std = 4.57), 9 (mean = 0.14, std = 0.36), 10 (mean = 0.24, std = 0.49), 11 (mean = 0.45, std = 0.67), 12 (mean = 0.24, std = 0.54), 13 (mean = 0.00, std = 0.00), 14 (mean = 0.00, std = 0.00)<br>Random Syn Motif Counts: 1 (mean = 3.92, std = 1.92), 2 (mean = 266889.39, std = 1546.80), 3 (mean = 133404.87, std = 397.34), 4 (mean = 534010.83, std = 1699.14), 5 (mean = 82.06, std = 9.40), 6 (mean = 86.25, std = 42.83), 7 (mean = 21.74, std = 11.83), 8 (mean = 6.75, std = 2.74), 9 (mean = 0.00, std = 0.06), 10 (mean = 0.00, std = 0.06), 11 (mean = 0.01, std = 0.12), 12 (mean = 0.01, std = 0.08), 13 (mean = 0.00, std = 0.00), 14 (mean = 0.00, std = 0.00)<br>Proximities Motif Counts: 1 (mean = 557816.00, std = None), 2 (mean = 3766065797.00, std = None), 3 (mean = 2705586217.00, std = None), 4 (mean = 8114160435.00, std = None), 5 (mean = 1469402757.00, std = None), 6 (mean = 821620414.00, std = None), 7 (mean = 262765795.00, std = None), 8 (mean = 6914301.00, std = None), 9 (mean = 21414903.00, std = None), 10 (mean = 27175777.00, std = None), 11 (mean = 222547569.00, std = None), 12 (mean = 100503502.00, std = None), 13 (mean = 32338897.00, std = None), 14 (mean = 4909292.00, std = None) |
| 30 | 30d | Motif Relative Frequency - H01 (All) | NaN | True Data Motif Counts: 1 (mean = 9.00, std = None), 2 (mean = 2641.00, std = None), 3 (mean = 9916.00, std = None), 4 (mean = 10139.00, std = None), 5 (mean = 65.00, std = None), 6 (mean = 79.00, std = None), 7 (mean = 49.00, std = None), 8 (mean = 0.00, std = None), 9 (mean = 0.00, std = None), 10 (mean = 0.00, std = None), 11 (mean = 2.00, std = None), 12 (mean = 0.00, std = None), 13 (mean = 0.00, std = None), 14 (mean = 0.00, std = None)<br>Configuration Model Motif Counts: 1 (mean = 0.20, std = 0.44), 2 (mean = 2815.22, std = 17.66), 3 (mean = 9884.73, std = 46.55), 4 (mean = 10177.57, std = 33.03), 5 (mean = 13.26, std = 3.94), 6 (mean = 2.47, std = 5.97), 7 (mean = 1.62, std = 4.67), 8 (mean = 0.09, std = 0.30), 9 (mean = 0.00, std = 0.00), 10 (mean = 0.00, std = 0.00), 11 (mean = 0.01, std = 0.08), 12 (mean = 0.00, std = 0.06), 13 (mean = 0.00, std = 0.00), 14 (mean = 0.00, std = 0.00)<br>Syn Shuffle Across Prox Edges Motif Counts: 1 (mean = 2.87, std = 1.69), 2 (mean = 2986.72, std = 80.70), 3 (mean = 1669.05, std = 51.16), 4 (mean = 3435.57, std = 94.71), 5 (mean = 8.83, std = 3.27), 6 (mean = 8.34, std = 6.05), 7 (mean = 4.67, std = 3.63), 8 (mean = 0.62, std = 0.77), 9 (mean = 0.01, std = 0.08), 10 (mean = 0.02, std = 0.13), 11 (mean = 0.01, std = 0.12), 12 (mean = 0.01, std = 0.08), 13 (mean = 0.00, std = 0.00), 14 (mean = 0.00, std = 0.00)<br>Random Syn Motif Counts: 1 (mean = 0.11, std = 0.32), 2 (mean = 1833.82, std = 60.04), 3 (mean = 911.83, std = 30.86), 4 (mean = 3662.59, std = 78.88), 5 (mean = 0.33, std = 0.62), 6 (mean = 0.40, std = 1.28), 7 (mean = 0.08, std = 0.40), 8 (mean = 0.02, std = 0.13), 9 (mean = 0.00, std = 0.00), 10 (mean = 0.00, std = 0.00), 11 (mean = 0.00, std = 0.00), 12 (mean = 0.00, std = 0.00), 13 (mean = 0.00, std = 0.00), 14 (mean = 0.00, std = 0.00)<br>Proximities Motif Counts: 1 (mean = 21240.00, std = None), 2 (mean = 10969177.00, std = None), 3 (mean = 6071205.00, std = None), 4 (mean = 16743583.00, std = None), 5 (mean = 2913183.00, std = None), 6 (mean = 3002536.00, std = None), 7 (mean = 1278298.00, std = None), 8 (mean = 64293.00, std = None), 9 (mean = 126906.00, std = None), 10 (mean = 175963.00, std = None), 11 (mean = 621990.00, std = None), 12 (mean = 281840.00, std = None), 13 (mean = 154254.00, std = None), 14 (mean = 21604.00, std = None) |
| 30 | 30e | Motif Relative Frequency - H01 (Inh Only) | NaN | True Data Motif Counts: 1 (mean = 1.00, std = None), 2 (mean = 27.00, std = None), 3 (mean = 40.00, std = None), 4 (mean = 163.00, std = None), 5 (mean = 1.00, std = None), 6 (mean = 4.00, std = None), 7 (mean = 3.00, std = None), 8 (mean = 0.00, std = None), 9 (mean = 0.00, std = None), 10 (mean = 0.00, std = None), 11 (mean = 0.00, std = None), 12 (mean = 0.00, std = None), 13 (mean = 0.00, std = None), 14 (mean = 0.00, std = None)<br>Configuration Model Motif Counts: 1 (mean = 0.01, std = 0.08), 2 (mean = 36.15, std = 1.79), 3 (mean = 43.17, std = 1.26), 4 (mean = 166.47, std = 2.56), 5 (mean = 0.06, std = 0.25), 6 (mean = 0.02, std = 0.25), 7 (mean = 0.01, std = 0.16), 8 (mean = 0.00, std = 0.00), 9 (mean = 0.00, std = 0.00), 10 (mean = 0.00, std = 0.00), 11 (mean = 0.00, std = 0.00), 12 (mean = 0.00, std = 0.00), 13 (mean = 0.00, std = 0.00), 14 (mean = 0.00, std = 0.00)<br>Syn Shuffle Across Prox Edges Motif Counts: 1 (mean = 0.24, std = 0.46), 2 (mean = 26.57, std = 6.10), 3 (mean = 16.70, std = 4.37), 4 (mean = 36.09, std = 6.73), 5 (mean = 0.11, std = 0.34), 6 (mean = 0.15, std = 0.53), 7 (mean = 0.05, std = 0.25), 8 (mean = 0.01, std = 0.10), 9 (mean = 0.00, std = 0.00), 10 (mean = 0.00, std = 0.00), 11 (mean = 0.00, std = 0.00), 12 (mean = 0.00, std = 0.00), 13 (mean = 0.00, std = 0.00), 14 (mean = 0.00, std = 0.00)<br>Random Syn Motif Counts: 1 (mean = 0.00, std = 0.00), 2 (mean = 17.56, std = 4.43), 3 (mean = 8.63, std = 3.09), 4 (mean = 35.02, std = 6.35), 5 (mean = 0.00, std = 0.00), 6 (mean = 0.00, std = 0.00), 7 (mean = 0.00, std = 0.00), 8 (mean = 0.00, std = 0.00), 9 (mean = 0.00, std = 0.00), 10 (mean = 0.00, std = 0.00), 11 (mean = 0.00, std = 0.00), 12 (mean = 0.00, std = 0.00), 13 (mean = 0.00, std = 0.00), 14 (mean = 0.00, std = 0.00)<br>Proximities Motif Counts: 1 (mean = 857.00, std = None), 2 (mean = 49602.00, std = None), 3 (mean = 34848.00, std = None), 4 (mean = 94884.00, std = None), 5 (mean = 13094.00, std = None), 6 (mean = 16507.00, std = None), 7 (mean = 6447.00, std = None), 8 (mean = 315.00, std = None), 9 (mean = 743.00, std = None), 10 (mean = 967.00, std = None), 11 (mean = 3032.00, std = None), 12 (mean = 1171.00, std = None), 13 (mean = 795.00, std = None), 14 (mean = 115.00, std = None) |
| 30 | 30f | Motif Relative Frequency - H01 (Exc Only) | NaN | True Data Motif Counts: 1 (mean = 0.00, std = None), 2 (mean = 125.00, std = None), 3 (mean = 127.00, std = None), 4 (mean = 316.00, std = None), 5 (mean = 0.00, std = None), 6 (mean = 0.00, std = None), 7 (mean = 0.00, std = None), 8 (mean = 0.00, std = None), 9 (mean = 0.00, std = None), 10 (mean = 0.00, std = None), 11 (mean = 0.00, std = None), 12 (mean = 0.00, std = None), 13 (mean = 0.00, std = None), 14 (mean = 0.00, std = None)<br>Configuration Model Motif Counts: 1 (mean = 0.01, std = 0.08), 2 (mean = 124.43, std = 1.84), 3 (mean = 126.63, std = 0.84), 4 (mean = 315.37, std = 1.54), 5 (mean = 0.04, std = 0.21), 6 (mean = 0.04, std = 0.49), 7 (mean = 0.02, std = 0.29), 8 (mean = 0.00, std = 0.06), 9 (mean = 0.00, std = 0.00), 10 (mean = 0.00, std = 0.00), 11 (mean = 0.00, std = 0.00), 12 (mean = 0.00, std = 0.00), 13 (mean = 0.00, std = 0.00), 14 (mean = 0.00, std = 0.00)<br>Syn Shuffle Across Prox Edges Motif Counts: 1 (mean = 0.10, std = 0.31), 2 (mean = 104.83, std = 11.31), 3 (mean = 56.74, std = 8.09), 4 (mean = 119.75, std = 12.60), 5 (mean = 0.11, std = 0.33), 6 (mean = 0.05, std = 0.25), 7 (mean = 0.04, std = 0.21), 8 (mean = 0.00, std = 0.06), 9 (mean = 0.00, std = 0.00), 10 (mean = 0.00, std = 0.00), 11 (mean = 0.00, std = 0.00), 12 (mean = 0.00, std = 0.00), 13 (mean = 0.00, std = 0.00), 14 (mean = 0.00, std = 0.00)<br>Random Syn Motif Counts: 1 (mean = 0.00, std = 0.06), 2 (mean = 70.12, std = 9.48), 3 (mean = 35.57, std = 6.33), 4 (mean = 141.68, std = 12.69), 5 (mean = 0.00, std = 0.06), 6 (mean = 0.01, std = 0.12), 7 (mean = 0.00, std = 0.00), 8 (mean = 0.00, std = 0.00), 9 (mean = 0.00, std = 0.00), 10 (mean = 0.00, std = 0.00), 11 (mean = 0.00, std = 0.00), 12 (mean = 0.00, std = 0.00), 13 (mean = 0.00, std = 0.00), 14 (mean = 0.00, std = 0.00)<br>Proximities Motif Counts: 1 (mean = 13917.00, std = None), 2 (mean = 6157073.00, std = None), 3 (mean = 3034464.00, std = None), 4 (mean = 9384467.00, std = None), 5 (mean = 1804679.00, std = None), 6 (mean = 1677490.00, std = None), 7 (mean = 669589.00, std = None), 8 (mean = 34993.00, std = None), 9 (mean = 70275.00, std = None), 10 (mean = 98293.00, std = None), 11 (mean = 378087.00, std = None), 12 (mean = 177686.00, std = None), 13 (mean = 93192.00, std = None), 14 (mean = 13418.00, std = None) |
| 32 | 32d | Response Correlation vs Synapses Between Neurons | NaN | 1 (n = 5350, err = 0.11797110607469749) 2 (n = 280, err = 0.13994874211810762) 3 (n = 34, err = 0.13324103791020708) 4+ (n = 11, err = 0.25867609439400974) |
| 35 | 35a,b | MICrONS Manual Proofread Cell Locations | 197 | 122 (excitatory), 75 (inhibitory) |
| 35 | 35c,d | H01 Manual Proofread Cell Locations | 67 | 49 (excitatory), 18 (inhibitory) |

**Appendix Table 3:**
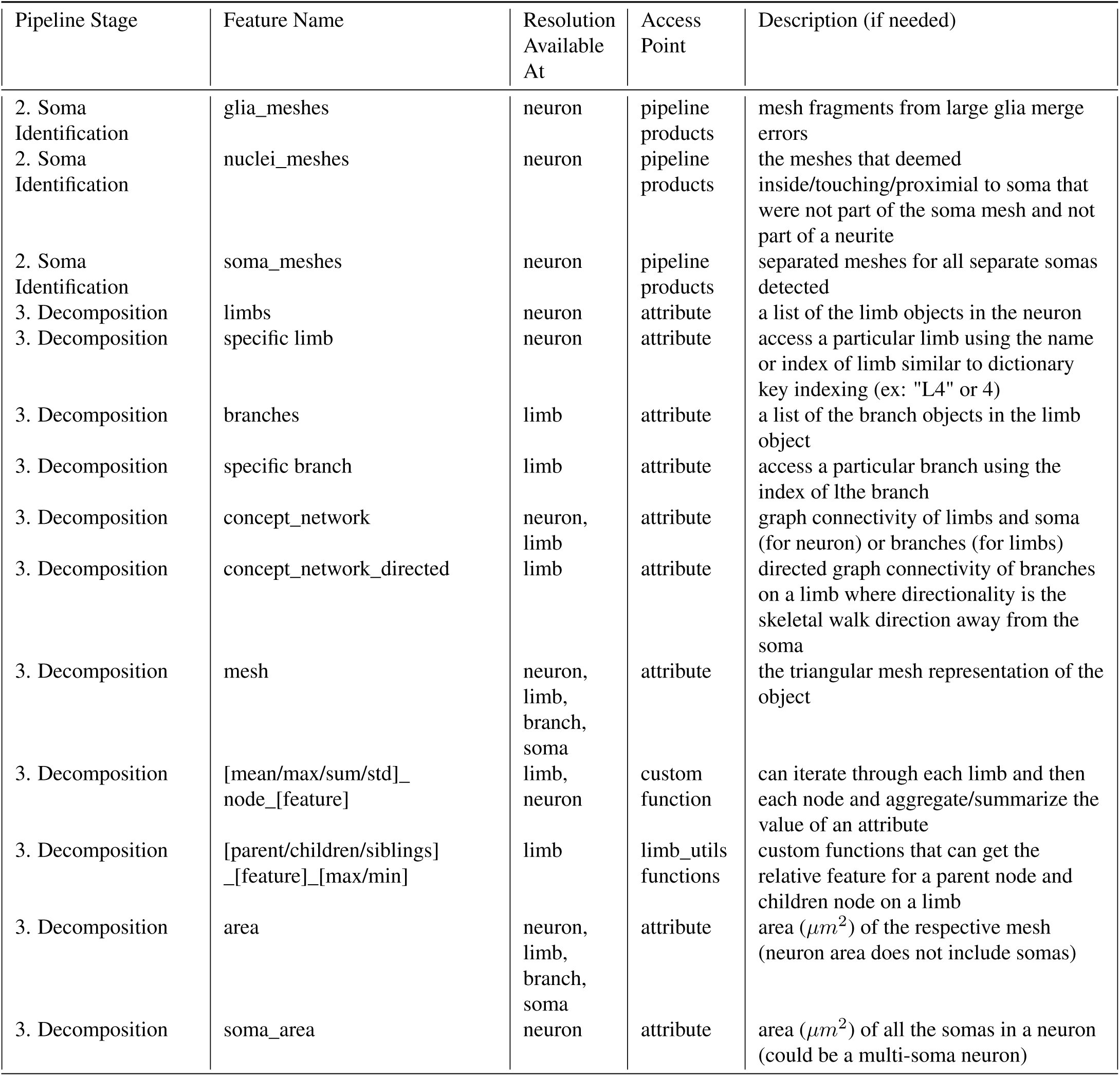
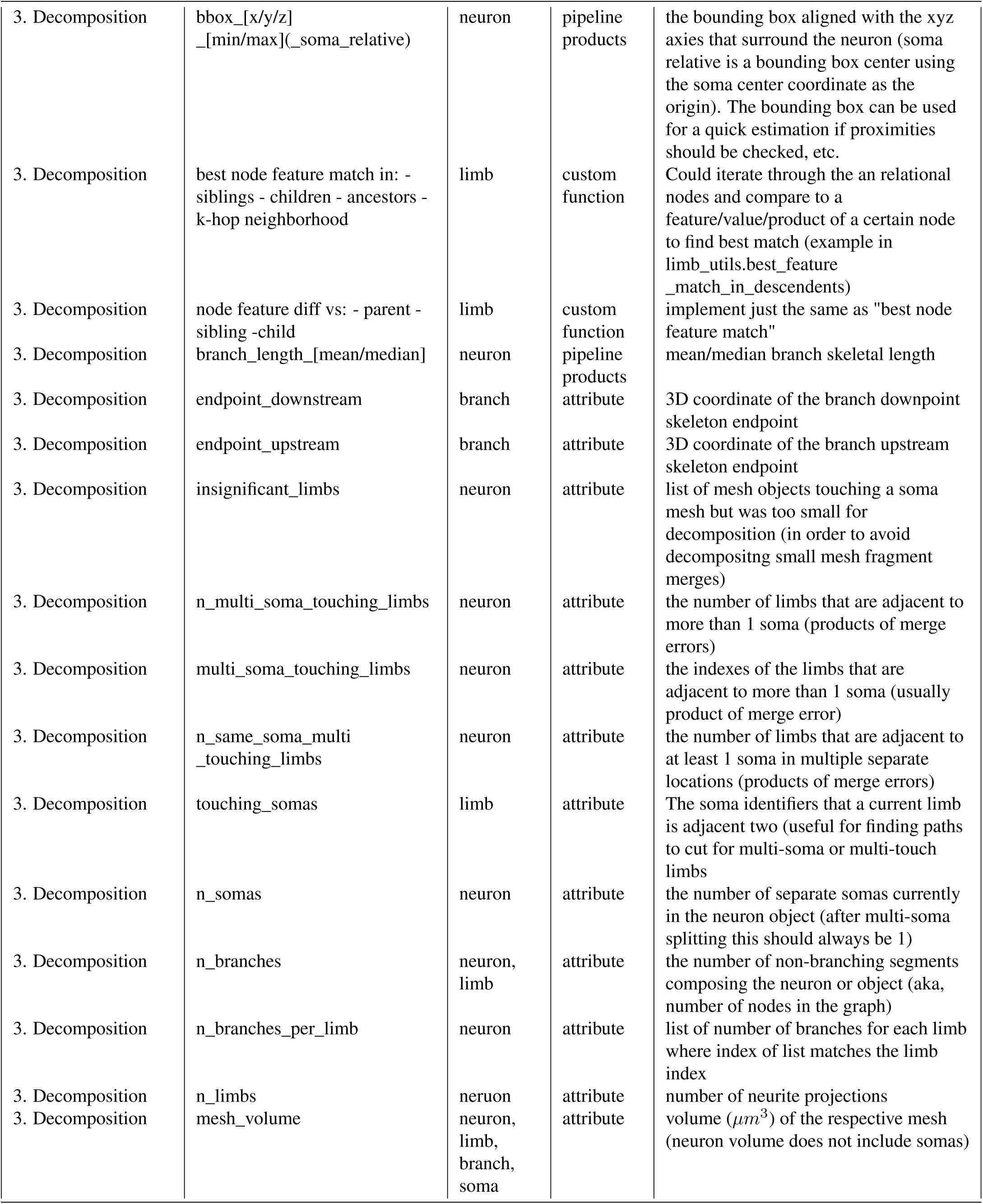
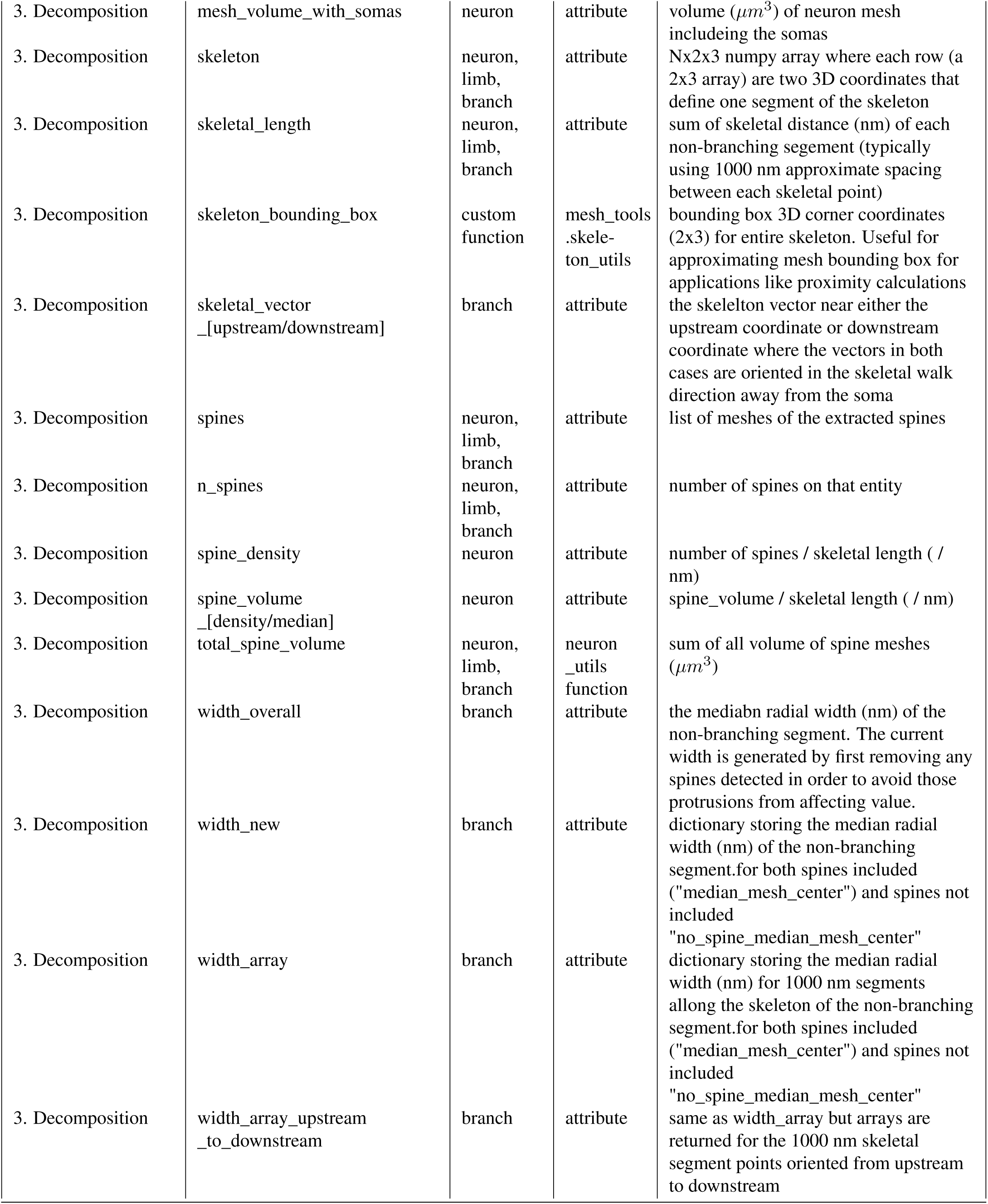
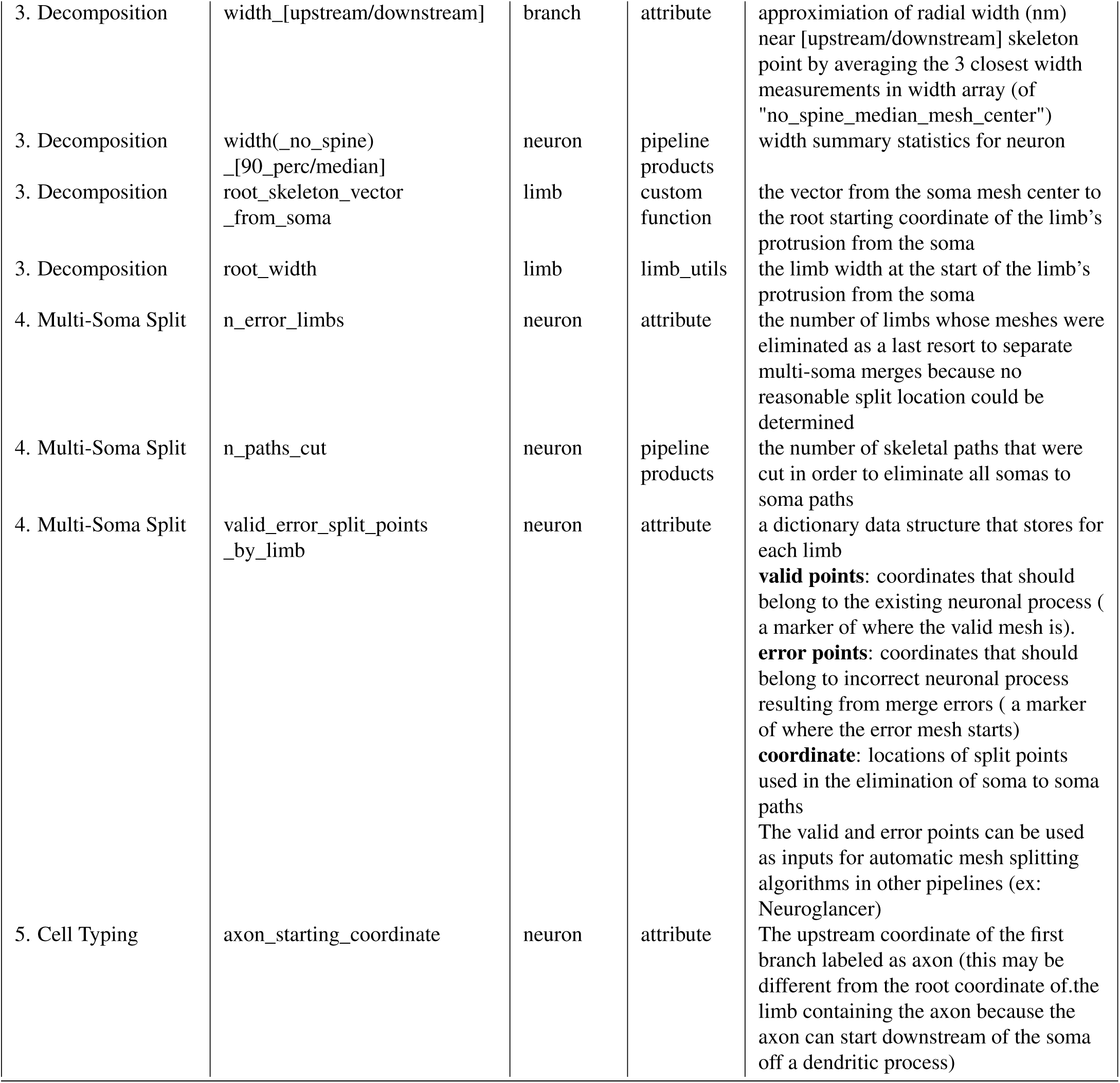
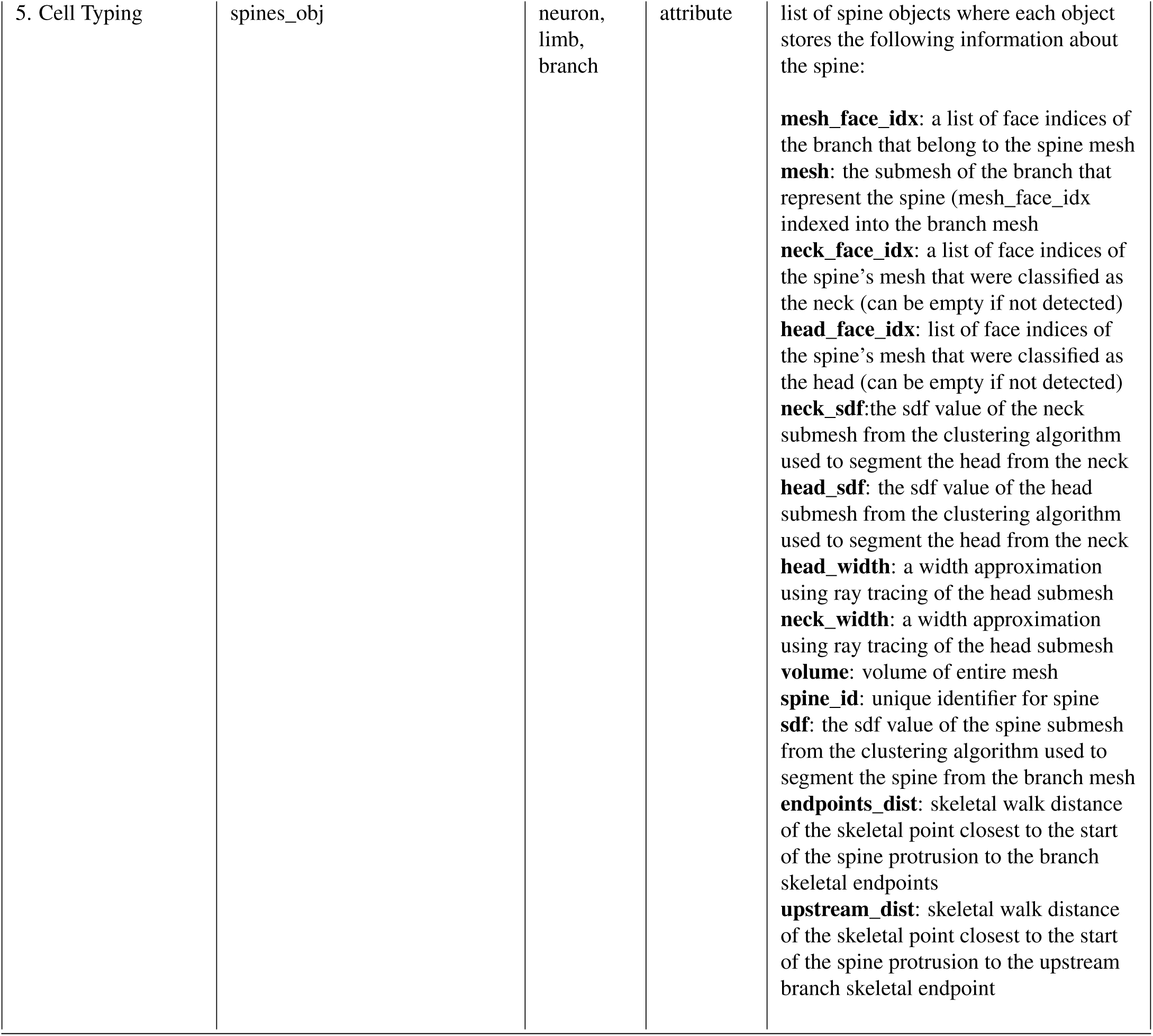
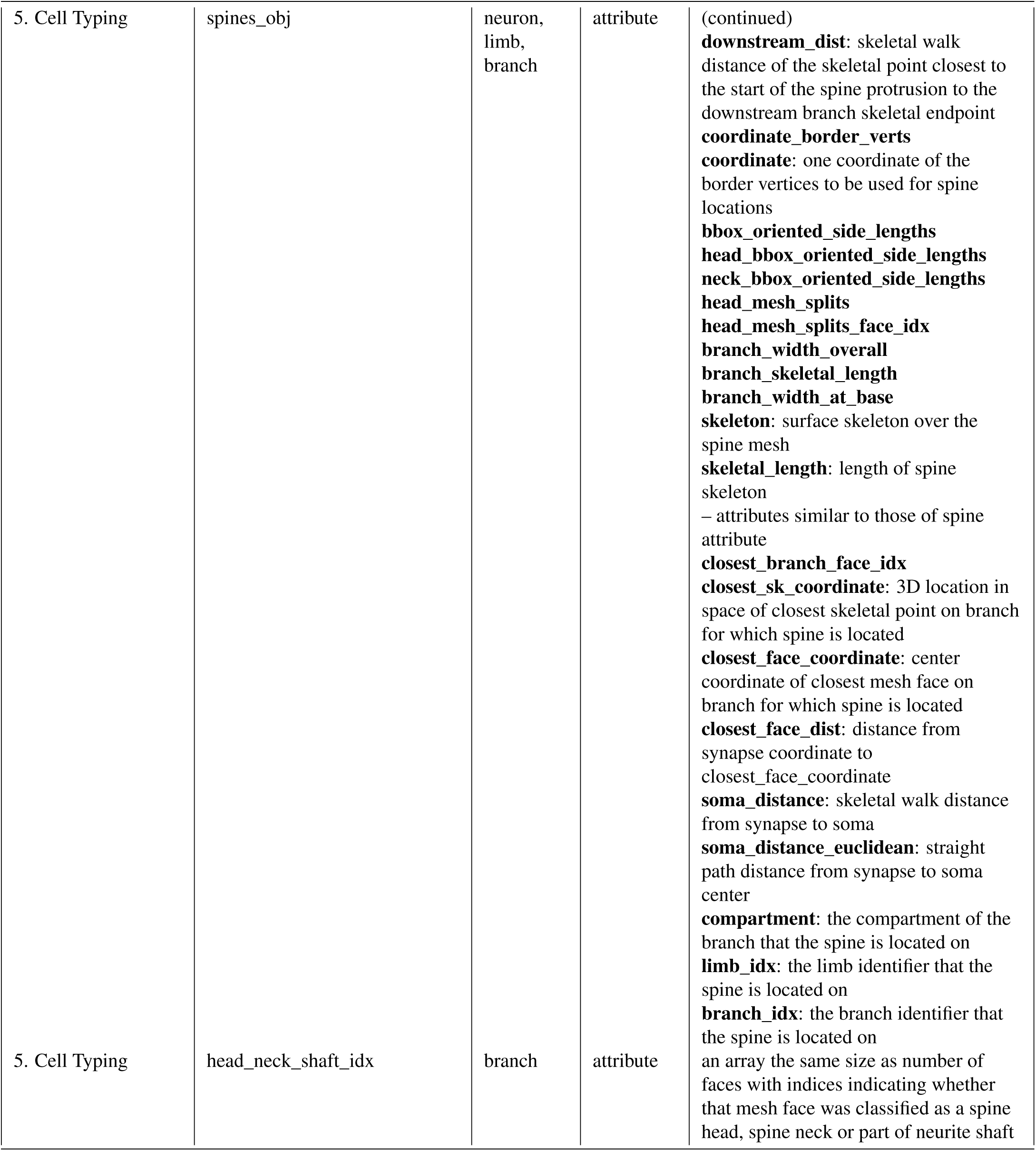
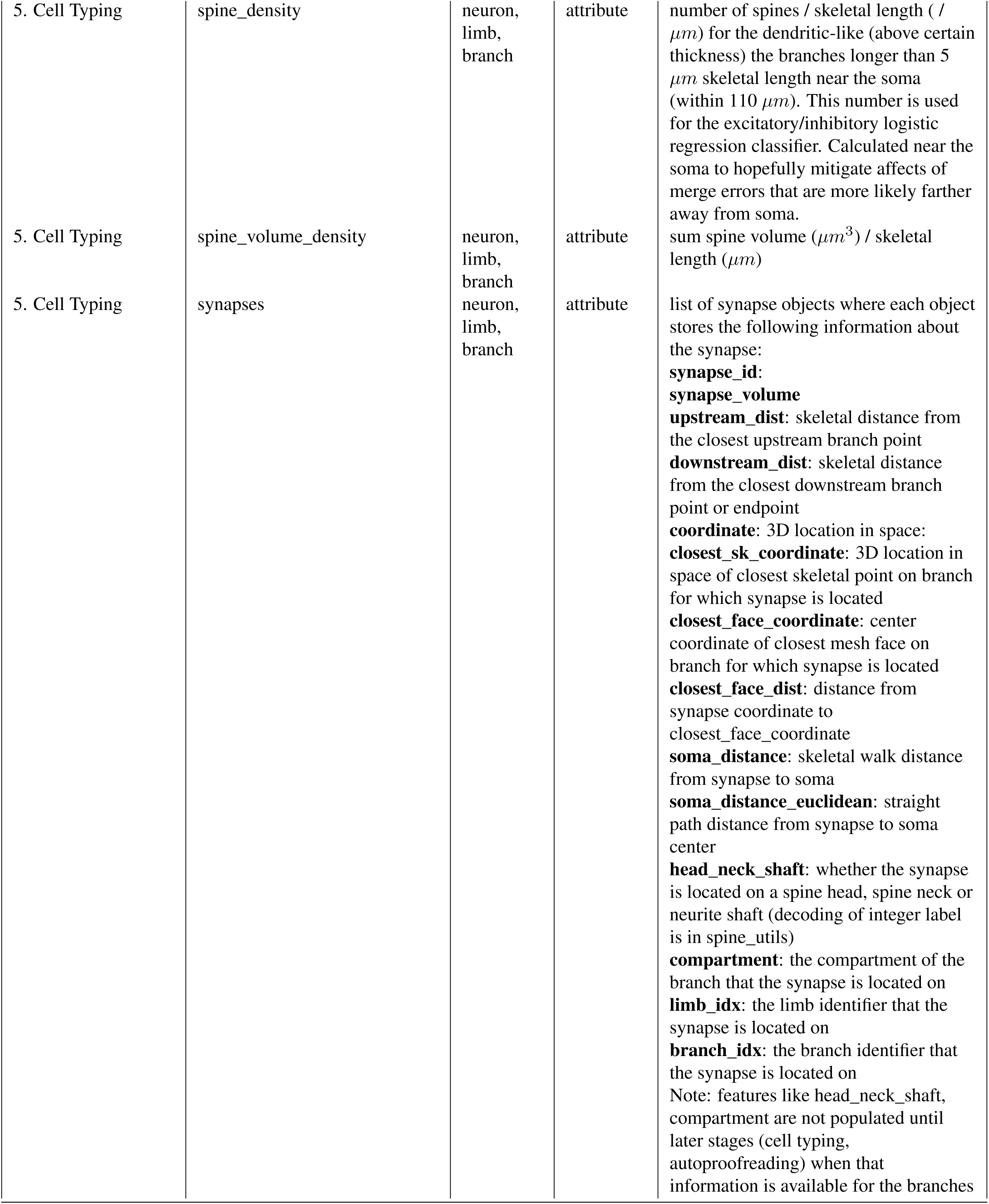
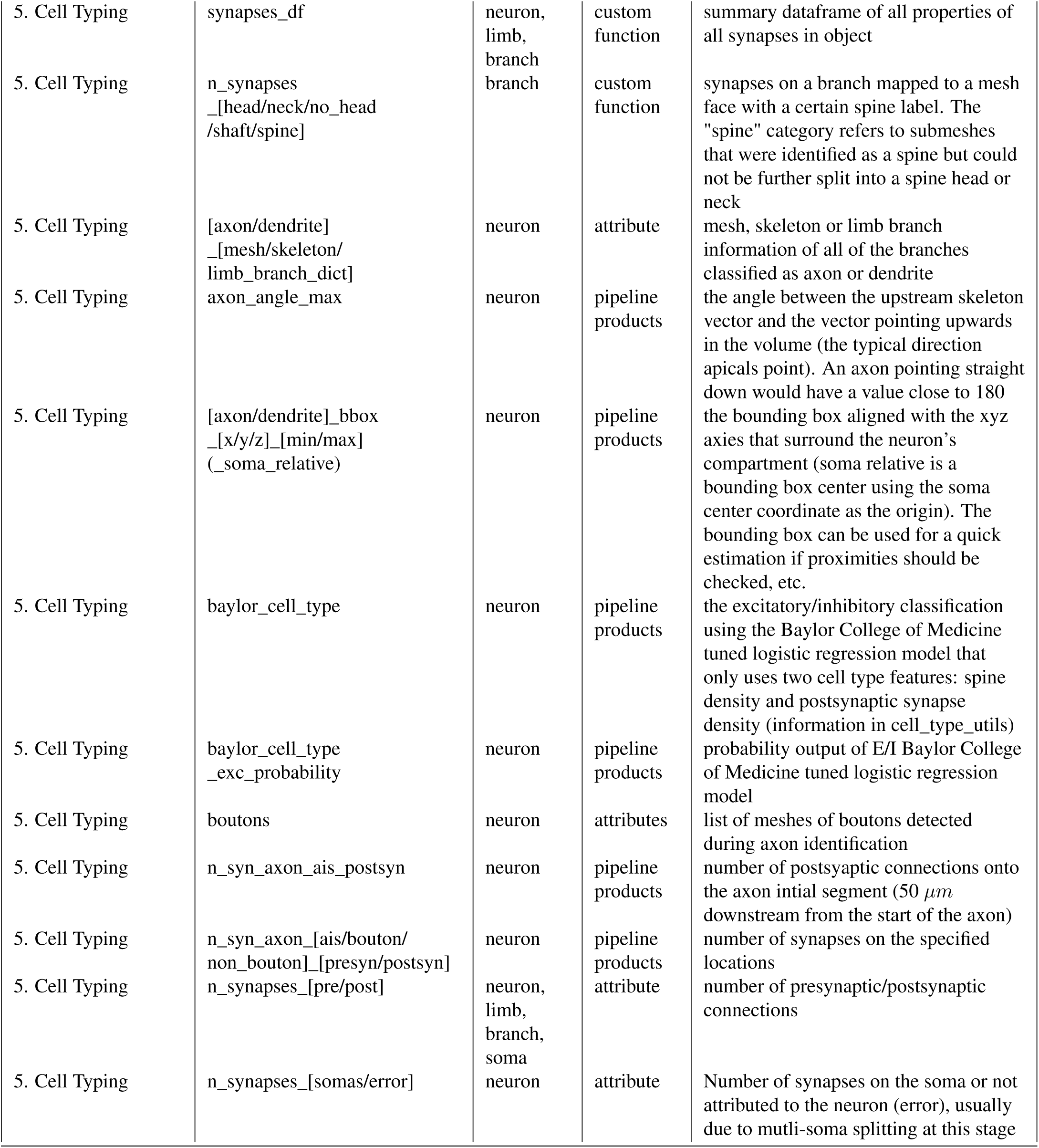
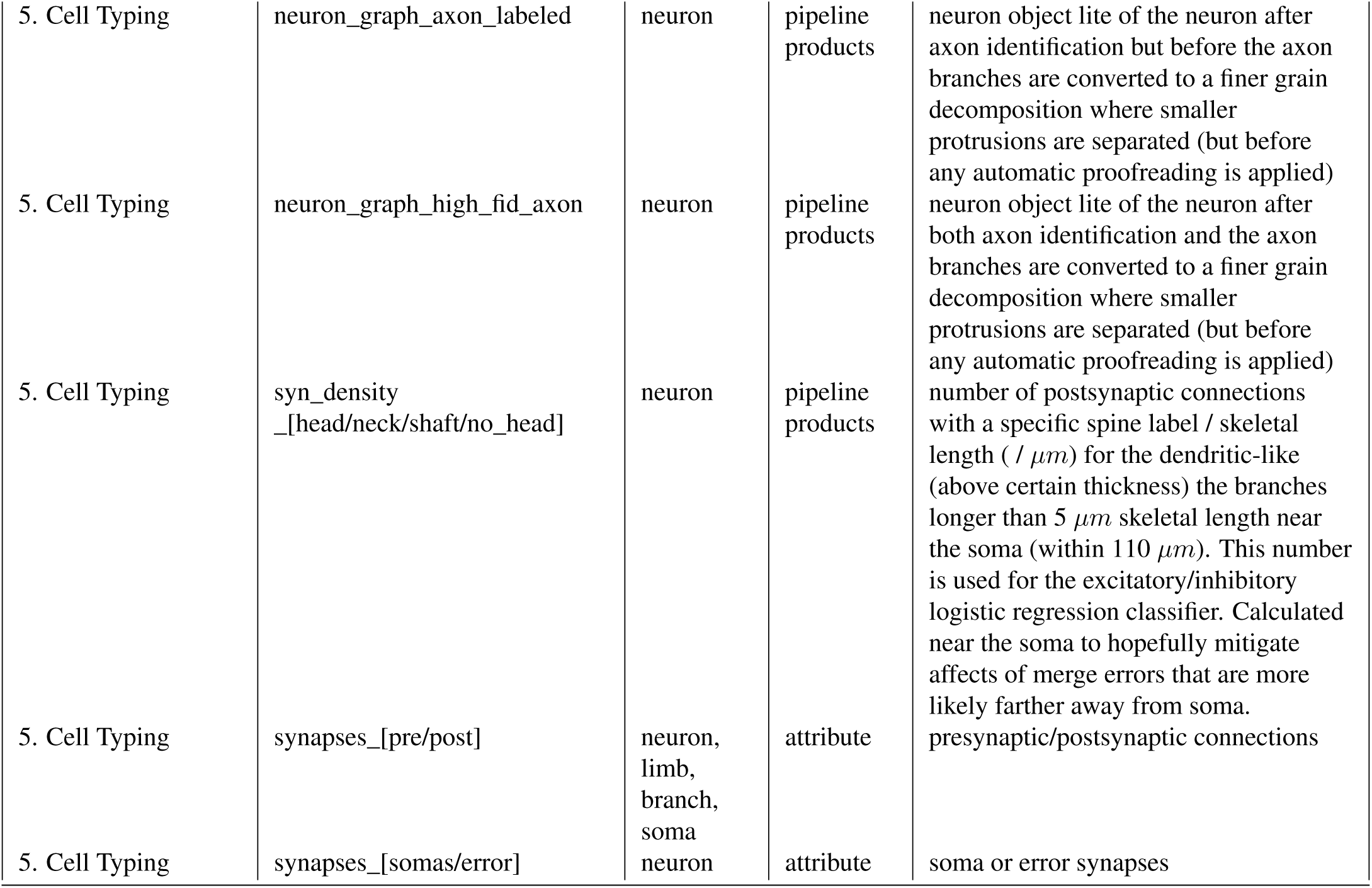
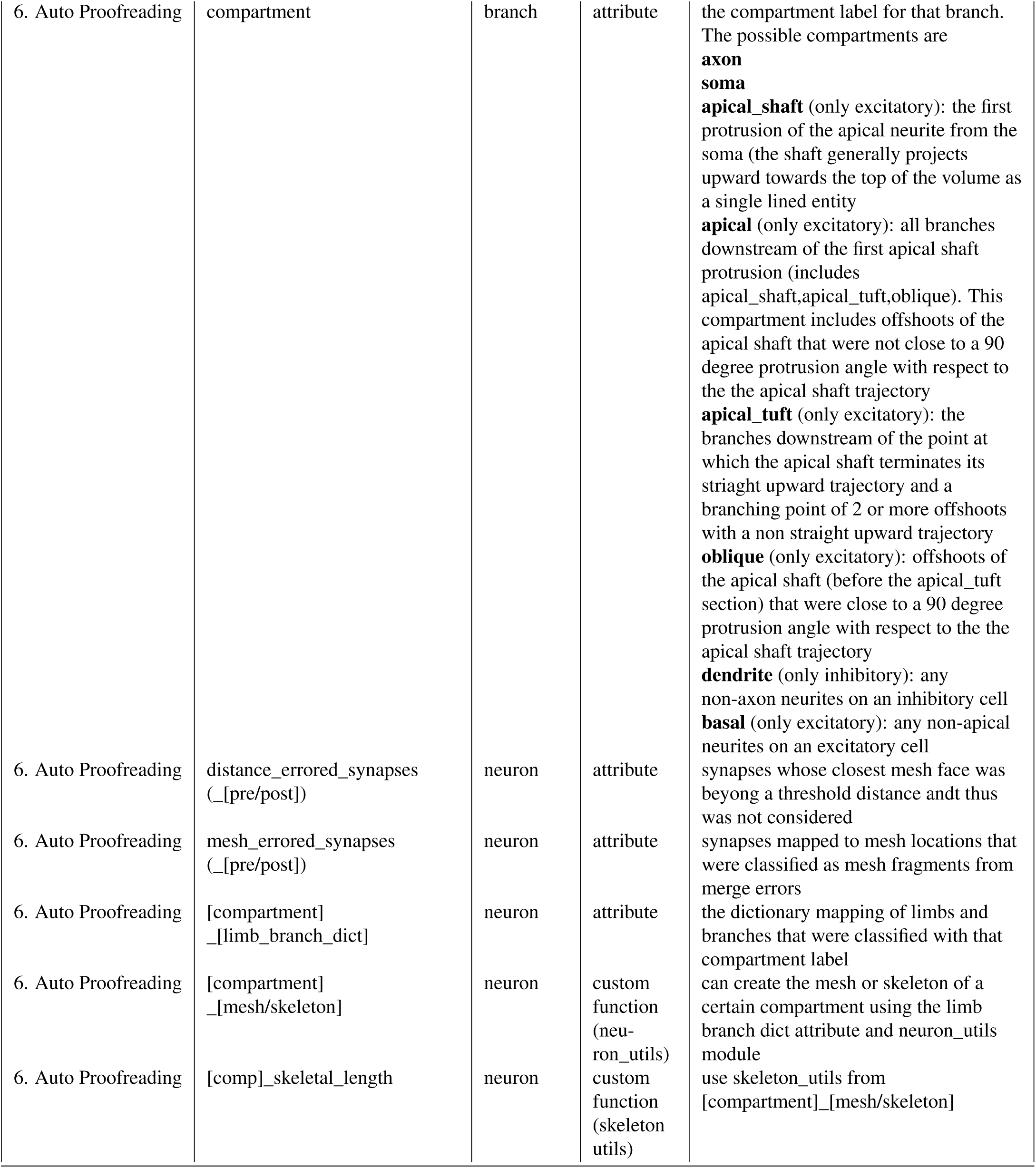
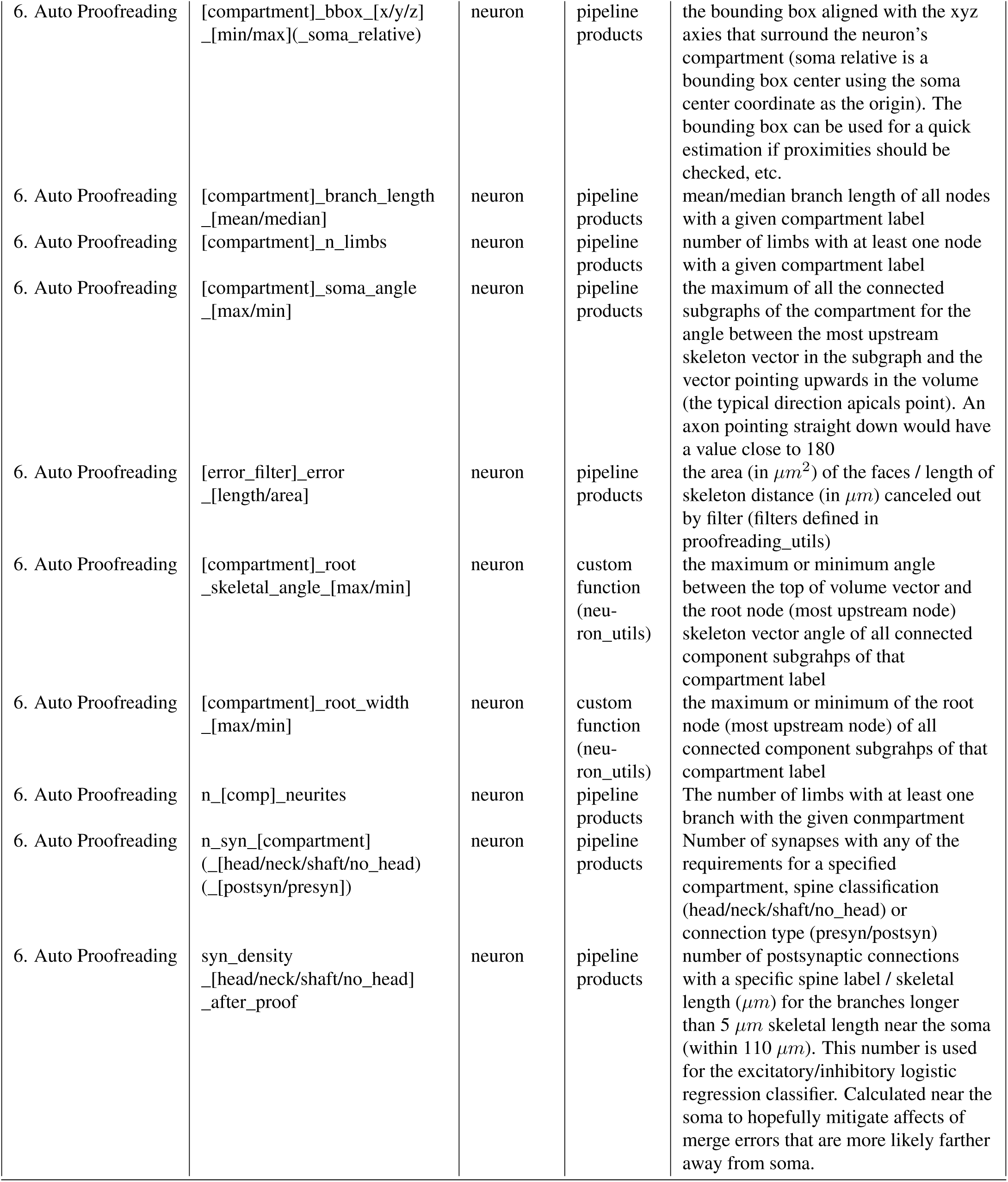
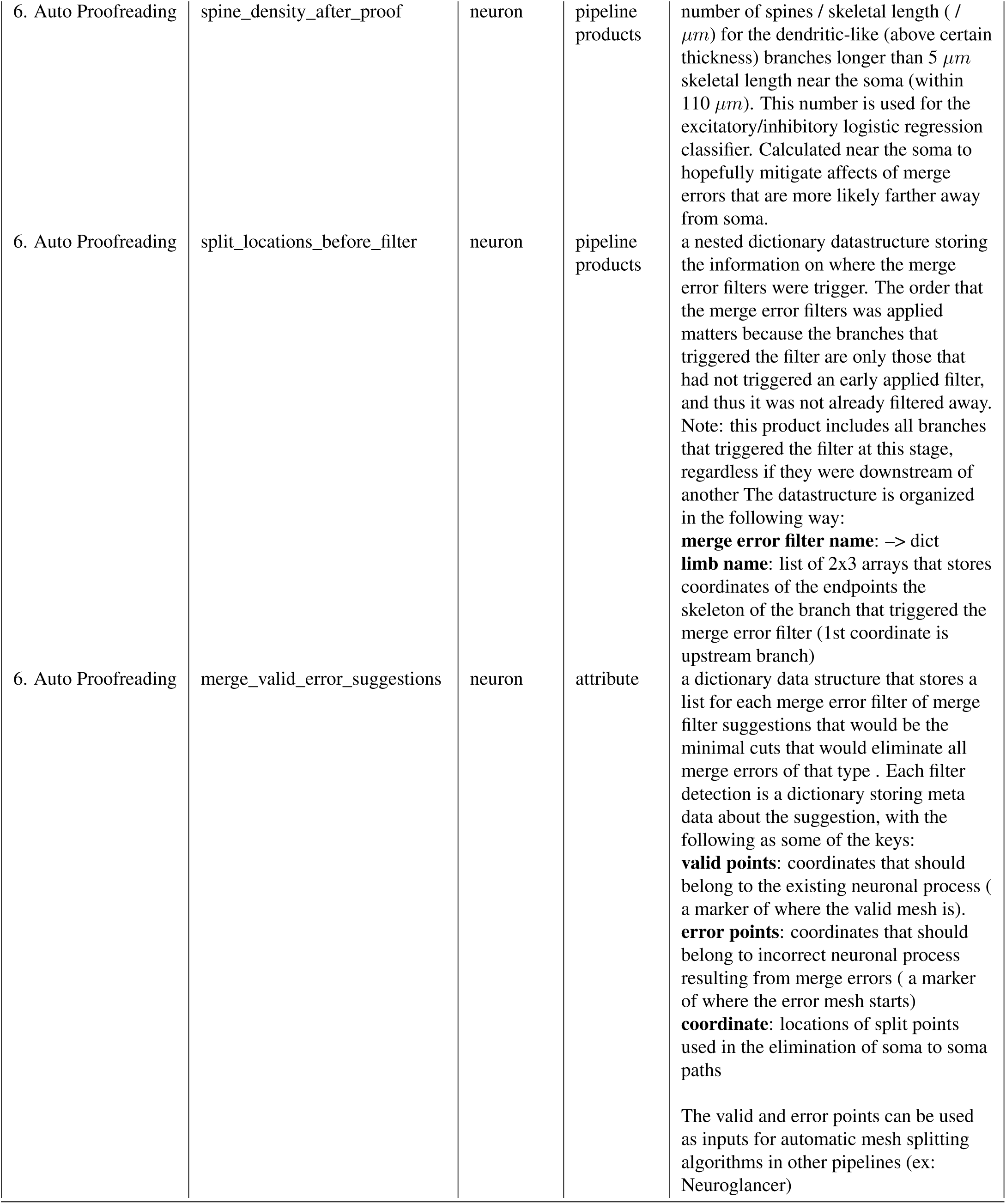
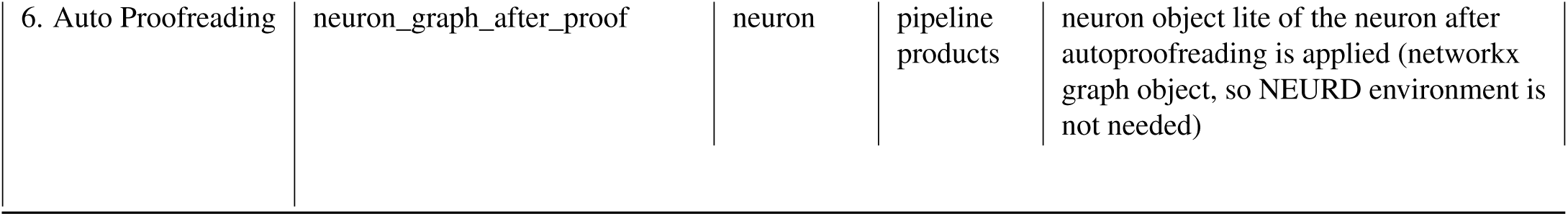
Neuron Feature Documentation.

